# IMAGE: High-powered detection of genetic effects on DNA methylation using integrated methylation QTL mapping and allele-specific analysis

**DOI:** 10.1101/615039

**Authors:** Yue Fan, Tauras P. Vilgalys, Shiquan Sun, Qinke Peng, Jenny Tung, Xiang Zhou

## Abstract

Identifying genetic variants that are associated with methylation variation – an analysis commonly referred to as methylation quantitative trait locus (mQTL) mapping -- is important for understanding the epigenetic mechanisms underlying genotype-trait associations. Here, we develop a statistical method, IMAGE, for mQTL mapping in sequencing-based methylation studies. IMAGE properly accounts for the count nature of bisulfite sequencing data and incorporates allele-specific methylation patterns from heterozygous individuals to enable more powerful mQTL discovery. We compare IMAGE with existing approaches through extensive simulation. We also apply IMAGE to analyze two bisulfite sequencing studies, in which IMAGE identifies more mQTL than existing approaches.

## Introduction

DNA methylation is a stable, covalent modification of cytosine residues that, in vertebrates, typically occurs at CpG dinucleotides. DNA methylation also functions as an important epigenetic regulatory mechanism, with known roles in genomic imprinting, X-inactivation, and suppression of transposable element activity [1, 2]. DNA methylation is thus thought to play a key role in responding to the environment and generating trait variation, including variation in disease susceptibility. In support of this idea, methylation levels have been associated with diabetes [3, 4], autoimmune diseases [5–7], metabolic disorders [8–10], neurological disorders [11, 12], and various forms of cancer [13–17].

Importantly, DNA methylation variation at individual CpG sites often has a strong genetic component [18–29]. Family-based and population-based studies have shown that DNA methylation levels are 34% heritable on average in adipose tissue and are 18-20% heritable on average in whole blood, with heritability estimates reaching as high as 97% [21, 24, 26, 30]. Genetic effects on DNA methylation levels can be explained, at least in part, by *cis*-acting SNPs located close to target CpG sites, where CpG methylation level is associated with the identity of physically linked alleles [23, 31–35]. Indeed, recent methylation quantitative trait loci (mQTL) mapping studies have shown that up to 28% of CpG sites in the human genome are associated with nearby SNPs [23, 26, 31, 32, 36]. Further, *cis*-mQTL often colocalize with disease-associated loci and *cis*-expression QTL (*cis-*eQTL) [26], suggesting that genetic effects on gene expression may be mediated by DNA methylation. Therefore, identifying *cis*-mQTL is an important step towards understanding the genetic basis of gene regulatory variation and, ultimately, organism-level traits.

Most mQTL mapping studies thus far rely on DNA methylation data generated using array-based platforms [36–38]. However, the falling cost of sequencing and the development of high-throughput sequencing-based approaches to measure DNA methylation levels makes mQTL mapping using sequencing data increasingly feasible. Sequencing-based approaches offer several advantages. They can extend the breadth of DNA methylation analysis to the full genome (e.g., via whole genome bisulfite sequencing [39]), increase the flexibility to target specific regions of interest (e.g., via capture methods [40]), improve the representation of genomic regions or regulatory elements that are poorly represented on current array platforms (e.g., via reduced representation bisulfite sequencing [41, 42]), and distinguish 5-hmc modifications from 5-mc modifications (e.g., via TET-assisted pyridine borane sequencing [43] or TAB-seq approaches [44]). Further, unlike arrays, which are largely limited to studies in humans, sequencing-based approaches can be applied to any species [45–48]. Therefore, sequencing-based approaches have become the workhorse of major initiatives like the 1001 Genomes Project in the plant model system *Arabidopsis thaliana* [49, 50]. Importantly, sequencing techniques also facilitate the estimation of allele-specific methylation levels, which should greatly improve the power of mQTL mapping approaches (as allele-specific expression estimates have been shown to do for eQTL mapping: [51, 52]). Early attempts to perform mQTL mapping with bisulfite sequencing data have yielded promising results [35, 49, 53]. However, existing mQTL mapping methods are designed with array data in mind [37, 38]. To maximize power, mQTL mapping using sequencing data requires new statistical methods development that can properly account for two of its distinctive features.

First, methylation data collected in sequencing studies are counts, not continuous representations like those produced by arrays. Specifically, methylation level estimates at a given cytosine base are based on both the total read count at the site and the subset of those reads that are unconverted by sodium bisulfite (or other processes [43]). Previous mQTL studies have dealt with these data by first computing a ratio between the methylated count and the total count, and then treating this ratio as an estimate of the true methylation level [35, 49]. However, the count nature of the raw data means that the mean and variance of the computed ratio are highly interdependent. This relationship is not captured by previously deployed linear regression methods, which likely leads to loss of power. Indeed, similar losses of power are well-documented for differential methylation analysis [40] and differential expression analysis of RNA-seq data [54–57]. To overcome this challenge, statistical methods for sequencing-based differential methylation analysis now adapt over-dispersed count models, including beta-binomial models [58–62] and binomial mixed models [40, 63, 64], to properly model the mean-variance relationship and potential over-dispersion. In differential methylation analysis, these approaches can substantially improve power compared with normalization based approaches [30, 65, 66]. Because mQTL mapping is conceptually similar and can be effectively viewed as genotype-based differential methylation analysis, extending over-dispersed binomial models to mQTL mapping is a promising approach.

Second, sequencing-based techniques are capable of measuring DNA methylation levels in heterozygotes in an allele-specific fashion (i.e., allele-specific methylation, ASM). When ASM estimates support differences in methylation levels between the two alleles carried by heterozygotes, they can be used to increase the power of mapping analysis. Indeed, assuming that additive genetic effects dominate, true *cis*- acting genetic differences in DNA methylation are expected to lead to both (i) differential methylation by genotype across all three genotypes at a biallelic site, and (ii) ASM in heterozygotes. These two types of evidence are only available in sequencing studies, since ASM is not generally detectable when DNA methylation is profiled using arrays. Notably, previous methods for detecting genotype-dependent ASM suggest that it is common across tissue types and species, is more often explained by *cis*-acting variants than *trans*-effects, and is enriched near genes that also display patterns of allele-specific expression [67–75]. Thus, integrating ASM analysis into mQTL mapping analyses should also contribute to understanding the basis of *cis*-regulatory effects on gene expression. There is strong precedent for such a combined strategy in other omics studies. For example, the methods implemented in TreCASE and WASP can integrate allele-specific expression information to greatly enhance the power of eQTL mapping [51, 76–78], and the software RASQUAL integrates allele-specific patterns with individual-level differences to facilitate QTL mapping of chromatin accessibility and ChIP-seq data [79]. However, to our knowledge, no method currently exists for integrating ASM with mQTL mapping in sequencing-based studies of DNA methylation.

Here, we develop a new statistical method for mQTL mapping in bisulfite sequencing studies that both accounts for the count-based nature of the data and takes advantage of ASM analysis to improve power. We refer to our method as IMAGE (Integrative Methylation Association with GEnotypes), which is implemented as an open source R package (www.xzlab.org/software.html). IMAGE jointly accounts for both allele-specific methylation information from heterozygous individuals and non-allele-specific methylation information across all individuals, enabling powerful ASM-assisted mQTL mapping. In addition, IMAGE relies on an over-dispersed binomial mixed model to directly model count data, which naturally accounts for sample non-independence resulting from individual relatedness, population stratification, or batch effects that are commonly observed in sequencing studies [40, 57]. We develop a penalized quasi-likelihood (PQL) approximation-based algorithm [64, 80, 81] to facilitate scalable model inference. We illustrate the effectiveness of IMAGE and compare it with existing approaches in simulations. We also apply IMAGE to map mQTLs in two bisulfite sequencing studies from wild baboons and wild wolves.

## Results

### Method Overview and Simulation Design

IMAGE is described in detail in the Materials and Methods, with additional information provided in the Additional file 1: Supplementary Text. Briefly, IMAGE combines the benefits of both standard mQTL mapping and ASM analysis by jointly modeling non-allele-specific (i.e. per-individual) methylation information across all individuals together with allele-specific methylation information (i.e. per-allele) from heterozygous individuals. This approach enables *cis*-mQTL mapping when the heterozygous SNP and the CpG site of interest are captured either on the same sequencing read or with known phasing information (Fig. 1). By combining both allele-specific and non-allele-specific information, IMAGE improves power over traditional mapping approaches that use non-allele-specific information alone. In addition, IMAGE relies on a binomial mixed model to directly model count data from bisulfite sequencing and naturally accounts for over-dispersion as well as sample non-independence. IMAGE uses a penalized quasi-likelihood based algorithm for scalable inference and is implemented in an open-source R package, freely available at http://www.xzlab.org/software.html.

**Fig. 1:**
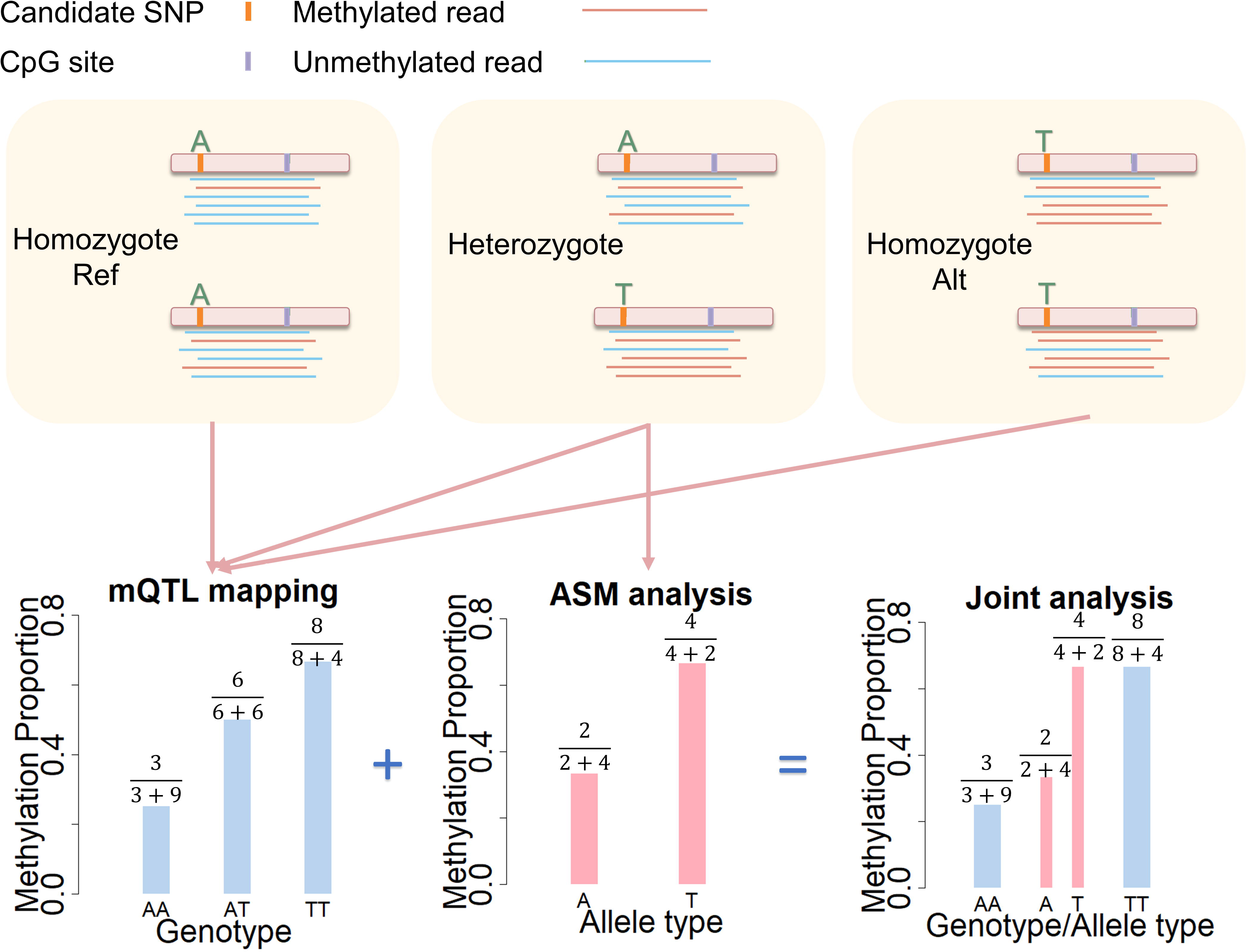
Schematic of ASM-assisted mQTL mapping. The top three panels show bisulfite sequencing data mapped to a CpG site where methylation level is associated with a nearby SNP, in an AA homozygote (left), an AT heterozygote (middle), and a TT homozygote (right). Note that, while illustrated in the panels, the allele-level methylation information in the two homozygotes is not observed. The bottom three panels depict three methods to detect SNP-CpG association: the standard mQTL mapping approach (left) uses non-allele-specific information from all three individuals to detect an association; the standard ASM analysis (middle) uses allele-level information from the heterozygotes only; and the joint analysis approach (right) presented here uses both types of information to achieve a gain in power. mQTL: methylation quantitative trait loci; ASM: allelic specific methylation.

We performed simulations to examine the effectiveness of IMAGE and compare it with other approaches. In each simulation, we started with real genotypes for n =50-150 individuals [82] and examined power and accuracy over a range of parameters: the background heritability *h*^2^; the over-dispersion variance (*σ*^2^; the SNP minor allele frequency *MAF*; the expected per-site total read *TR* across individuals; the average methylation ratio *π*_0_; the SNP effect size *PVE*; the sample size n; and the proportion of total environmental variance that is shared between two alleles ρ (a detailed explanation of these parameters is available in Materials and Methods). In the simulations, we examined the role of each of these eight modeling parameters in determining mQTL mapping power. To do so, we first created a baseline simulation scenario where we set the simulation parameters to typical values inferred from real data [40] (Materials and Methods). Afterwards, we changed one parameter at a time to create different simulation scenarios and examined the influence of each parameter on method performance. In each scenario, we simulated 10,000 SNP-CpG pairs. For 9,000 pairs, the methylation level at the CpG site was independent of the SNP genotype, while for the remaining 1,000 pairs, CpG site methylation was associated with the SNP genotype, such that genotype explained a fixed proportion of methylation levels equivalent to the parameter *PVE*. After simulation, we discarded the methylation measurements for CpG sites on non-informative individuals (i.e., those with total read counts of zero). We then applied IMAGE and five other approaches to analyze each SNP-CpG pair separately.

The five other approaches perform mQTL mapping using different information: (1) IMAGE-I, a special case of IMAGE, which uses only non-allele-specific, individual level information across all individuals; (2) IMAGE-A, another special case of IMAGE, which uses only allele-specific information from heterozygous individuals; (3) MACAU [40, 57], which uses a binomial mixed model to perform mQTL mapping using only non-allele-specific information; (4) GEMMA [83–85], which uses a linear mixed model to perform mQTL mapping using only non-allele-specific information; and (5) BB, which implements a beta-binomial model [40] to perform mQTL mapping using only non-allele-specific information. Note that, with the exception of IMAGE and IMAGE-A, all methods perform mQTL mapping using only non-allele-specific information. In addition, with the sole exception of GEMMA, all methods model counts directly. For GEMMA, we used normalized data in the form of M-values for analysis, following the previous literature [40, 57]. We performed 10 simulation replicates (each consisting of 10,000 SNP-CpG pairs) for each scenario and computed power based on a known false discovery rate (FDR) for each scenario by combining simulation replicates.

## Simulation Results

Overall, the simulation results show that IMAGE outperforms all other methods across all tested parameters (Fig. 2 and Additional file 2: Fig. S1). For example, in the baseline simulation scenario, at an FDR of 0.05, IMAGE reaches a power of 57.15% in a sample size of 100 individuals. IMAGE-I, IMAGE-A, MACAU, GEMMA and BB reach a power of 7.55%, 10.27%, 7.49%, 2.25% and 6.79%, respectively. The ranking of different methods is not sensitive to different FDR cutoffs. For example, at an FDR of 0.1, the power of IMAGE is 68.78%; while the power of IMAGE-I, IMAGE-A, MACAU, GEMMA and BB is 14.98%, 24.35%, 13.64%, 2.84% and 15.03%, respectively. The superior performance of IMAGE suggests that incorporating ASM information into mQTL mapping can greatly enhance power.

**Fig. 2:**
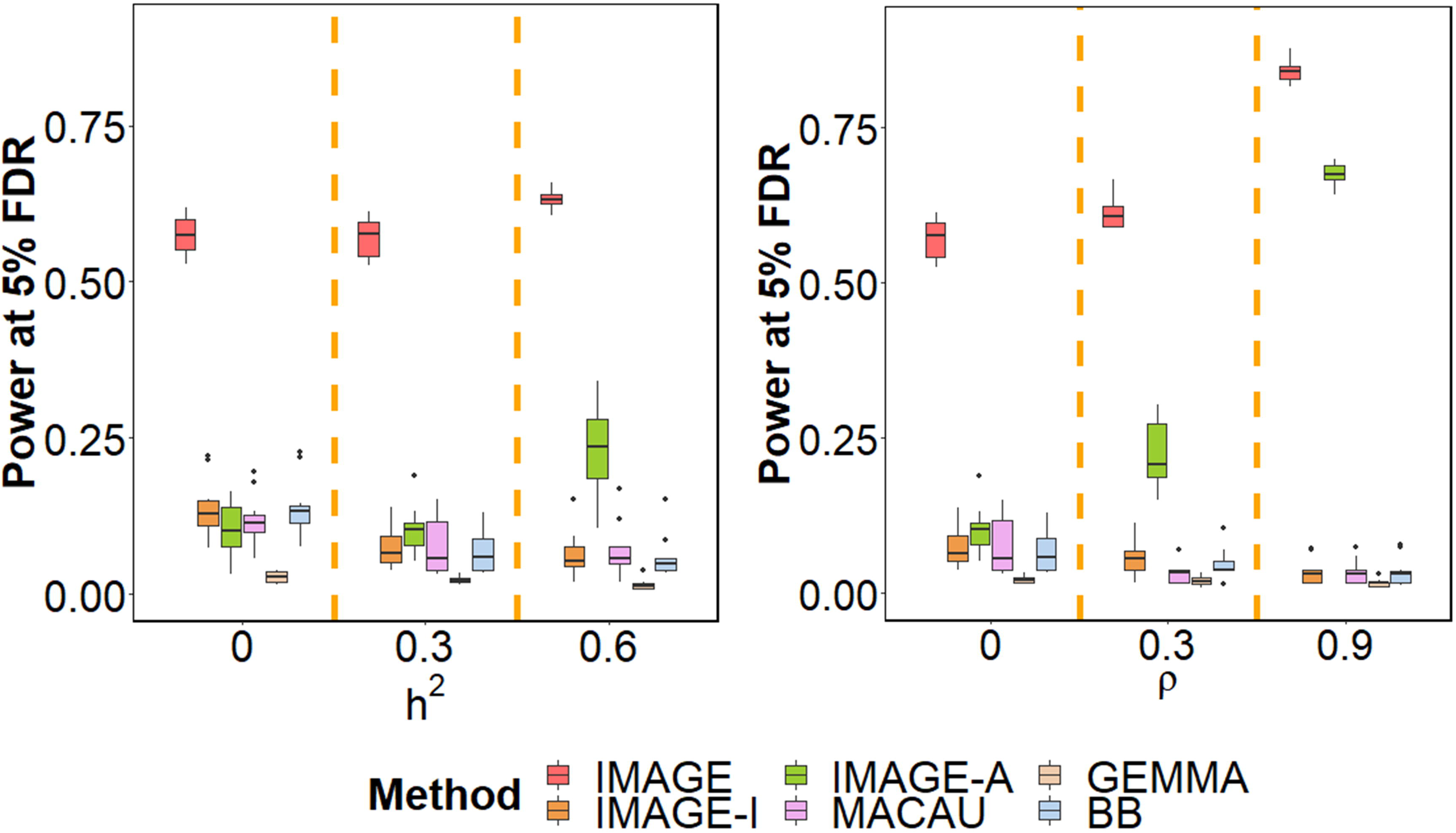
IMAGE achieves higher power to detect mQTL across various simulation settings. Power is measured by number of true mQTL detected at a false discovery rate (FDR) of 0.05. Each simulation setting is based on 10 simulation replicates, each including 10,000 simulated SNP-CpG pairs, 10% of which represent true mQTL. (A) We vary *h*^2^, the background heritability, to be either 0, 0.3, or 0.6, while maintaining other parameters at baseline. (B) We vary ρ, the proportion of common environmental variance, to be either 0, 0.3 or 0.9, while maintaining other parameters at baseline. The middle panel in (A) and the left panel in (B) correspond to the baseline simulation setting. Increasing both *h*^2^ and ρ, which capture genetic and common environmental background effects, respectively, results in increased power for methods that use ASM information (IMAGE and IMAGE-A), but losses in power for methods that do not use ASM information (IMAGE-I, MACAU, GEMMA, BB). FDR: false discovery rate.

Among the eight parameters we examined, six have similar effects on power across IMAGE and the five other models we compared. For example, the power of all methods increases with larger sample size *n* (Additional file 2: Fig. S1A), larger genetic effect size *PVE* (Additional file 2: Fig. S1B), larger minor allele frequency *MAF* (Additional file 2: Fig. S1C), larger read depth TR (Additional file 2: Fig. S1D), and larger over-dispersion variance (*σ*^2^, which implicitly increases the genetic effect size *PVE* (Additional file 2: Fig. S1E). In addition, the power of all methods is highest for CpG sites with intermediate methylation level *π*_0_, but reduced for both hypomethylated and hypermethylated sites (Additional file 2: Fig. S1F). The power dependence on *π*_0_ is presumably because higher methylation variance in the middle range of *π*_0_ leads to higher power.

Careful examination of the relative performance of different methods in different scenarios yields additional insights. First, among the mQTL mapping methods, we found that count-based approaches (IMAGE-I, MACAU, BB) often outperform a normalized data-based approach (GEMMA). Such performance differences become more apparent when sample size *n* is small (Additional file 2: Fig. S1A), methylation level *π*_0_ is either low or high (Additional file 2: Fig. S1F), or mean per-site read depth *TR* is low (Additional file 2: Fig. S1D). For example, when the mean total read *TR* = 10, the power of IMAGE-I, MACAU and BB is 5.8%, 4.56% and 5.33%, respectively (*n*=100); while the power of GEMMA is only 1.01%. When TR increases to 30, the power of IMAGE-I, MACAU and BB becomes 15.25%, 15.32% and 14.55%, respectively, while the power of GEMMA remains low, at 6.14%. The superior performance of count-based methods is consistent with previous observations [40, 57], suggesting that modeling sequencing data in the original count form has added benefits for mQTL mapping. For DNA methylation levels, this advantage may arise in part because uncertainty in DNA methylation level estimates is more accurately modeled in the count data than in normalized ratios. For example, a methylation level of one (completely hypermethylated) is strongly supported for a site-sample combination where read depth is very high, but weakly supported for combinations where read depth is low. The count-based methods effectively capture this distinction, which is lost in conversion to a single ratio.

Second, ASM-based approaches (IMAGE and IMAGE-A) often outperform mQTL mapping approaches that only use non-allele-specific data. This result holds even for IMAGE-A, even though it only models data for heterozygotes at nearby SNPs (and hence, uses only a subset of the data: 42% of the full set of simulated individuals on average). The generally higher power of ASM analysis likely stems from the fact that ASM methods control for both environmental and *trans*-acting genetic background effects (for each heterozygote, both alleles reside in the same individual, providing a natural internal control). Our simulations suggest that there are two important parameters that influence the relative power of ASM analysis and mQTL mapping. The first important parameter is background heritability, *h*^2^. Increased background heritability can reduce the performance of mQTL mapping methods, as increased confounding from polygenic effects of other SNPs likely increases the difficulty of identifying individual SNP associations [40, 57]. For example, when *h*^2^ = 0, the power of IMAGE-I, MACAU, GEMMA and BB is 13.57%, 11.62%, 2.69% and 13.88%, respectively. When *h*^2^ increases to 0.6, however, the power of IMAGE-I, MACAU, GEMMA and BB reduces to 6.48%, 7.05%, 1.50% and 5.92%, respectively. In contrast, ASM analysis relies on a model that explicitly accounts for the heritable component that arises from genetic background effects, and thus achieves relatively stable performance. For example, when *h*^2^ = 0, the power of IMAGE and IMAGE-A is 57.48% and 10.30%, respectively. When *h*^2^ increases to 0.6, the power of IMAGE and IMAGE-A actually increases, to 63.07% and 23.09%, respectively. This observation is consistent with the fact that the two alleles modeled in ASM, for each individual, share an identical genetic background that becomes easier to control for as its contribution to DNA methylation increases (i.e., as **h*^2^* increases). Thus, IMAGE-I outperforms IMAGE-A when background heritability is zero (*h*^2^ = 0), but performs worse when background heritability is moderate or high (*h*^2^ = 0.3 or 0.6; Fig. 2A).

The second important parameter is the ratio parameter ρ, which represents the relative contribution of shared/common environmental effects (i.e., the “trans” acting environment). also influences the relative power of ASM vs mQTL. For mQTL methods, increasing ρ necessarily increases the contribution of common environmental noise shared between the two alleles. Common environmental noise is not explicitly accounted for by mQTL models, thus leading to a reduction in power. For example, when ρ = 0, IMAGE-I, MACAU, GEMMA and BB detect 7.55%, 7.49%, 2.25% and 6.79% of true effects, respectively. When ρ increases to 0.9, the power of IMAGE-I, MACAU, GEMMA and BB reduces to 3.50%, 3.44%, 1.67% and 3.57%, respectively. In contrast, ASM analysis explicitly accounts for both common and independent environmental background effects, again because it measures DNA methylation in the two alleles in the same individual. ASM methods thus achieve better, not worse, performance with higher values of ρ. For example, when ρ = 0, the power of IMAGE and IMAGE-A is 57.15% and 10.27%, respectively. When ρ increases to 0.9, the power of IMAGE and IMAGE-A becomes 84.15% and 67.55%, respectively. Consequently, while mQTL methods have similar power as ASM when ρ is small, ASM can outperform mQTL when ρ is large (Fig. 2B).

In addition, we note that IMAGE can estimate FDR reasonably accurately by constructing an empirical null via permutations. In particular, IMAGE produces either calibrated or slightly conservative FDR estimates regardless of the values of *h*^2^ (Additional file 2: Fig. S2A), ρ (Additional file 2: Fig. S2B), *n* (Additional file 2: Fig. S2C), genetic effect size *PVE* (Additional file 2: Fig. S2D), *MAF* (Additional file 2: Fig. S2E), average read counts per site *TR* (Additional file 2: Fig. S2F), over-dispersion variance (*σ*^2^ (Additional file 2: Fig. S2G), or average methylation ratio *π*_0_ (Additional file 2: Fig. S2H).

Finally, we note that while we set PVE=0.10 and *h*^2^ = 0.30 in the baseline simulations to capture the realistic effect sizes and background heritability across all SNP-CpG pairs genome-wide, reasonable data filtering decisions will often increase mean PVE and *h*^2^ among SNP-CpG pairs tested in real data applications. For example, in the wolf and baboon data sets analyzed below the median PVE was approximately 0.15 and the median *h*^2^ estimate was near 0.5. For direct comparability, we therefore also created a simulation scenario in which we set PVE to 0.15 and *h*^2^ to 0.50 (Additional file 2: Fig. S1G). Notably, the relative power of different methods in this setting largely recapitulates our observations in the real data applications (see below).

### mQTL mapping in wild baboons

We applied our method to analyze a reduced representation bisulfite sequencing data collected on 67 baboons from the Amboseli ecosystem of Kenya [40, 45]. Detailed data description and processing steps are provided in Materials and Methods, with an illustrative processing diagram showing in Additional file 2: Fig. S3. Briefly, we extracted 49,196 SNP-CpG pairs from the bisulfite sequencing data, which consists of 13,753 unique SNPs and 45,210 unique CpG sites. We applied IMAGE together with the other five approaches described above to analyze each SNP-CpG pair individually. We performed permutations to estimate FDR for each method and we report results based on a fixed FDR cutoff.

Consistent with our simulations, our method achieves higher power compared with other methods in the baboon data set (Fig. 3A). For example, at an empirical FDR of 5%, IMAGE detected 7,043 associated SNP-CpG pairs, which is 45% more than that detected by the next best method (IMAGE-A, which detected 4,855 pairs at a 5% FDR). IMAGE-I, MACAU, GEMMA, and BB detected 3,585, 3,024, 2,629 and 3,259 pairs, respectively. Also consistent with the simulations, the higher power of IMAGE compared to other methods is robust with respect to different FDR cutoffs (Fig. 3A). We illustrate a few example sites that were only detected by IMAGE in Additional file 2: Fig. S4. For these sites, methylation levels measured in the heterozygotes are noisy and often indistinguishable from at least one type of homozygote (often because total read counts are unevenly distributed across alleles). However, by separating methylation levels in heterozygotes into the contribution from each individual allele and modeling ASM information together with non-allele-specific information, IMAGE remains capable of identifying mQTLs in these sites. In addition, consistent with simulations, we also observed that our method could detect more associated SNP-CpG pairs with increasing MAF (Additional file 2: Fig. S5A), increasing read depth TR (Additional file 2: Fig. S5B), increasing sample size (Additional file 2: Fig. S5C), or at intermediate methylation levels (Additional file 2: Fig. S5D).

**Fig. 3:**
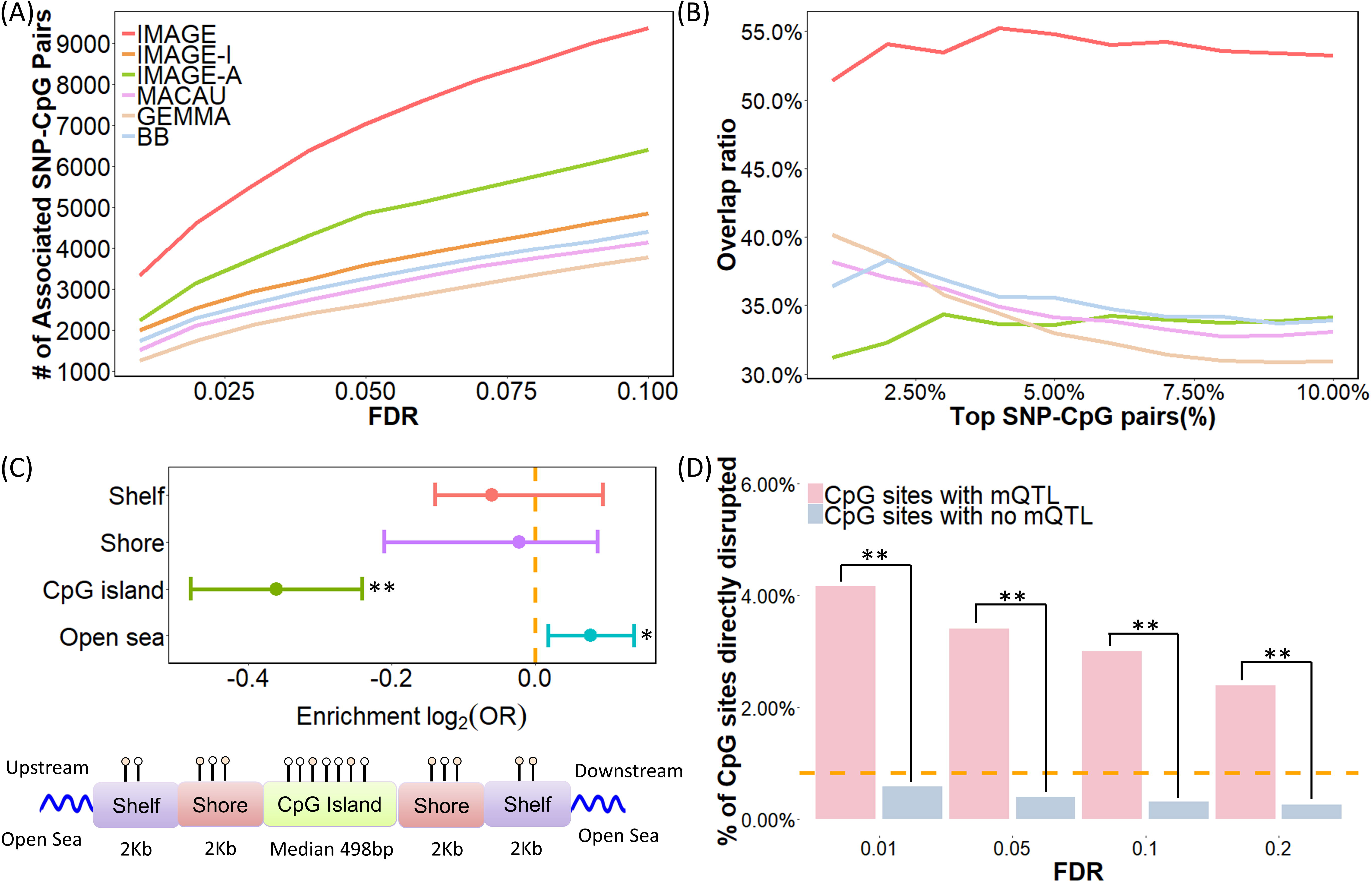
mQTL mapping results in the baboon RRBS data. (A) IMAGE identified more mQTL than the other five methods across a range of empirical FDR thresholds. (B) IMAGE identifies more consistent associations than the other methods in the subset analysis. Here, we randomly split individuals into two approximately equal sized subsets and analyzed the two subsets separately using each method. We then counted the number of overlapping mQTL identified in both subsets. The overlap ratio (y-axis) is plotted against the percentage of top mQTL ranked by statistical evidence for a SNP-CpG methylation association in each method (x-axis). (C) Upper panel: log2 odds ratio of detecting associated SNP-CpG pairs, together with the 95% CI, is computed for CpG sites residing in different annotated genomic regions. CpG sites with IMAGE-identified mQTL are enriched in open sea regions (p-value=0.0106) and depleted in CpG islands (p-value = 1.056 × 10^−9^). Bottom panel: all analyzed CpG sites have been annotated to genomic regions based on their relation to the nearest CpG island. CpG islands were annotated based on UCSC Genome Browser (average length = 672 bp in the data; min = 201 bp; max = 15,960 bp). Shore is the flanking region of CpG islands covering 0-2000bp distant from the CpG island. Shelf is the regions flanking island shores covering 2000-4000 bp distant from the CpG island. (D) A higher percentage of CpG sites are directly disrupted by the SNP in mQTL pairs compared to by chance alone (horizontal dashed line), and more so than in non-mQTL pairs (p-value <2.2 × 10^−16^). Such enrichment decays with increased FDR thresholds. *P<0.05 **P<0.01.

To validate the mQTLs we identified, we randomly split the sample into two approximately equal sized subsets (one with 34 individuals and the other with 33 individuals) and examined the consistency of the SNP-CpG pairs detected in the two subsets. We removed IMAGE-A from this analysis as it requires at least five heterozygous individuals, which is no longer satisfied for many SNP-CpG pairs in each of the two subsets. For the remaining methods, we found that IMAGE detects more consistent SNP-CpG pairs between the two subsets than the other approaches (Fig. 3B). For example, among the top 5% (*n*=2,511) associated SNP-CpG pairs based on IMAGE, 53.8% of them were identified in both subsets. In contrast, among the top 5% (n=2,511) associated SNP-CpG pairs based on IMAGE-I, MACAU, GEMMA and BB, 35.84%, 35.12%, 33.92% and 37.64% overlapped between the two subsets. The greater consistency of results from IMAGE thus provides convergent support for its increased power.

Next, we assessed the set of detected SNP-CpG associations by performing functional enrichment analysis to compare our findings against published results (Fig. 3C). Here, we refer to the CpG sites with associated mQTL as mCpG sites. We examined whether the set of mCpG sites were enriched in CpG islands, CpG island shores, CpG island shelves, or in genomic “open sea”. To do so, we obtained functional genomic annotation information from UCSC Genome Browser for the baboon genome, *Panu2.*0 and relied on the same criterion as [86] to annotate genomic regions (details in Materials and Methods). For each annotated category, we then computed the proportion of mCpG sites in the annotated regions and contrasted it to the proportion of non-mCpG sites analyzed in our original mQTL mapping analysis. We found that mCpG sites are significantly enriched in open seas compared to non-mCpG sites (69.74% vs 66.08%; Fisher’s exact test, *p-value* = 0.0106) but underrepresented in CpG islands (11.16% vs 14.33%; *p*-value =1.056 × 10^−9^). The results are consistent with previous observations [87, 88], partly because CpG islands are often enriched in evolutionarily conserved promoter regions [89–91] that harbor fewer regulatory genetic variants, and partly because power to detect mQTL is lower in hypomethylated regions [92]. The results are qualitatively consistent across sites with different mean CpG methylation levels, although do not reach statistical significance in all bins likely due to the smaller number of sites and the resulting lower power in each bin (Additional file 2: Fig. S6). Importantly, despite the higher number of mCpG sites detected by IMAGE, the evidence for both enrichment in open sea and underrepresentation in CpG islands is also stronger in the IMAGE analysis than for other methods (Additional file 3: Table S1).

Finally, we counted the percentage of SNP-CpG pairs for which the SNP directly resides in the CpG sequence, abolishing the CpG site and therefore resulting in an entirely unmethylated alternate allele [69, 93]. These sites, by definition, should exhibit mQTL and ASM. 403 CpG sites in our data set were disrupted by SNPs, and 59.6% of them (*n*=240) were indeed identified as significant mCpG sites. For 95.70% of those we did not detect (*n*=156), the non-disrupted CpG was also hypomethylated in our sample (<10% methylation level), which would make it impossible to detect an mQTL (i.e., because both disrupted and non-disrupted alleles are hypomethylated). CpG sites disrupted by SNPs accounted for 3.72% of significant mCpG sites (compared to the 0.89% expected by chance), but only 0.43% of non-mCpG sites, in support of the accuracy of our mQTL mapping approach (Fisher’s exact test p-value <2.2 × 10^−16^). In addition, as expected, the percentage of significant mCpG sites accounted for by CpG sites disrupted by SNPs gradually decreases with less stringent FDR cutoffs (Fig. 3D). Importantly, IMAGE also outperforms the other five methods on this metric (Additional file 3: Table S2).

### mQTL analysis in wild wolves

Finally, we applied IMAGE to analyze a second RRBS data set collected on 63 grey wolves from Yellowstone National Park [46, 94]. We applied the same data processing procedure described above for baboons, followed by mQTL mapping. In total, we extracted 279,223 SNP-CpG pairs from the bisulfite sequencing data, which consists of 77,039 unique SNPs and 242,784 unique CpG sites. IMAGE again achieved higher power compared with the other methods (Fig. 4A). At an empirical FDR of 5%, IMAGE detected 34,779 significantly associated SNP-CpG pairs, which is 50% more than that detected by the next best method (IMAGE-A), and 262% more than the other four methods (Fig. 4A and Additional file 2: Fig. S7). As in the baboons, subset analysis confirmed that IMAGE detects more consistent SNP-CpG pairs than the other approaches (Fig. 4B). For example, among the top 5% (*n*=14,091) associated SNP-CpG pairs based on IMAGE analysis, 53.8% of them are consistent between the two subsets, compared to 20.5 – 30.7% for the other four methods tested. Consistent with results from simulations and the baboon data, we also observed that our method could detect more associated SNP-CpG pairs with intermediate methylation levels, increasing MAF, increasing read depth, and increasing sample size (Additional file 2: Fig. S5).

**Fig. 4:**
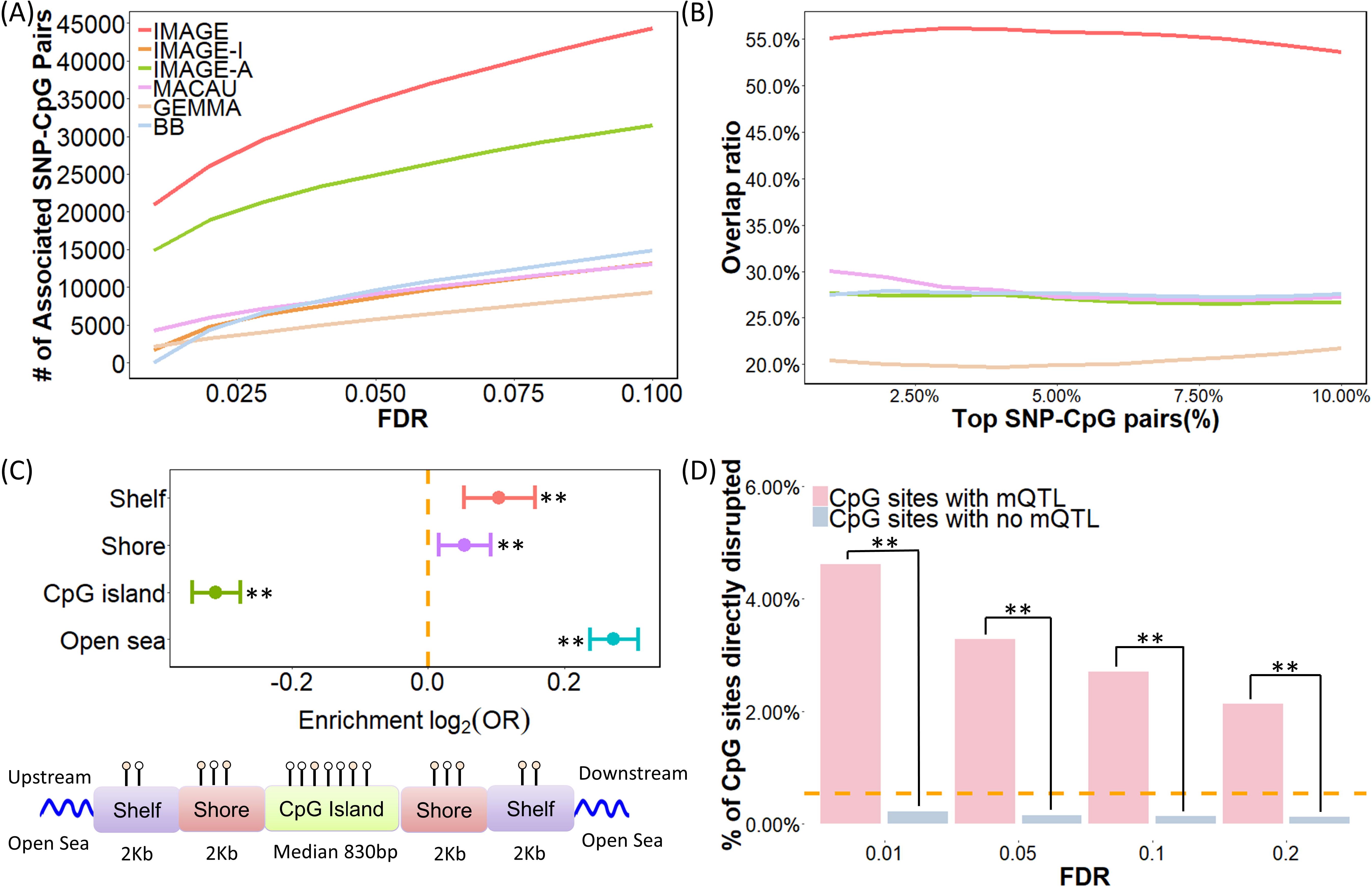
mQTL mapping results in the wolf RRBS data. Methods for analysis include: IMAGE (red), IMAGE-I (orange), IMAGE-A (green), MACAU (pink), GEMMA (brown), BB (blue). (A) IMAGE identified more associated SNP-CpG pairs than the other five methods across a range of empirical FDRs constructed by permutation. (B) IMAGE identifies more consistent associations than the other methods in the subset analysis. Here, we randomly split individuals into two approximately equal sized subsets and applied methods to analyze the two subsets separately. We count the number of overlapping associations between the top SNP-CpG pairs in the two subsets. The overlap ratio (y-axis) is plotted against the percentage of top SNP-CpG pairs (x-axis). (C) Upper panel: log2 odds ratio of detecting associated SNP-CpG pairs, together with the 95% CI, is computed for CpG sites residing in different annotated genomic regions. CpG sites associated with SNPs identified by IMAGE are enriched in open sea regions (p-value <2.2 × 10^−16^) and depleted in CpG island regions (p-value <2.2 × 10^−16^). Shores are defined as the 2000 bp regions flanking CpG islands; shelves are defined as the 2000 bp regions flanking the island shores (2000-4000 bp from CpG islands). Bottom panel: all analyzed CpG sites have been annotated to genomic regions based on their relation to the nearest CpG island. CpG islands were annotated based on UCSC Genome Browser (average length = 830 bp in the data; min = 201 bp; max = 322,257 bp). Shore is the flanking region of CpG islands covering 0-2000bp distant from the CpG island. Shelf is the regions flanking island shores covering 2000-4000 bp distant from the CpG island. (D) A higher percentage of CpG sites are directly disrupted by the SNP in the mQTL pairs compared to by chance alone (horizontal dashed line), and more so than in non-mQTL pairs (p-value <2.2 × 10^−16^). Such enrichment decays with increased FDR thresholds. *P<0.05. **P<0.01.

Finally, consistent with the baboon results, mCpG sites in the wolves were significantly enriched in open sea compared to non-mCpG sites (31.77% vs 26.31%; *p*-value < 2.2 × 10^−16^) and were underrepresented in CpG islands (30.17% vs 37.43%; *p*-value <2.2 × 10^−16^) (Fig. 4C). In the wolves, we also observed significant (albeit much weaker) enrichment of mCpG sites in shelf regions (12.49% vs 11.63%; *p*-value =9.001 × 10^−5^) and shore regions (25.57% vs 24.64%; *p*-value =5.890 × 10^−3^). The higher frequency of mCpG sites in CpG island shelves and shores is consistent with previous studies [87, 88] and likely reflects greater power to detect enrichment in the wolf data set, which yields a larger number of analyzable SNP-CpG pairs than in the baboons (m = 242,784 in wolf vs m = 45,210 in baboon). The enrichment in open sea and underrepresentation of mCpG sites in CpG islands are robust regardless of whether we stratify sites based on mean methylation levels, although the shelf/shore results are noisier (Additional file 2: Fig. S8). Again, we found that enrichment results were stronger in the IMAGE analysis than when using other methods (Additional file 3: Table S3); and that mCpG sites were more likely to be disrupted by their associated SNPs than non-mCpG sites (3.66% versus 0.18%; *p*-value <2.2 × 10^−16^) (Fig. 4D; see also Additional file 3: Table S4).

## Discussion

Here, we present IMAGE, a new statistical method with a scalable computational algorithm, for mQTL mapping in bisulfite sequencing studies. IMAGE relies on a binomial mixed model to account for the count nature of over-dispersed bisulfite sequencing data, models multiple sources of methylation level variance, and incorporates allelic-specific methylation patterns from heterozygous individuals into mQTL mapping. Both simulations and two real data sets support its increased power over other commonly used methods.

A key feature of our method is its ability to incorporate allele-specific methylation information into mQTL mapping. In RNA sequencing studies, it has been well documented that incorporating ASE information can greatly improve the power of eQTL mapping [51, 76–78]. Our results confirm that this observation generalizes to mQTL mapping and provides substantial benefits over approaches that cannot or do not use allele-specific data. Notably, these benefits are not limited to the RRBS data we examined here: IMAGE can also be applied to analyze data generated via whole genome bisulfite sequencing (WGBS) [39] or by newer approaches that distinguish 5-hmc modifications from 5-mc modifications [43, 44]. Doing so would greatly facilitate detection of methylation-associated genetic variants genome-wide, including variants associated with different types of methylation marks.

Notably, although secondary to the methods advance itself, our real data applications show that mQTL mapping can be successfully executed using bisulfite sequencing data alone, in the absence of independently generated genotype data. Specifically, we used the same bisulfite sequencing data set to both extract methylation measurements and call SNP genotypes. Our approach dovetails with previous observations that accurate genotyping data can be obtained from RNA sequencing data [95], bisulfite sequencing data [78], or ChIP sequencing data [96], which simultaneously reduces experimental cost and increases the utility of different sequencing data types. Because of these benefits, molecular QTL mapping without separate DNA sequencing or genotyping is gaining popularity [97]. For example, a recent study performed eQTL mapping and ASE analysis using RNA sequencing alone and demonstrated that this strategy achieves approximately 50% power compared to traditional eQTL mapping strategies that rely on independently derived genotype data, even though it only uses the 12.66% of SNPs represented in blood-derived RNAseq reads [45]. Here, we also show that genotyping and phenotyping from the same data set can facilitate well-powered mQTL mapping. Notably, unlike RNA-seq data, because allele-specific methylation information is represented as the ratio between methylated reads and total reads mapped to the same allele, our approach is also less likely to be affected by allele-specific mapping biases (mitigating another argument for generating independent genotype data). Thus, our mQTL mapping approach has the potential to both increase the utility and applicability of functional genomic data types and improve accessibility of this type of analysis across species.

Our method is not without limitations. For example, to enable ASM-assisted mQTL mapping, our method makes a key modeling assumption: that the allelic effect size estimated from heterozygotes is equivalent to the genotype effect size estimated from mQTL mapping across all genotype classes. This assumption is generally satisfied for *cis* genetic effects when the SNP is close to the CpG site [98], and is shared, for gene expression phenotypes, with ASE-assisted eQTL mapping methods (e.g. TreCASE and WASP [51, 52]). However, in rare occasions, the equal effect size assumption may be violated. For example, if ASM arises because of genomic imprinting instead of sequence variation, the allelic effect size may be much smaller than the mQTL effect size obtained across all individuals. Such a violation would lead IMAGE to lose power relative to classical mQTL mapping approaches. Notably, imprinted regions are quite rare in vertebrate genomes (less than 1% of genes are imprinted) [99]. However, excluding imprinted loci prior to IMAGE mapping or substituting the IMAGE-I approach for these loci may slightly improve performance. Additionally, in unphased data, an important limitation of IMAGE is that it can only be used to analyze adjacent SNP-CpG pairs that are covered by the same sequencing reads. Analyzing only adjacent SNP-CpG pairs can limit the discovery of mQTLs. Therefore, it would be important to extend IMAGE to analyze distant SNP-CpG pairs in unphased data, using, for example, strategies presented in [100]. Certainly, if SNP data can be phased, IMAGE can also be applied to analyze SNP-CpG pairs that are separated by longer distances. In principle, using phased data could improve mQTL mapping power even further, if physically linked CpG sites display consistent ASM. Because the baboon and wolf data we analyzed here are not associated with an extensive genetic reference panel, we did not attempt to extend our analysis to phased data. Nevertheless, exploring the benefits of phased data or extending IMAGE to analyzing distant SNP-CpG pairs in unphased data is an important future direction.

Another limitation of IMAGE is that type I error may not be well controlled when methylation background heritability is high (>0.6, Additional file 3: Table S5), when the sample size is small (<100, Additional file 3: Table S6), or when the genotype minor allele frequency is low (<0.1, Additional file 3: Table S7). As a result, we recommend calibrating the false discovery rate against a permutation-derived empirical null, as we have done here (we note that calibrating against permutations has become an increasingly common approach in functional genomic mapping studies in any case [101, 102]). Finally, while our method is reasonably efficient and can be readily applied to analyze hundreds of individuals and tens of thousands of SNP-CpG pairs (Table 1), new algorithms will be needed to adapt IMAGE to data sets that are orders of magnitude larger.

**Table 1:** Computational time for analyzing differently sized data sets, for count-based mQTL mapping methods. Computing time is based on analysis of 100,000 SNP-CpG pairs with baseline simulation parameters and varying sample size, using a single thread on a Xeon E5-2683 2.00GHz processor.

| Method | Sample Size | Time (min) |
| --- | --- | --- |
| IMAGE | 50 | 295.87 |
|  | 100 | 410.10 |
|  | 150 | 543.48 |
| IMAGE-I | 50 | 81.10 |
|  | 100 | 63.93 |
|  | 150 | 108.27 |
| IMAGE-A | 50 | 16.63 |
|  | 100 | 36.51 |
|  | 150 | 69.25 |
| MACAU | 50 | 214.85 |
|  | 100 | 476.58 |
|  | 150 | 883.60 |
| BB | 50 | 417.63 |
|  | 100 | 2029.44 |
|  | 150 | 3967.03 |

Nevertheless, in its current form, IMAGE is well-suited to analyzing sequencing-based DNA methylation data sets of the size and scale typically generated in recent studies [103]. Thus, it can be flexibly deployed to investigate the genetic architecture of gene regulatory variation, the relative role of genes and the environment in shaping the epigenome, or the mediating role of DNA methylation in linking environmental conditions to downstream phenotypes, including human disease (e.g., via Mendelian randomization or related approaches [104, 105]).

## Materials and Methods

### Method Overview

Both mQTL mapping and ASM analysis examine one CpG site-SNP pair at a time to identify SNPs associated with DNA methylation levels. However, these two approaches rely on different information to model the genotype-DNA methylation level relationship. Specifically, mQTL mapping focuses on modeling the methylated read counts and total read counts at the individual level across all samples, without differentiating between the contributions from the two alleles contained within each individual. In contrast, ASM analysis focuses on modeling methylated read counts and total read counts in an allele-specific fashion, restricting it to heterozygotes for the SNP of interest (otherwise, the contributions of each allele cannot be decoupled). mQTL mapping has the benefit of using the entire sample, not just heterozygotes. In contrast, ASM has the benefit of internal control, since both alleles within each heterozygote experience the same genetic and environmental background.

To take advantage of both approaches, IMAGE independently models each CpG-SNP site pair. For each individual measured at a CpG-SNP pair, we denote *y*_*i*_ and *r*_*i*_ as the methylated read count and total read count for the *i*^th^ individual (combined across alleles), for *i* = 1, …, *n*. We denote the corresponding methylated and total read counts mapped to each of the two alleles of the *i*^th^ individual as *y*_*ii*_ and *r*_*il*_, for *l* = 1 or 2. Thus, *y*_*i*_ = *y*_*i1*_ + *y*_*i2*_ and *r*_*i*_ = *r*_*i1*_ + *r*_*i2*_. Note that *y*_*il*_ and *r*_*il*_ are only observed in heterozygotes, so are treated as missing data in homozygotes (more details below). We then model the methylated read counts for each allele as a function of the total read counts for the same allele using a binomial model:

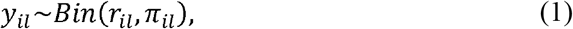

where π_*ii*_ is the true methylation level for the *l*^th^ allele in the *i*^th^ individual. We further model the logit-transformed methylation proportion π_*il*_ as a function of allele genotype:

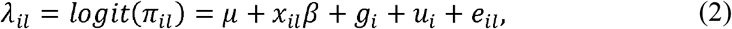

where µ is the intercept; *x*_*il*_ is the *l*^th^ allele type for the *i*^th^ individual for the SNP of interest (*x*_*il*_ = 0 or 1, corresponding to the reference allele and alternative allele, respectively); and β is the corresponding allele/genotype effect size. In addition to these fixed effects, we model three random effects to account for different sources of over-dispersion. Specifically, *g*_*i*_ represents the genetic background/polygenic effect on DNA methylation for the *i*^th^ individual and can be used to account for kinship or other population structure in the sample. We assume 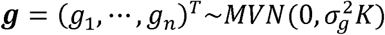, where *K* is a known *n* by *n* genetic relatedness matrix that can be estimated either from genotype or pedigree data. *u*_i_ represents individual-level environmental effects that we assume are independent across individuals but shared between the two alleles within the same individual. We assume 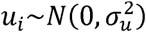. Finally, *e*_*ii*_ represents the residual error and is used to account for independent noise that varies across both individuals and alleles (e.g. stochastic events). We assume 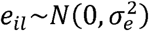. We standardize the genetic relatedness matrix *K* to ensure that the mean of the diagonal elements of *K* equals 1, or 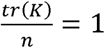. When this is the case, 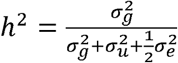, and can be interpreted as the approximate background heritability of DNA methylation levels (details in the Additional file 1: Supplementary Text). Here, the background heritability represents the proportion of variance in the latent parameter} explained by the genetic effects from all SNPs other than the SNP of focus (i.e. x). Therefore, the background heritability is the usual heritability minus the genetic effect of x. Our primary goal is to test the null hypothesis that genotype is not associated with methylation levels, or equivalently, H_0_: β = 0.

While the above model is fully specified for heterozygous individuals, it is not fully specified in homozygotes, where *y*_*il*_ and *r*_*il*_ are not observed. For homozygotes, only the sums of the reads across both alleles, *y*_*i*_ = *y*_*i1*_ + *y*_*i2*_ and *r*_*i*_ = *r*_*i1*_ + *r*_*i2*_, are observed. Therefore, for homozygotes, we derive a model for *y*_*i*_ and *r*_*i*_ based on equation (1) by summing over all possible values of *y*_*il*_ and *r*_*il*_:

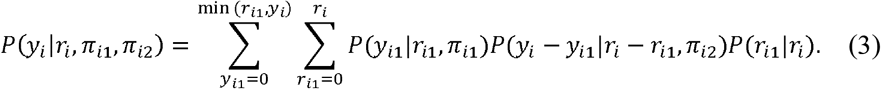

In equation (3), we assume that the model specified in equation (1) for the two alleles are independent of each other; thus 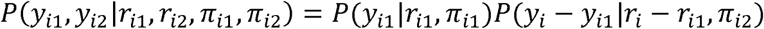. We further assume that 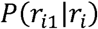 follows a binomial distribution *r*_*i1*_~*Bin*(*r*_*i*_, 0.5), which reflects the assumption that both alleles are equally likely to be represented in the sequencing data. Even with these two assumptions, the probability 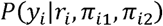 in equation (3) does not have an analytic form and can only be evaluated numerically, which is highly computationally inefficient for parameter estimation and inference. To enable scalable computation, we therefore approximate the distribution in equation (3) using a binomial distribution (details in the Additional file 1: Supplementary Text). Numerical simulations demonstrate the accuracy of this approximation across a range of settings (Additional file 2: Fig. S9).

The model defined in equations (1), (2) (for heterozygous individuals) and (3) (for homozygous individuals) allows us to perform ASM-assisted mQTL mapping to identify SNPs associated with DNA methylation levels. Due to the random effects terms in the model, the joint likelihood based on these equations consists of a high-dimensional integration that cannot be solved analytically. Here, we rely on the penalized quasilikelihood (PQL) algorithm that is commonly used for fitting generalized linear mixed models [64, 80, 81] to perform parameter estimation. Based on the parameter estimates, we further calculate a Wald statistic for testing the null hypothesis that *H*_0_: *β* = 0 and obtaining a corresponding p-value.

We refer to the above model as IMAGE, which is implemented as a freely available R software package at www.xzlab.org/software.html.

## Simulations

We performed simulations to examine the effectiveness of our method and compare it with other approaches. To do so, we first randomly selected 150 individuals from the 1958 birth cohort study, which is a part of the control samples that were used in the Wellcome Trust Case Control Consortium Study (WTCCC) [82]. We then obtained genotypes for 394,117 SNPs on chromosome 1 for these selected individuals. In the simulations, we examined the influence of sample size on power by choosing three different sample sizes: n = 50, 100 or 150. For n = 150, we used all 150 samples; for n < 150, we randomly selected the corresponding number of individuals from the 150 samples. For each simulation replicate, we computed the genetic relatedness matrix K from the SNP data using GEMMA [83–85]. We examined the influence of SNP minor allele frequency (MAF) on power by dividing the 394,117 SNPs into three different MAF bins: an MAF bin centered on 0.1, which contains SNPs with an MAF between 0.05 and 0.15 (p=100,631); an MAF bin centered on 0.3, which contains SNPs with an MAF between 0.25 and 0.35 (p=51,800); and an MAF bin including 0.5 which contains SNPs with an MAF between 0.45 and 0.50 (p=23,619). To simulate SNP-CpG site pairs, given a combination of sample size and MAF bin, we randomly selected one SNP from the appropriate MAF bin and simulated methylation counts and total read counts based on the following procedure.

For the total read counts, we first used a negative binomial distribution *NB*(*TR*, *ϕ*) to simulate the total read count *r*_*i*_ for each individual. Here, *TR* is the mean parameter and *ϕ* is the dispersion parameter. We set *TR* = 10, 20, or 30, close to the median estimate across all CpG sites from the baboon data (details of the data are described in the next section; median estimate in the real data = 23). We set *ϕ* = 3, which is close to the median estimates obtained from the baboon data (median estimate in the real data = 2.80). To obtain the total read count mapped to each of the two alleles, we further simulated a proportion parameter *q*_*i*_, which represents the proportion of reads mapped to one allele out of the two alleles. Specifically, *q*_*i*_ was simulated from a beta distribution Beta(a, b), where we set the shape parameters *a* and *b* to both be 10, so that the simulated *q*_*i*_ is symmetric around 0.5 and is within the range of (0.3, 0.7) in 93.6% of cases. With *r*_*i*_ and *q*_*i*_, we simulated the total read count mapped to one of the two alleles from *r*_*i1*_~*Bin*(*r*_i_, *q*_i_) and set the total read count mapped to the other allele as *r*_*i2*_ = *r*_*i*_- *r*_*i1*_.

For the methylated read counts, we performed simulations using a combination of five parameters. These five parameters include the intercept µ, which characterizes the baseline methylation level (interpretable as the mean methylation level within a given population); *h*^2^, which represents background heritability; *σ*^2^, which is the over-dispersion variance; ρ, which characterizes the proportion of common environmental variance (i.e., for those effects that are shared between the two alleles in each individual) with respect to both the common environmental variance and the independent environmental variance that are independent between both individuals and alleles within individuals; and *PVE*, which represents genotype effect size in terms of proportion of phenotypic variance explained (PVE) by genotype. With these four parameters, we first simulated the genetic random effects **g** = (*g*_1_,., *g*_*n*_)^*T*^ (an *n*-vector) across all individuals from a multivariate normal distribution with covariance 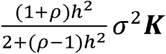 to guarantee that the background heritability for our population of simulated individuals is *h*^2^ (details in the Additional file 1: Supplementary Text). For each individual at a time, we then simulated the environmental random effects (*e*_i1_,*e*_i2_) and *u*_i_ together as a bivariate vector 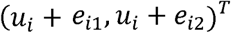 from a bivariate normal distribution with a covariance Σ, where 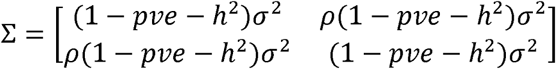.

For sites where methylation level was not associated with genotype, the SNP effect β was set to zero and the background genetic effects, environmental effects, and an intercept (μ) were then summed together to yield the latent variable π_il_ through 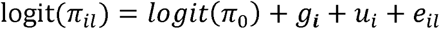 for the *l*^th^ allele in *i*^th^ individual. For sites with true mQTL, we used 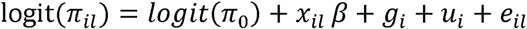 to yield the latent variable π_il_, where *x*_il_ is the allele genotype for the *l*^th^ allele in the *i*^th^ individual. We randomly draw 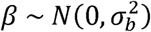 for each CpG site in turn, where 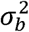 is set to ensure that genetic effects explain a fixed *PVE* in logit(*n*_il_), on average. We set *PVE* to be 5%, 10%, or 15% to represent different mean mQTL effect sizes and we derive 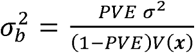, where the function *V*(•) denotes the sample variance computed across individuals with X being a genotype vector of size *n*. Finally, we simulated the methylated read counts for each allele based on a binomial distribution with a rate parameter determined by the total read counts r_i_ and the methylation proportion π_il_; that is, y_il_~Bin(r_il_,π_il_) for the l^th^ allele in i individual. For heterozygotes we retained the allele-level data (*y*_i1_,*y*_i2_) and (*r*_i1_, *r*_i2_). For homozygotes we collapsed the allele-level data into individual-level data, *y*_i_ = *y*_i1_ + *y*_i2_ and *r*_i_ = *r*_i1_ + *r*_i2_.

Using the procedure described above, we first simulated data under a baseline simulation scenario of *n* = 100, *h*^2^ = 0.3 *π*_0_ = 0.5 *MAF* = 0.3, ρ= 0, *TR*^2^ = 20, σ= 0.7, and PVE = 0.1 for mQTL sites. We then varied one parameter at a time to generate different simulation scenarios to examine the influence of each parameter, following [40]. Here, we varied the baseline methylation level *π*_0_ to be either 0.1, 0.5, or 0.9 to represent low, moderate, or high levels of DNA methylation. We varied *h*^2^ = 0.0, 0.3, or 0.6 to represent no, medium, or high background heritability. We varied *σ*^2^ = 0.3, 0.5, or 0.7, to represent different levels of over-dispersion. We varied p= 0, 0.3, or 0.9 to represent different levels of common environment influence. For each simulated combination of parameters, we performed 10 simulation replicates consisting of 10,000 CpG sites each. Among these sites, DNA methylation levels at 1,000 of them were associated with the SNP genotype (β ≠ 0) while DNA methylation levels for the remaining 9,000 were not (β = 0).

### Baboon RRBS data

We applied our method to a bisulfite sequencing data set from 69 wild baboons from the Amboseli ecosystem in Kenya [40, 45]. These data were generated using RRBS on the Illumina HiSeq 2000 platform, with 100 bp single-end sequencing reads. We obtained the raw fastq files from NCBI (accession number PRJNA283632), removed adaptor contamination and low-quality bases using the program Trim Galore (version 0.4.3) [106], and then mapped reads to the baboon reference genome (*Panu2.0*) using BSseeker2 [107] (Additional file 2: Fig. S3; more details in Additional file 1: Supplementary Text). After removing two samples that had extremely low sequencing read depths (57,734 and 58,070 reads, respectively), sequencing read depth ranged from 5.00 to 79.78 million reads (median = 24.48 million reads; sd = 13.69 million).

We performed SNP calling in the bisulfite sequencing data using CGmaptools, a SNP calling program specifically designed for bisulfite sequencing data. CGmaptools examines one individual at a time using the BayesWC SNP calling strategy [78]. Following the authors’ recommendations, we used a conservative error rate of 0.01 and a dynamic p-value to account for different read depth per site. Further, we modified the source code to make CGmaptools output homozygous reference genotypes as well. After SNP calling, we indexed and merged variant call files (VCFs) using VCFtools [108]. We then obtained a common set of SNPs where the position was called in at least 50% individuals (including homozygous reference calls). For each individual, we filtered out SNPs that were called using less than three reads. For each SNP, we filtered out variants that had an estimated MAF < 0.05. Finally, we filtered out 989 multiallelic SNPs to obtain a final call set of 289,103 analysis-ready SNPs (mean = 203,864 SNPs typed per sample; median = 204,554; sd = 34,768). We computed the genetic relatedness matrix ***K*** in GEMMA, using this SNP data set.

To validate the SNP genotype data, we compared the variants identified from the bisulfite sequencing data to a set of previously identified SNP variants in baboons [109]. These previously identified SNPs were obtained from 44 different wild baboons from East Africa, including members of the baboon population from which the RRBS data were generated but also members of baboon populations outside Amboseli, via low-coverage DNA sequencing (range: 0.6x to 4.35x; median = 1.91x; sd = 0.77x). This data set identified a total of 24,770,393 SNPs, with an average of 17,725,780 SNPs genotyped per individual (median = 18,139,340; sd = 4,315,590). Because of the low sequencing depth in the DNA sequencing data set, we expected that variants called from the bisulfite sequencing data would not completely overlap with variants identified from the DNA sequencing data. Indeed, we found that 50.9% of our called variants are located at a known variant from the DNA sequencing study, with the remaining SNPs being novel. Importantly, among overlapping variants, 99.5% have the same alternate allele, in support of the accuracy of SNP calling from bisulfite sequencing data. Additionally, we observe more overlap in called variants with higher alternate allele frequency, reaching 72.5% for variants with an alternate allele frequency > 0.5 in the RRBS data (Additional file 2: Fig. S10A). The allele frequency estimates from the two data sets for overlapping variants are reasonably well correlated (Spearman correlation r = 0.551; *p*-value < 2.2 × 10^−16^; Additional file 2: Fig. S10B).

In addition to genotyping, we used CGmaptools to obtain CpG-SNP pairs where the SNP and CpG site were profiled on the same sequencing read. The distance between the SNP-CpG site pairs ranges from 1 bp to 104 bp, with a median distance of 37 bp (mean = 39.75 bp; sd = 26.15 bp; Additional file 2: Fig. S10C). We extracted the methylation level estimates for each CpG site in the form of the number of methylated read counts and the number of total read counts, at the individual level for homozygotes and for each allele separately for heterozygotes. We obtained a total of 522,965 SNP-CpG pairs, with 82,217 unique SNPs and 391,137 unique CpG sites. Following [49], we excluded CpG sites (i) that were measured in less than 20 individuals; (ii) where methylation levels fell below 10% or above 90% in at least 90% of measured individuals; (iii) that had a mean read depth less than 5; or (iv) that were paired with a SNP with MAF<0.05 across individuals for whom DNA methylation estimates were available. To avoid potential mapping bias, we also excluded CpG sites with apparent differences in methylation levels between reference and alternate alleles that were larger than 0.6. Note that excluding these sites is a conservative strategy and may remove truly associated SNP-CpG pairs where mQTL are unusually large effect size. After filtering, our final data consisted of 49,196 SNP-CpG pairs, with 13,753 unique SNPs and 45,210 unique CpG sites, and an average of 33,539 SNP-CpG pairs measured per individual.

For these SNP-CpG pairs, the median number of reads per SNP across all individuals was 23 (mean = 31.21; sd = 30.08), and the median number of reads per allele was 13 in heterozygous individuals (mean = 18.75; sd = 19.75). To check the quality of DNA methylation estimates for these CpG sites, we examined their distribution across individuals. Similar to other RRBS data sets [110], we observed a bimodal distribution pattern of methylation levels, including a large number of hypomethylated and hypermethylated CpG sites (Additional file 2: Fig. S10D). Next, we examined the accuracy of methylation measurements obtained from our pipeline by comparing the mean methylation at each CpG site obtained here to those estimated in a previous study that focused on a subset of 61 individuals but used a different mapping and DNA methylation estimation pipeline [111]. As expected, the overall distribution of DNA methylation levels is almost identical between our pipeline and the previous study for the 15,605 overlapping sites (Additional file 2: Fig. S10E). In addition, site-specific DNA methylation level estimates are highly correlated (Spearman correlation r = 0.855, *p*-value < 2.2 × 10^−16^; Additional file 2: Fig. S10E). Finally, we checked whether our data suggest mapping bias in favor of the reference allele. Among the CpG sites we analyzed, we observed no bias in methylation level estimates between the reference and the alternate alleles (Additional file 2: Fig. S10F).

We applied five different approaches (details in the Results), together with our primary IMAGE method, to analyze the baboon DNA methylation data. Most of these methods are count based, and algorithms for count based models can be computationally unstable in the presence of covariates. To control for confounding effects from covariates, for each SNP in turn, we removed the effects of age, sex and the top two methylation principal components based on M-values [112] and used the genotype residuals for analysis. One method, IMAGE-A, requires a relatively large number of heterozygous individuals and was thus only applied to analyze sites for which we identified at least 5 heterozygotes (38,250 SNP-CpG pairs). All other methods were applied to all 49,196 SNP-CpG pairs. Because different methods have different type I error control and one method (IMAGE-A) analyzes a different number of SNP-CpG pairs, to ensure fair comparison, we performed permutations to construct empirical null distributions. Specifically, we combined the count data from the heterozygotes (*y*_i1_, *y*_i2_),(*r*_i1_,*r*_i2_) with the count data from the homozygotes (*y*_i_,*r*_i_), treated the two alleles of each heterozygote as two samples and treated each homozygote as one sample, permuted the sample label 10 times to create null permutations, and applied each method to analyze the permuted data. We note that an alternative permutation strategy would be to permute (*y*_*i*_,*r*_i_) along with covariates across individuals. In this strategy, the number of methylated reads for each allele (out of total reads for each allele) in heterozygotes could then be sampled from a binomial distribution with probability 0.5, conditional on *y*_i_ and r_i_ − y_i_ respectively. This alternative strategy is not ideal for small sample sizes, but is likely to work well for large samples (approximately n>150). Therefore, we have also implemented this alternative permutation strategy in the software and recommend users to explore both strategies and select one that performs the best for their data. Regardless of which permutation strategy one uses, the statistics from the permuted data allowed us to construct an empirical null distribution. With the empirical null distribution, we estimated the empirical false discovery rate (FDR) for different methods at different p-value thresholds. We then compared the number of associations detected by different methods at a fixed FDR cutoff.

Finally, following [86], we annotated CpG sites into four categories based on genomic locations obtained from the UCSC Genome Browser: island, shore, shelf and open sea. CpG islands are defined as short (approximately 1 kb) regions of high CpG density in an otherwise CpG-sparse genome [113]. A large proportion of CpG islands have been shown to be associated with gene promoters [114, 115]. The methylation level at the CpG islands is often associated with transcription repression [116, 117]. CpG shores are defined as the 2 kb of sequence flanking a CpG island and CpG shelfs are defined as the 2 kb of sequence further flanking CpG shores. Both CpG shores and shelfs have been reported to be more dynamic than the CpG island itself [90, 118, 119]. The methylation variation at shores and shelfs have been associated with various diseases. Finally, the remaining regions outside of CpG island/shore/shelf are denoted as open seas [120]. We downloaded the CpG island annotations for *Panu2.0* directly from UCSC; annotated the 2 kb region upstream and downstream of the CpG island boundaries as the shore; annotated the 2 kb regions upstream and downstream of the CpG shores as the CpG shelves; and annotated the remaining regions as open sea (Figure 3E).

### Wolf RRBS data

We also applied our method to analyze a second bisulfite sequencing data set, from 63 grey wolves from Yellowstone National Park in the United States [46, 94]. The wolf data are RRBS data collected on the Illumina HiSeq 2500 platform using 100 bp single end sequencing reads. We obtained bam files for 35 individuals from NCBI (accession number PRJNA299792) [46] and the fastq files for the remaining individuals from accession number PRJNA488382 [94]. We processed all files using the same procedure described in the previous section, using Trim Galore and BSseeker2, with the dog genome *canFam 3.1* [121] as the reference genome. Per-individual sequencing read depth ranges from 9.53 to 75.18 million reads per individual (median = 31.36 million reads; sd = 12.91 million). We used the same SNP calling procedure described for baboons and applied the same filtering criteria to obtain a final call set of 518,774 SNPs, with an average of 360,063 SNPs genotyped per individual (median = 440,898; sd = 103,522). We also computed the genetic relatedness matrix K with these SNPs using GEMMA.

To validate variants identified in the wolf data set, we compared the called variants from the bisulfite sequencing data to an existing SNV data base from the current Ensembl release for the dog genome *canFam 3.1*. We found that 17.9% of variants overlapped with known variants from Ensembl. Importantly, among overlapping variants, 99.1% of them have the same alternative allele as reported in Ensembl. In addition, the proportion of overlapping variants increases with increasing alternate allele frequency and reaches 41.3% when we focus on variants that have an alternate allele frequency > 0.5 in the RRBS data (Additional file 2: Fig. S11A).

We followed the same procedure described for baboons to extract methylation measurements on SNP-CpG pairs. In the wolves, the distance between SNP-CpG site in each pair ranges from 1 bp to 103 bp, with a median of 35 bp (mean = 38.41bp; sd = 25.63bp; Additional file 2: Fig. S11B). We obtained a total of 861,474 SNP-CpG pairs, representing 144,670 unique SNPs and 684,681 unique CpG sites. Following quality control filtering, we obtained a final set of 279,223 SNP-CpG pairs, representing 77,039 unique SNPs and 242,784 unique CpG sites, with an average of 179,412 SNP-CpG pairs measured per individual. In this set, the median number of reads per SNP across all individuals is 25 (mean = 31.16; sd = 29.33) and the median number of reads per allele is 14 in heterozygotes (mean = 17.45; sd = 18.90). Methylation levels across sites display the expected bimodal distribution pattern (Additional file 2: Fig. S11C), and we observed no bias in methylation level estimates between the reference and the alternate alleles (Additional file 2: Fig. S11D).

We applied the same analysis procedure to analyze the wolf data as we did for the baboon data set. IMAGE-A was used to analyze 236,092 SNP-CpG pairs where the data set included at least 5 heterozygotes while the other methods were applied to all 279,223 SNP-CpG pairs. We used permutation to construct empirical null distributions for FDR control and controlled for the effects of sex and the top two methylation principal components in the same procedure described in the baboon data. Finally, we annotated CpG sites into island, shore, shelf and open sea categories as described above, based on the *canFam3.1* genome.

## Additional files

Additional file 1: Supplementary text on IMAGE modeling and inference details. (DOCX 43kb)

Additional file 2: Supplementary figures on the performance evaluation of IMAGE and on the quality control of the real data applications. (DOCX 3,783kb)

Additional file 3: Supplementary tables on functional enrichment analyses and type I error control examination. (DOCX 42kb)

## Ethics approval and consent to participate

Not applicable.

## Consent for publication

Not applicable.

## Availability of data and materials

Baboon RRBS fastq files are available in the Sequence Read Archive (SRA) of NCBI under accession PRJNA283632 [40, 45]. Wolf RRBS bam files for 35 wolves are available under accession PRJNA299792 [46] and the fastq files for the other 27 wolves are available under accession PRJNA488382 [94]. The Trim Galore! Software is available from https://www.bioinformatics.babraham.ac.uk/projects/trim_galore/ [106]. The BS Seeker 2 software is available from http://pellegrini-legacy.mcdb.ucla.edu/bs_seeker2/[107]. The VCFtools software is available from http://vcftools.sourceforge.net/[108]. The CGmaptools software is available from https://cgmaptools.github.io/[78]. The GEMMA[83–85], MACAU[40, 57], BB[40], and PQLseq[64] software packages are available from http://www.xzlab.org/software.html.

IMAGE is an open source R package that is freely available from GitHub [122] https://github.com/fanyue322/IMAGE, CRAN (https://cran.r-project.org/web/packages/IMAGE/index.html) and http://www.xzlab.org/software.html. Source code for the software release used in the paper has been placed into a DOI-assigning repository[123] (https://doi.org/10.5281/zenodo.3334384). The code to reproduce all the analyses presented in the paper are available on GitHub[124] (https://github.com/fanyue322/IMAGEreproduce) and deposited on Zenodo[125] (https://doi.org/10.5281/zenodo.3334388).

## Authors’ contributions

JT and XZ conceived the idea and provided funding support. YF and XZ developed the method and designed the experiments. YF implemented the software and performed simulations with assistance from SS and QP. YF and TPV performed real data analysis. YF, JT, and XZ wrote the manuscript with input from all other authors.

## Acknowledgements

We thank Yichen Si at the University of Michigan for helping with the initial exploration of the method. This study also makes use of data generated by the Amboseli Baboon Research Project (ABRP), data generated in the Yellowstone grey wolf population, and the Wellcome Trust Case Control Consortium (WTCCC). A full list of ABRP past and current funding sources and contributors to these data is available at http://amboselibaboons.nd.edu. A full list of the investigators who contributed to the generation of the WTCCC data is available from http://www.wtccc.org.uk/. Funding for the WTCCC project was provided by the Wellcome Trust under award 076113 and 085475.

## Reference

1. Murrell, A., et al., An association between variants in the IGF2 gene and Beckwith-Wiedemann syndrome: interaction between genotype and epigenotype. Human Molecular Genetics, 2004. 13(2): p. 247–255.

2. Jones, P.A., Functions of DNA methylation: islands, start sites, gene bodies and beyond. Nature Reviews Genetics, 2012. 13(7): p. 484–492.

3. Dayeh, T., et al., Genome-Wide DNA Methylation Analysis of Human Pancreatic Islets from Type 2 Diabetic and Non-Diabetic Donors Identifies Candidate Genes That Influence Insulin Secretion. Plos Genetics, 2014. 10(3): p. e1004160.

4. Davegardh, C., et al., DNA methylation in the pathogenesis of type 2 diabetes in humans. Molecular Metabolism, 2018. 14: p. 12–25.

5. Bellamy, N., et al., Rheumatoid-Arthritis in Twins – a Study of Etiopathogenesis Based on the Australian-Twin-Registry. Annals of the Rheumatic Diseases, 1992. 51(5): p. 588–593.

6. Deapen, D., et al., A Revised Estimate of Twin Concordance in Systemic Lupus-Erythematosus. Arthritis and Rheumatism, 1992. 35(3): p. 311–318.

7. Jarvinen, P. and K. Aho, Twin Studies in Rheumatic Diseases. Seminars in Arthritis and Rheumatism, 1994. 24(1): p. 19–28.

8. Soriano-Tarraga, C., et al., Epigenome-wide association study identifies TXNIP gene associated with type 2 diabetes mellitus and sustained hyperglycemia. Human Molecular Genetics, 2016. 25(3): p. 609–619.

9. Dick, K.J., et al., DNA methylation and body-mass index: a genome-wide analysis. Lancet, 2014. 383(9933): p. 1990–1998.

10. Guay, S.P., et al., Epigenome-wide analysis in familial hyper-cholesterolemia identified new loci associated with high-density lipoprotein cholesterol concentration. Epigenomics, 2012. 4(6): p. 623–639.

11. Iossifov, I., et al., The contribution of de novo coding mutations to autism spectrum disorder. Nature, 2014. 515(7526): p. 216–221.

12. Amir, R.E., et al., Rett syndrome is caused by mutations in X-linked MECP2, encoding methyl-CpG-binding protein 2. Nature Genetics, 1999. 23(2): p. 185–188.

13. Cui, H.M., et al., Loss of IGF2 imprinting: A potential marker of colorectal cancer risk. Science, 2003. 299(5613): p. 1753–1755.

14. Vu, T.H., A.H. Nguyen, and A.R. Hoffman, Loss of IGF2 imprinting is associated with abrogation of long-range intrachromosomal interactions in human cancer cells. Human Molecular Genetics, 2010. 19(5): p. 901–919.

15. Byun, H.M., et al., Examination of IGF2 and H19 loss of imprinting in bladder cancer. Cancer Research, 2007. 67(22): p. 10753–10758.

16. Feinberg, A.P., M.A. Koldobskiy, and A. Gondor, Epigenetic modulators, modifiers and mediators in cancer aetiology and progression. Nature Reviews Genetics, 2016. 17(5): p. 284–299.

17. Timp, W. and A.P. Feinberg, Cancer as a dysregulated epigenome allowing cellular growth advantage at the expense of the host. Nature Reviews Cancer, 2013. 13(7): p. 497–510.

18. Chong, S.Y. and E. Whitelaw, Epigenetic germline inheritance. Current Opinion in Genetics & Development, 2004. 14(6): p. 692–696.

19. Anway, M.D., Epigenetic transgenerational actions of endocrine disruptors and male fertility. Science, 2010. 328(5979): p. 690–690.

20. Kaminsky, Z.A., et al., DNA methylation profiles in monozygotic and dizygotic twins. Nature Genetics, 2009. 41(2): p. 240–245.

21. Heijmans, B.T., et al., Heritable rather than age-related environmental and stochastic factors dominate variation in DNA methylation of the human IGF2/H19 locus. Human Molecular Genetics, 2007. 16(5): p. 547–554.

22. Ollikainen, M., et al., DNA methylation analysis of multiple tissues from newborn twins reveals both genetic and intrauterine components to variation in the human neonatal epigenome. Human Molecular Genetics, 2010. 19(21): p. 4176–4188.

23. Bell, J.T., et al., DNA methylation patterns associate with genetic and gene expression variation in HapMap cell lines. Genome Biology, 2011. 12(1): p. R10.

24. Bell, J.T., et al., Epigenome-Wide Scans Identify Differentially Methylated Regions for Age and Age-Related Phenotypes in a Healthy Ageing Population. Plos Genetics, 2012. 8(4): p. 189–200.

25. Lam, L.L., et al., Factors underlying variable DNA methylation in a human community cohort. Proceedings of the National Academy of Sciences of the United States of America, 2012. 109: p. 17253–17260.

26. Grundberg, E., et al., Global Analysis of DNA Methylation Variation in Adipose Tissue from Twins Reveals Links to Disease-Associated Variants in Distal Regulatory Elements. American Journal of Human Genetics, 2013. 93(6): p. 1158–1158.

27. Gutierrez-Arcelus, M., et al., Passive and active DNA methylation and the interplay with genetic variation in gene regulation. Elife, 2013. 2: p. e00523.

28. Polderman, T.J.C., et al., Meta-analysis of the heritability of human traits based on fifty years of twin studies. Nature Genetics, 2015. 47(7): p. 702–709.

29. Hannon, E., et al., Characterizing genetic and environmental influences on variable DNA methylation using monozygotic and dizygotic twins. PLoS genetics, 2018. 14(8): p. e1007544.

30. McRae, A.F., et al., Contribution of genetic variation to transgenerational inheritance of DNA methylation. Genome Biology, 2014. 15(5): p. R73.

31. Wagner, J.R., et al., The relationship between DNA methylation, genetic and expression inter-individual variation in untransformed human fibroblasts. Genome Biology, 2014. 15(2): p. R37.

32. Shi, J.X., et al., Characterizing the genetic basis of methylome diversity in histologically normal human lung tissue. Nature Communications, 2014. 5: p. 3365.

33. Luijk, R., et al., An alternative approach to multiple testing for methylation QTL mapping reduces the proportion of falsely identified CpGs. Bioinformatics, 2015. 31(3): p. 340–345.

34. Pai, A.A., et al., A Genome-Wide Study of DNA Methylation Patterns and Gene Expression Levels in Multiple Human and Chimpanzee Tissues. Plos Genetics, 2011. 7(2): p. e1001316.

35. Banovich, N.E., et al., Methylation QTLs Are Associated with Coordinated Changes in Transcription Factor Binding, Histone Modifications, and Gene Expression Levels. Plos Genetics, 2014. 10(9): p. e1004663.

36. Gibbs, J.R., et al., Abundant Quantitative Trait Loci Exist for DNA Methylation and Gene Expression in Human Brain. Plos Genetics, 2010. 6(5): p. e1000952.

37. Hannon, E., et al., Methylation QTLs in the developing brain and their enrichment in schizophrenia risk loci. Nature Neuroscience, 2016. 19(1): p. 48–54.

38. Day, K., et al., Heritable DNA Methylation in CD4(+) Cells among Complex Families Displays Genetic and Non-Genetic Effects. Plos One, 2016. 11(10): p. e0165488.

39. Cokus, S.J., et al., Shotgun bisulphite sequencing of the Arabidopsis genome reveals DNA methylation patterning. Nature, 2008. 452(7184): p. 215–219.

40. Lea, A.J., J. Tung, and X. Zhou, A Flexible, Efficient Binomial Mixed Model for Identifying Differential DNA Methylation in Bisulfite Sequencing Data. Plos Genetics, 2015. 11(11): p. e1005650.

41. Boyle, P., et al., Gel-free multiplexed reduced representation bisulfite sequencing for large-scale DNA methylation profiling. Genome Biology, 2012. 13(10): p. R92.

42. Gu, H.C., et al., Preparation of reduced representation bisulfite sequencing libraries for genome-scale DNA methylation profiling. Nature Protocols, 2011. 6(4): p. 468–481.

43. Liu, Y., et al., Bisulfite-free direct detection of 5-methylcytosine and 5-hydroxymethylcytosine at base resolution. Nature Biotechnology, 2019. 37(4): p. 424–429.

44. Yu, M., et al., Base-Resolution Analysis of 5-Hydroxymethylcytosine in the Mammalian Genome. Cell, 2012. 149(6): p. 1368–1380.

45. Tung, J., et al., The genetic architecture of gene expression levels in wild baboons. Elife, 2015. 4: p. e04729.

46. Koch, I.J., et al., The concerted impact of domestication and transposon insertions on methylation patterns between dogs and grey wolves. Molecular Ecology, 2016. 25(8): p. 1838–1855.

47. Chatterjee, A., et al., Mapping the zebrafish brain methylome using reduced representation bisulfite sequencing. Epigenetics, 2013. 8(9): p. 979–989.

48. Stubbs, T.M., et al., Multi-tissue DNA methylation age predictor in mouse. Genome Biology, 2017. 18(1): p. 68.

49. Schmitz, R.J., et al., Patterns of population epigenomic diversity. Nature, 2013. 495(7440): p. 193–198.

50. Alonso-Blanco, C., et al., 1,135 Genomes Reveal the Global Pattern of Polymorphism in Arabidopsis thaliana. Cell, 2016. 166(2): p. 481–491.

51. Hu, Y.J., et al., Proper Use of Allele-Specific Expression Improves Statistical Power for cis-eQTL Mapping with RNA-Seq Data. Journal of the American Statistical Association, 2015. 110(511): p. 962–974.

52. van de Geijn, B., et al., WASP: allele-specific software for robust molecular quantitative trait locus discovery. Nature Methods, 2015. 12(11): p. 1061–1063.

53. Cheung, W.A., et al., Functional variation in allelic methylomes underscores a strong genetic contribution and reveals novel epigenetic alterations in the human epigenome. Genome Biology, 2017. 18(1): p. 50.

54. Soneson, C. and M. Delorenzi, A comparison of methods for differential expression analysis of RNA-seq data. Bmc Bioinformatics, 2013. 14(1): p. 91.

55. Kvam, V.M., P. Lu, and Y.Q. Si, A Comparison of Statistical Methods for Detecting Differentially Expressed Genes from Rna-Seq Data. American Journal of Botany, 2012. 99(2): p. 248–256.

56. Zhang, Z.H., et al., A Comparative Study of Techniques for Differential Expression Analysis on RNA-Seq Data. Plos One, 2014. 9(8): p. e103207.

57. Sun, S.Q., et al., Differential expression analysis for RNAseq using Poisson mixed models. Nucleic Acids Research, 2017. 45(11): p. e106.

58. Feng, H., K.N. Conneely, and H. Wu, A Bayesian hierarchical model to detect differentially methylated loci from single nucleotide resolution sequencing data. Nucleic Acids Research, 2014. 42(8): p. e69–e69.

59. Sun, D.Q., et al., MOABS: model based analysis of bisulfite sequencing data. Genome Biology, 2014. 15(2): p. R38.

60. Dolzhenko, E. and A.D. Smith, Using beta-binomial regression for high-precision differential methylation analysis in multifactor whole-genome bisulfite sequencing experiments. Bmc Bioinformatics, 2014. 15(1): p. 215.

61. Park, Y. and H. Wu, Differential methylation analysis for BS-seq data under general experimental design. Bioinformatics, 2016. 32(10): p. 1446–1453.

62. Wu, H., et al., Detection of differentially methylated regions from whole-genome bisulfite sequencing data without replicates. Nucleic Acids Research, 2015. 43(21): p. e141–e141.

63. Weissbrod, O., et al., Association testing of bisulfite-sequencing methylation data via a Laplace approximation. Bioinformatics, 2017. 33(14): p. I325–I332.

64. Sun, S., et al., Heritability estimation and differential analysis of count data with generalized linear mixed models in genomic sequencing studies. Bioinformatics, 2019. 35(3): p. 487–496.

65. Dubin, M.J., et al., DNA methylation in Arabidopsis has a genetic basis and shows evidence of local adaptation. Elife, 2015. 4: p. e05255.

66. Orozco, L.D., et al., Epigenome-Wide Association of Liver Methylation Patterns and Complex Metabolic Traits in Mice. Cell Metabolism, 2015. 21(6): p. 905–917.

67. Li, Y.R., et al., The DNA Methylome of Human Peripheral Blood Mononuclear Cells. Plos Biology, 2010. 8(11): p. e1000533.

68. Peng, Q. and J.R. Ecker, Detection of allele-specific methylation through a generalized heterogeneous epigenome model. Bioinformatics, 2012. 28(12): p. I163–I171.

69. Fang, F., et al., Genomic landscape of human allele-specific DNA methylation. Proceedings of the National Academy of Sciences of the United States of America, 2012. 109(19): p. 7332–7337.

70. Kerkel, K., et al., Genomic surveys by methylation-sensitive SNP analysis identify sequence-dependent allele-specific DNA methylation. Nature Genetics, 2008. 40(7): p. 904–908.

71. Schalkwyk, L.C., et al., Allelic Skewing of DNA Methylation Is Widespread across the Genome. American Journal of Human Genetics, 2010. 86(2): p. 196–212.

72. Xie, W., et al., Base-Resolution Analyses of Sequence and Parent-of-Origin Dependent DNA Methylation in the Mouse Genome. Cell, 2012. 148(4): p. 816–831.

73. Shoemaker, R., et al., Allele-specific methylation is prevalent and is contributed by CpG-SNPs in the human genome. Genome Research, 2010. 20(7): p. 883–889.

74. Gertz, J., et al., Analysis of DNA Methylation in a Three-Generation Family Reveals Widespread Genetic Influence on Epigenetic Regulation. Plos Genetics, 2011. 7(8): p. e1002228.

75. Kaplow, I.M., et al., A pooling-based approach to mapping genetic variants associated with DNA methylation. Genome Research, 2015. 25(6): p. 907–917.

76. Sun, W., A Statistical Framework for eQTL Mapping Using RNA-seq Data. Biometrics, 2012. 68(1): p. 1–11.

77. Wilson, D.R., W. Sun, and J.G. Ibrahim, Mapping Tumor-Specific Expression QTLs In Impure Tumor Samples. bioRxiv, 2017: p. 136614.

78. Guo, W.L., et al., CGmapTools improves the precision of heterozygous SNV calls and supports allele-specific methylation detection and visualization in bisulfite-sequencing data. Bioinformatics, 2018. 34(3): p. 381–387.

79. Kumasaka, N., A.J. Knights, and D.J. Gaffney, Fine-mapping cellular QTLs with RASQUAL and ATAC-seq. Nature Genetics, 2016. 48(4): p. 473–473.

80. Breslow, N.E. and D.G. Clayton, Approximate Inference in Generalized Linear Mixed Models. Journal of the American Statistical Association, 1993. 88(421): p. 9–25.

81. Chen, H., et al., Control for Population Structure and Relatedness for Binary Traits in Genetic Association Studies via Logistic Mixed Models. American Journal of Human Genetics, 2016. 98(4): p. 653–666.

82. Power, C. and J. Elliott, Cohort profile: 1958 British Birth Cohort (National Child Development Study). International Journal of Epidemiology, 2006. 35(1): p. 34–41.

83. Zhou, X. and M. Stephens, Genome-wide efficient mixed-model analysis for association studies. Nature Genetics, 2012. 44(7): p. 821–824.

84. Zhou, X., P. Carbonetto, and M. Stephens, Polygenic Modeling with Bayesian Sparse Linear Mixed Models. Plos Genetics, 2013. 9(2): p. e1003264.

85. Zhou, X. and M. Stephens, Efficient multivariate linear mixed model algorithms for genome-wide association studies. Nature Methods, 2014. 11(4): p. 407–409.

86. Olsson, A.H., et al., Genome-Wide Associations between Genetic and Epigenetic Variation Influence mRNA Expression and Insulin Secretion in Human Pancreatic Islets. Plos Genetics, 2014. 10(12): p. e1004735.

87. Ronn, T., et al., A Six Months Exercise Intervention Influences the Genome-wide DNA Methylation Pattern in Human Adipose Tissue. Plos Genetics, 2013. 9(6): p. e1003572.

88. Volkov, P., et al., A Genome-Wide mQTL Analysis in Human Adipose Tissue Identifies Genetic Variants Associated with DNA Methylation, Gene Expression and Metabolic Traits. Plos One, 2016. 11(6): p. e0157776.

89. Antequera, F. and A. Bird, CpG islands as genomic footprints of promoters that are associated with replication origins. Current Biology, 1999. 9(17): p. R661–R667.

90. Irizarry, R.A., et al., The human colon cancer methylome shows similar hypo- and hypermethylation at conserved tissue-specific CpG island shores. Nature Genetics, 2009. 41(2): p. 178–186.

91. Ziller, M.J., et al., Charting a dynamic DNA methylation landscape of the human genome. Nature, 2013. 500(7463): p. 477–481.

92. Zhang, D.D., et al., Genetic Control of Individual Differences in Gene-Specific Methylation in Human Brain. American Journal of Human Genetics, 2010. 86(3): p. 411–419.

93. Zhi, D., et al., SNPs located at CpG sites modulate genome-epigenome interaction. Epigenetics, 2013. 8(8): p. 802–806.

94. Thompson, M.J., et al., An epigenetic aging clock for dogs and wolves. Aging-Us, 2017. 9(3): p. 1055–1068.

95. McKenna, A., et al., The Genome Analysis Toolkit: A MapReduce framework for analyzing next-generation DNA sequencing data. Genome Research, 2010. 20(9): p. 1297–1303.

96. del Rosario, R.C.H., et al., Sensitive detection of chromatin-altering polymorphisms reveals autoimmune disease mechanisms. Nature Methods, 2015. 12(5): p. 458–464.

97. Deelen, P., et al., Calling genotypes from public RNA-sequencing data enables identification of genetic variants that affect gene-expression levels. Genome Medicine, 2015. 7(1): p. 30.

98. Do, C., et al., Genetic-epigenetic interactions in cis: a major focus in the post-GWAS era. Genome Biology, 2017. 18(1): p. 120.

99. Wilkinson, L.S., W. Davies, and A.R. Isles, Genomic imprinting effects on brain development and function. Nature Reviews Neuroscience, 2007. 8(11): p. 832–843.

100. Knowles, D.A., et al., Allele-specific expression reveals interactions between genetic variation and environment. Nature Methods, 2017. 14(7): p. 699–702.

101. Zhang, Y. and J.S. Liu, Fast and Accurate Approximation to Significance Tests in Genome-Wide Association Studies. Journal of the American Statistical Association, 2011. 106(495): p. 846–857.

102. Segal, B.D., et al., Fast approximation of small p-values in permutation tests by partitioning the permutations. Biometrics, 2018. 74(1): p. 196–206.

103. Tobi, E.W., et al., DNA methylation signatures link prenatal famine exposure to growth and metabolism. Nature Communications, 2015. 6: p. 5592.

104. Richardson, T.G., et al., Systematic Mendelian randomization framework elucidates hundreds of CpG sites which may mediate the influence of genetic variants on disease. Human Molecular Genetics, 2018. 27(18): p. 3293–3304.

105. Huang, J.V., et al., DNA methylation in blood as a mediator of the association of mid-childhood body mass index with cardio-metabolic risk score in early adolescence. Epigenetics, 2018. 13(10-11): p. 1072–1087.

106. Krueger, F., Trim Galore!: A wrapper tool around Cutadapt and FastQC to consistently apply quality and adapter trimming to FastQ files. 2015, 0.4.

107. Guo, W.L., et al., BS-Seeker2: a versatile aligning pipeline for bisulfite sequencing data. Bmc Genomics, 2013. 14(1): p. 774.

108. Danecek, P., et al., The variant call format and VCFtools. Bioinformatics, 2011. 27(15): p. 2156–2158.

109. Wall, J.D., et al., Genomewide ancestry and divergence patterns from low-coverage sequencing data reveal a complex history of admixture in wild baboons. Molecular Ecology, 2016. 25(14): p. 3469–3483.

110. Lea, A.J., et al., Maximizing ecological and evolutionary insight in bisulfite sequencing data sets. Nature Ecology & Evolution, 2017. 1(8): p. 1074–1083.

111. Lea, A.J., et al., Resource base influences genome-wide DNA methylation levels in wild baboons (Papio cynocephalus). Molecular Ecology, 2016. 25(8): p. 1681–1696.

112. Du, P., et al., Comparison of Beta-value and M-value methods for quantifying methylation levels by microarray analysis. Bmc Bioinformatics, 2010. 11: p. 587.

113. Gardinergarden, M. and M. Frommer, Cpg Islands in Vertebrate Genomes. Journal of Molecular Biology, 1987. 196(2): p. 261–282.

114. Ioshikhes, I.P. and M.Q. Zhang, Large-scale human promoter mapping using CpG islands. Nature Genetics, 2000. 26(1): p. 61–63.

115. Saxonov, S., P. Berg, and D.L. Brutlag, A genome-wide analysis of CpG dinucleotides in the human genome distinguishes two distinct classes of promoters. Proceedings of the National Academy of Sciences of the United States of America, 2006. 103(5): p. 1412–1417.

116. Esteller, M., CpG island hypermethylation and tumor suppressor genes: a booming present, a brighter future. Oncogene, 2002. 21(35): p. 5427–5440.

117. Bird, A., DNA methylation patterns and epigenetic memory. Genes & Development, 2002. 16(1): p. 6–21.

118. Doi, A., et al., Differential methylation of tissue- and cancer-specific CpG island shores distinguishes human induced pluripotent stem cells, embryonic stem cells and fibroblasts. Nature Genetics, 2009. 41(12): p. 1350–1353.

119. Bibikova, M., et al., High density DNA methylation array with single CpG site resolution. Genomics, 2011. 98(4): p. 288–295.

120. Sandoval, J., et al., Validation of a DNA methylation microarray for 450,000 CpG sites in the human genome. Epigenetics, 2011. 6(6): p. 692–702.

121. Lindblad-Toh, K., et al., Genome sequence, comparative analysis and haplotype structure of the domestic dog. Nature, 2005. 438(7069): p. 803–819.

122. Fan Y, V.T., Sun S, Peng Q, Tung J, Zhou X. IMAGE: high-powered detection of genetic effects on DNA methylation using integrated methylation QTL mapping and allele-specific analysis. Source Code Github Repository, https://github.com/fanyue322/IMAGE(2019).

123. Fan Y, V.T., Sun S, Peng Q, Tung J, Zhou X. IMAGE: high-powered detection of genetic effects on DNA methylation using integrated methylation QTL mapping and allele-specific analysis. Source Code DOI, https://doi.org/10.5281/zenodo.3334384(2019).

124. Fan Y, V.T., Sun S, Peng Q, Tung J, Zhou X. IMAGE: high-powered detection of genetic effects on DNA methylation using integrated methylation QTL mapping and allele-specific analysis. Analysis Code Github Repository, https://github.com/fanyue322/IMAGEreproduce(2019).

125. Fan Y, V.T., Sun S, Peng Q, Tung J, Zhou X. IMAGE: high-powered detection of genetic effects on DNA methylation using integrated methylation QTL mapping and allele-specific analysis. Analysis Code Zenodo, https://doi.org/10.5281/zenodo.3334388(2019).

