## Supplemental Information for "IMAGE: High-powered detection of genetic effects on DNA methylation using integrated methylation QTL mapping and allele-specific analysis" (from another version of this preprint: version 1, 2019-04-22)

**Supplementary Text**

**1. Likelihood Approximation for Homozygous Individuals**

Here, we provide details on the approximation used for homozygous individuals. All notations follow the main text. Recall that we denote $y_{i}$ and $r_{i}$ as the methylated read count and total read count for *i*th individual, for $i=1,\cdots,n$. We denote the corresponding read counts mapped to the haplotype of the two alleles of *i*th individual as $y_{il}$ and $r_{il}$, for $l=1$ or 2; thus we have $y_{i}=y_{i1}+y_{i2}$ and $r_{i}=r_{i1}+r_{i2}$. Recall that, for homozygous individuals, we derive a model on $y_{i}$ and $r_{i}$ by summing over all possible values of $y_{il}$ and $r_{il}$:

$$P\left( y_{i} | r_{i},\pi_{i1},\pi_{i2} \right)=\sum_{y_{i1}=0}^{min(r_{i1},y_{i})} \sum_{r_{i1}=0}^{r_{i}} P(y_{i1}|r_{i1},\pi_{i1})P(y_{i}-y_{i1}|r_{i}-r_{i1},\pi_{i2})P(r_{i1}|r_{i}),$$

(1)

where $P(y_{i1}|r_{i1},\pi_{i1})$ follows a binomial distribution $y_{i1}\sim Bin(r_{i1},\pi_{i1})$; $P(y_{i}-y_{i1}|r_{i}-r_{i1},\pi_{i2})$ follows a binomial distribution $y_{i}-y_{i1}\sim Bin(r_{i}-r_{i1},\pi_{i2})$; and $P\left( r_{i1} | r_{i} \right)$ follows a binomial distribution $r_{i1}\sim Bin(r_{i},0.5)$. Here, $y_{i1}$ and $y_{i2}$ are conditionally independent given $\pi_{i1}$ and $\pi_{i2}$, though $\pi_{i1}$ and $\pi_{i2}$ are correlated with each other. To ensure scalable computation, we approximate the above $P\left( y_{i} | r_{i},\pi_{i1},\pi_{i2} \right)$ using a binomial mixed model in the following simple form:

$$y_{i}\sim Bin\left( r_{i},\tilde{\pi}_{i} \right), (2)$$

where $\tilde{\pi}_{i}$ is a mean that depends on $\pi_{i1}$ and $\pi_{i2}$, and thus is a function of ($\mu,\beta,{x_{i1},x_{i2},g}_{i},u_{i},e_{i1},e_{i2}$). We obtain an estimate of $\tilde{\pi}_{i}$ by matching the first moment in the likelihood function defined by equation (2) to the first moment in the likelihood function defined by equation (1). Specifically, the expectation of $y_{i}$ under equation (1) can be expressed as:

$$E\left( y_{i} \right)=E\left( y_{i1}+y_{i2} \right)=E\left( E\left( y_{i1}+y_{i2}|r_{i1},r_{i2} \right) \right)=r_{i}\frac{\pi_{i1}+\pi_{i2}}{2} .$$

Therefore, by matching moments, we have

$$\tilde{\pi}_{i}=(\pi_{i1}+\pi_{i2})/2$$

$$=(g^{-1}\left( \mu+x_{i1}\beta+g_{i}+{u_{i}+e}_{i1} \right)+g^{-1}\left( \mu+x_{i2}\beta+g_{i}+{u_{i}+e}_{i2} \right))/2, (3)$$

with

$$(\begin{matrix} {u_{i}+e}_{i1} \\ {u_{i}+e}_{i2} \end{matrix})\sim N(0,{(\sigma}_{u}^{2}+\sigma_{e}^{2})\left( \begin{matrix} 1 & \rho\\ \rho& 1 \end{matrix} \right)), \rho=\frac{\sigma_{u}^{2}}{\sigma_{u}^{2}+\sigma_{e}^{2}},$$

where $\mu$ is the intercept;$x_{il}$ is the *l*th allele type for *i*th individual for the SNP of interest ($x_{il}=0$ or 1, corresponding to the reference allele and alternative allele, respectively); $\beta$ is the corresponding allele/genotype effect size; $g_{i}$ represents the genetic background/polygenic effect of *i*th individual; $u_{i}$ represents the common individual-level environmental effect that varies across individuals but is shared between the two alleles within the same individual;$e_{il}$ represents the residual error and is used to account for independent environmental noise that varies across both individuals and alleles; $g(\cdot)$ denotes the logit link function.

Because we focus on homozygous individuals where $x_{i1}=x_{i2}=x_{i}$, we denote $\lambda_{0}=\mu+x_{i}\beta+g_{i}+u_{i}+(e_{i1}+e_{i2})/2$, which is the average of $\mu+x_{i1}\beta+g_{i}+{u_{i}+e}_{i1}$ and $\mu+x_{i2}\beta+g_{i}+{u_{i}+e}_{i2}$. Due to the $g^{-1}$ function, equation (3) is not in an easy-to-evaluate form. Instead, we further approximate each term in equation (3) through a first order Taylor expansion at $\lambda_{0}$:

$$g^{-1}\left( \mu+x_{i1}\beta+g_{i}+{u_{i}+e}_{i1} \right)\approx\frac{e^{\lambda_{0}}}{e^{\lambda_{0}}+1}\left( 1+\frac{\frac{e_{i1}-e_{i2}}{2}}{e^{\lambda_{0}}+1} \right),$$

$$g^{-1}\left( \mu+x_{i1}\beta+g_{i}+{u_{i}+e}_{i2} \right)\approx\frac{e^{\lambda_{0}}}{e^{\lambda_{0}}+1}\left( 1+\frac{\frac{e_{i2}-e_{i1}}{2}}{e^{\lambda_{0}}+1} \right).$$

Therefore, we have

$$\tilde{\pi}_{i}\approx\frac{e^{\lambda_{0}}}{e^{\lambda_{0}}+1},$$

or equivalently

$$g\left( \tilde{\pi}_{i} \right)\approx\lambda_{0}=\mu+x\beta+g_{i}+u_{i}+\frac{e_{i1}+e_{i2}}{2} (5)$$

for homozygous individuals. Note the above equation (5) holds exactly for heterozygous individuals, since we have $g\left( \pi_{il} \right)=\mu+x_{il}\beta+g_{i}+u_{i}+e_{il}$ for each allele of the heterozygous individuals (main text equation (2)). Subsequently, we can define an approximate heritability as

$$h^{2}=\frac{\sigma_{g}^{2}}{\sigma_{g}^{2}+\sigma_{u}^{2}+\sigma_{e}^{2}/2}. (6)$$

**2. IMAGE Inference Algorithm**

For homozygous individual $i$, if we denote the random effects for individual $i$ as $b_{i}=g_{i}+u_{i}+\frac{e_{i1}+e_{i2}}{2}$, we have:

| $y_{i}\sim Bin\left( r_{i},\tilde{\pi}_{i} \right), \mathrm{logit}\left( \tilde{\pi}_{i} \right)=\mu+x_{i}\beta+b_{i}.$ | $(7)$ |
| --- | --- |

For heterozygous individual $i$, the corresponding model is as follows:

| $y_{il}\sim Bin\left( r_{il},\pi_{il} \right), \mathrm{logit}\left( \pi_{il} \right)=\mu+x_{il}\beta+b_{il},l=1,2.$ | $(8)$ |
| --- | --- |

We treat the data for a homozygous individual, $\left( y_{i},r_{i} \right)$, as one observation. We treat the data for a heterozygous individual $\left( y_{i1},r_{i1},y_{i2},r_{i2} \right)$as two observations $(\left( y_{i1},r_{i1} \right),\left( y_{i2},r_{i2} \right)$). Given $n_{1}$ homozygous individuals (both types of homozygotes) and $n_{2}$ heterozygous individuals, we have$n^{'}=n_{1}+2n_{2}$ observations in total. To facilitate algebra derivation, we ordered observations by listing homozygous observations first, followed by heterozygous observations. The first and second moments in the likelihood functions defined in equations (7) and (8) can be written in vector and matrix forms as:

|  | $E\left( g\left( \boldsymbol{\pi} \right) \right)=\mu+\boldsymbol{x}\beta,$  $V\left( g\left( \pi\right) \right)=\sigma_{g}^{2}\left( \begin{matrix} \boldsymbol{K}_{n_{1}n_{1}} & \boldsymbol{K}_{n_{1}n_{2}}\otimes\begin{matrix} [1 & 1] \end{matrix} \\ \boldsymbol{K}_{n_{2}n_{1}}\otimes\left[ \begin{matrix} 1 \\ 1 \end{matrix} \right] & \boldsymbol{K}_{n_{2}n_{2}}\otimes\left[ \begin{matrix} 1 & 1 \\ 1 & 1 \end{matrix} \right] \end{matrix} \right){+\sigma_{u}^{2}\left( \begin{matrix} \boldsymbol{I}_{n_{1}\times n_{1}} & \boldsymbol{0}_{n_{1}\times n_{2}} \\ \boldsymbol{0}_{n_{2}\times n_{1}} & \boldsymbol{I}_{n_{2}\times n_{2}}\otimes[\begin{matrix} 1 & 1 \\ 1 & 1 \end{matrix}] \end{matrix} \right)+\sigma}_{e}^{2}\left( \begin{matrix} \frac{1}{2}\boldsymbol{I}_{n_{1}\times n_{1}} & \boldsymbol{0}_{n_{1}\times n_{2}} \\ \boldsymbol{0}_{n_{2}\times n_{1}} & \boldsymbol{I}_{n_{2}\times n_{2}}\otimes[\begin{matrix} 1 & 0 \\ 0 & 1 \end{matrix}] \end{matrix} \right),$ | $(9)$ |
| --- | --- | --- |

where both $g\left( \boldsymbol{\pi} \right)$ and $\boldsymbol{x}$ are $n^{'}$ dimensional vectors by stacking the corresponding values from homozygous observations and heterozygous observations; $\boldsymbol{K}_{n_{1}n_{1}}$, $\boldsymbol{K}_{n_{2}n_{2}}$, and$\boldsymbol{K}_{n_{1}n_{2}}$ are the corresponding kinship matrices computed among homozygous observations, among heterozygous observations, and between homozygous and heterozygous observations, respectively; and $\otimes$ denotes a Kronecker product.

We denote $\boldsymbol{b=g+u+e}$, as the $n^{'}$-vector of random effects. Our likelihood for all observations requires integrating out the random effects and thus is in a form of an n-dimensional integration

$$L=\int\prod_{i=1}^{n^{'}} f(\boldsymbol{y}_{\boldsymbol{i}}|\mu, \beta,\mathbf{b})f\left( \mathbf{b} | \sigma_{g}^{2},\sigma_{e}^{2}, \rho\right)d\boldsymbol{b}.$$

We develop an algorithm based on the penalized quasilikelihood (PQL) [1] for parameter inference for IMAGE. Briefly, PQL employs an iterative numerical optimization procedure. In each iteration, we introduce a set of pseudo-data $\tilde{\boldsymbol{y}}$ to replace the originally observed count data $\boldsymbol{y}$. The pseudo-data $\tilde{\boldsymbol{y}}$ is obtained based on a second order Taylor expansion using the conditional distribution $P(y_{i}|\mu,\beta,\boldsymbol{b})$ using the first and second order moments $E(y_{i}|\mu,\beta,\boldsymbol{b})$ and $V(y_{i}|\mu,\beta,\boldsymbol{b})$, both evaluated at the current estimates of the fixed coefficients as well as the random effects $\boldsymbol{b}$. With the pseudo-data, the complex likelihood function for the original data $\boldsymbol{y}$ is replaced by a much simpler linear mixed model (LMM) likelihood function for the pseudo-data $\tilde{\boldsymbol{y}}$, thereby alleviating much of the computational burden associated with our count model. With pseudo-data $\tilde{\boldsymbol{y}}$, we can perform inference and update parameters using the standard average information (AI) algorithm for LMMs [2-4]. By iterating between the approximation step of obtaining the pseudo-data $\tilde{\boldsymbol{y}}$ and the inference step of updating the parameter estimates via the AI algorithm, PQL allows us to perform inference in a computationally efficient fashion. Below, we describe the estimation and inference procedure in detail.

The observations $\left( y_{i},r_{i} \right) (i=1,2,\ldots,n^{'}; n^{'}=n_{1}+2n_{2}$; note our slight abuse of notation here) are independent conditional on the unobserved random effects $\boldsymbol{b}$ and the fixed effects $\mu+\boldsymbol{x}\beta,$ with conditional mean $E\left( y_{i} | \mu,\beta,\boldsymbol{b} \right)\mathbf{=}\pi_{i}r_{i}=g^{-1}(\mu+x_{i}\beta+b_{i})$ and conditional variance $V\left( y_{i} | \mu,\beta,\boldsymbol{b} \right)={\nu(\pi_{i})=\pi}_{i}(1-\pi_{i})r_{i}$. We use these two conditional moments to obtain the quasi-likelihood for $ith$ observation as $ql_{i}\left( \mu, \beta|\boldsymbol{b} \right)=\int_{y_{i}}^{\pi_{i}r_{i}} \frac{y_{i}-t}{\nu(t)}dt$, which serve as an approximation for the conditional likelihood. The joint likelihood function can be approximated by the joint quasi-likelihood function

$$ql\left( \mu,\beta,{\sigma_{g}^{2},\sigma}_{u}^{2},\sigma_{e}^{2} \right)=log\int\left( \prod_{i=1}^{n} ql_{i}\left( \boldsymbol{b,}\mu, \beta\right) \right)P\left( \boldsymbol{b|}{\sigma_{g}^{2},\sigma}_{u}^{2},\sigma_{e}^{2} \right)d\boldsymbol{b}.$$

We use Laplace approximation to further approximate the above function and obtain:

|  | $\tilde{ql}\left( \mu,\beta,{\sigma_{g}^{2},\sigma}_{u}^{2},\sigma_{e}^{2} \right)=\frac{1}{2}\log\left\vert\mathbf{VD}+\mathbf{I} \right\vert+\sum_{i=1}^{n} {ql}_{i}\left( \tilde{\boldsymbol{b}}, \mu,\beta\right)-\frac{1}{2}{\tilde{\boldsymbol{b}}}^{T}\mathbf{V}^{-1}\tilde{\boldsymbol{b}},$ | $(10)$ |
| --- | --- | --- |

where$\mathbf{V}=\sigma_{g}^{2}\boldsymbol{V}_{\mathbf{1}}+\sigma_{u}^{2}\boldsymbol{V}_{\mathbf{2}}+\sigma_{e}^{2}\boldsymbol{V}_{\mathbf{3}}$,$\tilde{\boldsymbol{u}}=\underset{\boldsymbol{u}}{\mathrm{argmax}} \left( \sum_{i=1}^{n} {ql}_{i}\left( \left. \mu,\beta\right|\boldsymbol{b} \right)-\frac{1}{2}\boldsymbol{b}^{T}\mathbf{V}^{-1}\boldsymbol{b} \right)$ and $\mathbf{D}=diag\left( 1/{g'\left( \lambda_{i} \right)} \right)$ is a diagonal weight matrix. $\boldsymbol{V}_{\mathbf{1}}=\left( \begin{matrix} K_{n_{1}n_{1}} & K_{n_{1}n_{2}}\otimes\begin{matrix} [1 & 1] \end{matrix} \\ K_{n_{2}n_{1}}\otimes\left[ \begin{matrix} 1 \\ 1 \end{matrix} \right] & K_{n_{2}n_{2}}\otimes\left[ \begin{matrix} 1 & 1 \\ 1 & 1 \end{matrix} \right] \end{matrix} \right)$, $\boldsymbol{V}_{\mathbf{2}}=\left( \begin{matrix} \boldsymbol{I}_{n_{1}\times n_{1}} & \mathbf{0}_{n_{1}\times n_{2}} \\ \mathbf{0}_{n_{2}\times n_{1}} & \boldsymbol{I}_{n_{2}\times n_{2}}\otimes[\begin{matrix} 1 & 1 \\ 1 & 1 \end{matrix}] \end{matrix} \right)$,$\boldsymbol{V}_{\mathbf{3}}=\left( \begin{matrix} \frac{1}{2}\boldsymbol{I}_{n_{1}\times n_{1}} & \mathbf{0}_{n_{1}\times n_{2}} \\ \mathbf{0}_{n_{2}\times n_{1}} & \boldsymbol{I}_{n_{2}\times n_{2}}\otimes[\begin{matrix} 1 & 0 \\ 0 & 1 \end{matrix}] \end{matrix} \right)$.

We treat the approximated quasi-likelihood function $\tilde{ql}$ in equation (10) as the target function. And we obtain estimates for $(\boldsymbol{b}, \mu, \beta)$ and (${\sigma_{g}^{2},\sigma}_{u}^{2},\sigma_{e}^{2}$) alternately from equation (10).

Specifically, we first obtain estimates for $(\boldsymbol{b}, \mu, \beta)$ conditional on the current estimates of (${\sigma_{g}^{2},\sigma}_{u}^{2},\sigma_{e}^{2}$). To do so, following [1], we assume that the iterative weights vary slowly with respect to the conditional mean; that is

$$\frac{\partial\mathbf{D}}{\partial\lambda_{i}}\approx0.$$

We then obtain the first order derivatives with respect to either $(\mu, \beta)$ or $\boldsymbol{b}$, and set the two first order derivatives to zero; that is

|  | $\left( \mathbf{1},\boldsymbol{x} \right)^{T}\mathbf{D}\Delta\left( \boldsymbol{y}-\boldsymbol{\lambda r} \right)=\mathbf{0},$ | $(11)$ |
| --- | --- | --- |
|  | $\mathbf{D}\Delta\left( \boldsymbol{y}-\boldsymbol{\lambda r} \right)-\mathbf{V}^{-\mathbf{1}}\boldsymbol{b}=\mathbf{0},$ | $(12)$ |

where $\boldsymbol{\lambda}=(\lambda_{1}, \cdots,\lambda_{n_{1}+2n_{2}})$ and $\Delta=diag\left\{ g'\left( \lambda_{i} \right) \right\}$.

We now define the pseudo-data

|  | $\tilde{y_{i}}=\eta_{i}+g^{'}\left( \lambda_{i} \right)\left( y_{i}-\lambda_{i}r_{i} \right),$ | $(13)$ |
| --- | --- | --- |

and our equation (11) becomes

|  | $\boldsymbol{b}=\mathbf{V}\mathbf{H}^{-1}\left[ \tilde{\boldsymbol{y}}-\left( \mathbf{1},\boldsymbol{x} \right)\left( \begin{matrix} \mu\\ \boldsymbol{\beta} \end{matrix} \right) \right],$ | $(14)$ |
| --- | --- | --- |

where $\mathbf{H}=\mathbf{D}^{-1}+\mathbf{V}$. Substituting equation (14) into equation (12), we can obtain the estimates

|  | $\left( \begin{matrix} \hat{\mu} \\ \hat{\boldsymbol{\beta}} \end{matrix} \right)= \left[ \left( \mathbf{1},\boldsymbol{x} \right)^{T}\mathbf{H}^{-1}\left( \mathbf{1},\boldsymbol{x} \right) \right]^{-1}\left( \mathbf{1},\boldsymbol{x} \right)^{T}\boldsymbol{H}^{-1}\tilde{\boldsymbol{y}},$ | $(15)$ |
| --- | --- | --- |

and

|  | $\hat{\mathbf{b}}=\mathbf{V}\mathbf{H}^{-1}\left[ \tilde{\boldsymbol{y}}-\left( \mathbf{1},\boldsymbol{x} \right)\left( \begin{matrix} \hat{\mu} \\ \hat{\boldsymbol{\beta}} \end{matrix} \right) \right],$ | $(16)$ |
| --- | --- | --- |

Both are conditional on the variance component estimates (${\sigma_{g}^{2},\sigma}_{u}^{2},\sigma_{e}^{2}$).

Next, we obtain estimates for the variance components (${\sigma_{g}^{2},\sigma}_{u}^{2},\sigma_{e}^{2}$) conditional on the current estimates of $(\boldsymbol{b}, \mu, \beta)$*.* Here we first define the variance components $\tau_{1}=\sigma_{g}^{2}$,$\tau_{2}=\sigma_{u}^{2}$ and $\tau_{3}=\sigma_{g}^{2}$. We then integrate out the fixed effects $\mu$ and $\beta$ in equation (10) to obtain the restricted likelihood function as

$$\tilde{ql}_{R}\left( \tau_{1},\tau_{2},\tau_{3} \right)=c_{R}-\frac{1}{2}\log\left| \mathbf{H} \right|-\frac{1}{2}\log\left| \left( \mathbf{1},\boldsymbol{x} \right)^{T}\mathbf{H}^{-1}\left( \mathbf{1},\boldsymbol{x} \right) \right|-\frac{1}{2}{\tilde{\boldsymbol{y}}}^{T}\mathbf{P}\tilde{\boldsymbol{y}},$$

where $\mathbf{P}= \mathbf{H}^{-1}-\mathbf{H}^{-1}\left( \mathbf{1},\boldsymbol{x} \right)^{T}\left( \left( \mathbf{1},\boldsymbol{x} \right)^{T}\mathbf{H}^{-1}\left( \mathbf{1},\boldsymbol{x} \right) \right)^{-1}\left( \mathbf{1},\boldsymbol{x} \right)\mathbf{H}^{-1}$, and $c_{R}$ is a constant. We use the AI algorithm to obtain variance component estimates. In particular, we obtain the first derivatives as

$$\frac{\partial\tilde{ql}_{R}\left( {\sigma_{g}^{2},\sigma}_{u}^{2},\sigma_{e}^{2} \right)}{\partial\tau_{i}}=\frac{1}{2}\left\{ {\tilde{\boldsymbol{y}}}^{T}\mathbf{P}\boldsymbol{V}_{\boldsymbol{i}}\mathbf{P}\tilde{\boldsymbol{y}}-tr\left( \mathbf{P}\boldsymbol{V}_{\boldsymbol{i}} \right) \right\}, i\in\left\{ 1,2,3 \right\},$$

and the second derivatives as

$$\frac{\partial^{2}\tilde{ql}_{R}\left( {\sigma_{g}^{2},\sigma}_{u}^{2},\sigma_{e}^{2} \right)}{\partial\tau_{i}\partial\tau_{j}}=\frac{1}{2}tr\left( \boldsymbol{V}_{\boldsymbol{i}}\mathbf{P}\boldsymbol{V}_{\boldsymbol{j}}\mathbf{P} \right)-{\tilde{\boldsymbol{y}}}^{T}\mathbf{P}\boldsymbol{V}_{\boldsymbol{i}}\mathbf{P}\boldsymbol{V}_{\boldsymbol{j}}\mathbf{P}\tilde{\boldsymbol{y}}; i,j\in\{1,2,3\}$$

The second derivatives constitute the observed information matrix. Because the elements in the expected information matrix are

$$E\left[ \frac{\partial^{2}\tilde{ql}_{R}\left( {\sigma_{g}^{2},\sigma}_{u}^{2},\sigma_{e}^{2} \right)}{\partial\tau_{i}\partial\tau_{j}} \right]=-\frac{1}{2}tr\left( \boldsymbol{V}_{\boldsymbol{i}}\mathbf{P}\boldsymbol{V}_{\boldsymbol{j}}\mathbf{P} \right); i,j\in\left\{ 1,2,3 \right\},$$

we can obtain the average information (AI) matrix as an average of the above two matrices; that is

$$\mathbf{AI=}\left[ \begin{matrix} {\tilde{\boldsymbol{y}}}^{T}\mathbf{P}\boldsymbol{V}_{\boldsymbol{1}}\mathbf{P}\boldsymbol{V}_{\boldsymbol{1}}\mathbf{P}\tilde{\boldsymbol{y}} & {\tilde{\boldsymbol{y}}}^{T}\mathbf{P}\boldsymbol{V}_{\boldsymbol{1}}\mathbf{P}\boldsymbol{V}_{\boldsymbol{2}}\mathbf{P}\tilde{\boldsymbol{y}} & {\tilde{\boldsymbol{y}}}^{T}\mathbf{P}\boldsymbol{V}_{\boldsymbol{1}}\mathbf{P}\boldsymbol{V}_{\boldsymbol{3}}\mathbf{P}\tilde{\boldsymbol{y}} \\ {\tilde{\boldsymbol{y}}}^{T}\mathbf{P}\boldsymbol{V}_{\boldsymbol{2}}\mathbf{P}\boldsymbol{V}_{\boldsymbol{1}}\mathbf{P}\tilde{\boldsymbol{y}} & {\tilde{\boldsymbol{y}}}^{T}\mathbf{P}\boldsymbol{V}_{\boldsymbol{2}}\mathbf{P}\boldsymbol{V}_{\boldsymbol{2}}\mathbf{P}\tilde{\boldsymbol{y}} & {\tilde{\boldsymbol{y}}}^{T}\mathbf{P}\boldsymbol{V}_{\boldsymbol{2}}\mathbf{P}\boldsymbol{V}_{\boldsymbol{3}}\mathbf{P}\tilde{\boldsymbol{y}} \\ {\tilde{\boldsymbol{y}}}^{T}\mathbf{P}\boldsymbol{V}_{\boldsymbol{3}}\mathbf{P}\boldsymbol{V}_{\boldsymbol{1}}\mathbf{P}\tilde{\boldsymbol{y}} & {\tilde{\boldsymbol{y}}}^{T}\mathbf{P}\boldsymbol{V}_{\boldsymbol{3}}\mathbf{P}\boldsymbol{V}_{\boldsymbol{2}}\mathbf{P}\tilde{\boldsymbol{y}} & {\tilde{\boldsymbol{y}}}^{T}\mathbf{P}\boldsymbol{V}_{\boldsymbol{3}}\mathbf{P}\boldsymbol{V}_{\boldsymbol{3}}\mathbf{P}\tilde{\boldsymbol{y}} \end{matrix} \right]\boldsymbol{.}$$

With the first and second order derivatives, we can perform Newton-Raphson update with the AI algorithm and obtain estimates for $\left( {\sigma_{g}^{2},\sigma}_{u}^{2},\sigma_{e}^{2} \right)$.

As a summary, we implement the PQL algorithm that consists of the following steps:

1. Initialize the parameters, $\mu^{\left( 0 \right)}\boldsymbol{,}\beta^{\left( 0 \right)}$,$\boldsymbol{\tau}^{\left( 0 \right)}=\left( \tau_{1}^{\left( 0 \right)},\tau_{2}^{\left( 0 \right)},\tau_{3}^{\left( 0 \right)} \right)^{T},$ and obtain the pseudo-data ${\tilde{\boldsymbol{y}}}^{\left( 0 \right)}$ as in equation (12). Set $t=1$.
2. Update $\boldsymbol{\tau}^{\left( t \right)}\boldsymbol{=}\boldsymbol{\tau}^{\left( t-1 \right)}\boldsymbol{+}\mathbf{AI}^{-1}\left( \frac{\partial{ql}_{R}\left( {\sigma_{g}^{2},\sigma}_{u}^{2},\sigma_{e}^{2} \right)}{\partial\boldsymbol{\tau}} \right)$;
3. Update $\mu^{\left( t \right)}\boldsymbol{,}\beta^{\left( t \right)}$ and $\boldsymbol{b}^{\left( t \right)}$ with $\boldsymbol{\tau}^{\left( t \right)}$ and ${\tilde{\boldsymbol{y}}}^{\left( t-1 \right)}$ as in equations (15) and (16);
4. Update ${\tilde{\boldsymbol{y}}}^{\left( t \right)}$ using the $\mu^{\left( t \right)}, \beta^{\left( t \right)}$ and $\boldsymbol{b}^{\left( t \right)}$ as in equation (13);
5. Set $t=t+1$, and repeat steps 2-4 until convergence.

Once we obtain parameter estimates $(\mu,\beta,{\sigma_{g}^{2},\sigma}_{u}^{2},\sigma_{e}^{2})$, we can construct the Wald test based on

$\hat{\beta}=\left( \boldsymbol{x}^{T}\mathbf{P}_{\boldsymbol{c}}\boldsymbol{x} \right)^{-1}\boldsymbol{x}^{T}\mathbf{P}_{\boldsymbol{c}}\tilde{\boldsymbol{y}}$ and $\mathrm{Var}\left( \hat{\beta} \right)=\left( \boldsymbol{x}^{T}\mathbf{P}_{\boldsymbol{c}}\boldsymbol{x} \right)^{-1}$,

where $\boldsymbol{P}_{\boldsymbol{c}}= \mathbf{H}^{-1}-\mathbf{H}^{-1}\mathbf{W}^{T}\left( \mathbf{W}^{T}\mathbf{H}^{-1}\mathbf{W}^{T} \right)^{-1}\mathbf{W}^{T}\mathbf{H}^{-1}$.

Because of the approximation using PQL, when sample size is small (<150), we observe an inflation of p-values in the simulations and real data applications (e.g. the genomic control factor equals 1.21 and 1.12 in the two real data sets). We therefore perform an additional step of genomic control to adjust for p-values in a *post hoc* fashion. Specifically, we normalize $\chi^{2}$ statistics (across all SNP-CpG pairs) by dividing max(1, $\lambda_{gc}$) and then use the normalized $\chi^{2}$ statistics to obtain the final p-values.

**3. M-values Calculation**

For methods that use normalized data, we performed “M” value transformation. To do so, we first divided the number of methylated reads by the number of unmethylated reads. Afterwards, we performed a log2-transformation to obtain the final normalized data. The final normalized data is in the form of $\log_{2} \left( \frac{methylated reads+a}{unmethylated reads+a} \right)$, where a small value $a$ = 0.01 is added to avoid zero values inside the log function.

**Supplementary Figures**

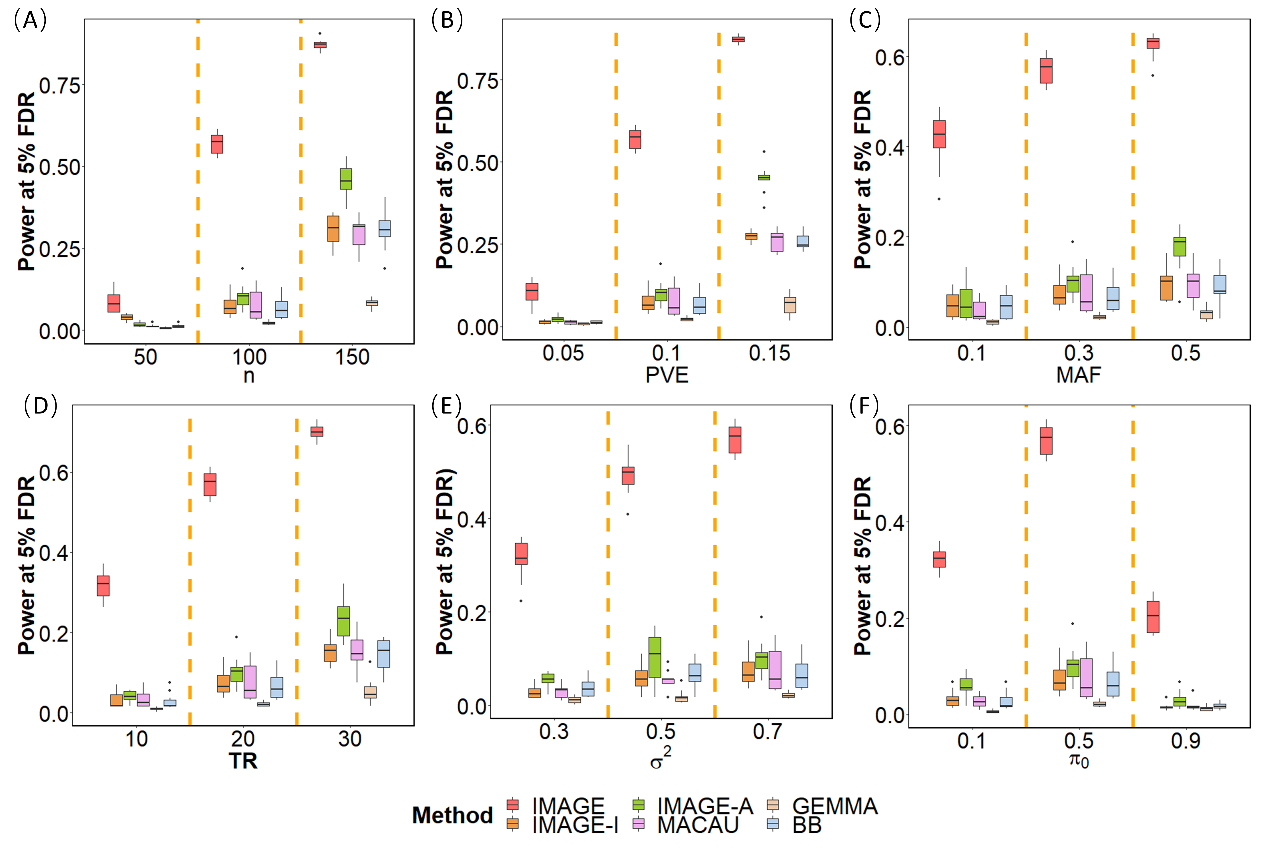

**Supplementary Figure 1**: IMAGE achieves higher power to detect mQTL across various simulation settings. Power is measured as the proportion of true positives detected at a false discovery rate (FDR) of 0.05. Each simulation setting consists of 10 simulation replicates, and each replicate includes 10,000 simulated SNP-CpG pairs, with 1,000 mQTL and 9,000 non-mQTL. (A) We vary $n$ to be either 50, 100, or 150, while maintaining other parameters at the baseline setting. With increasing sample size, all methods gain power. (B) We vary Percentage of Variance Explained by the SNP ($PVE)$ to be either 0.05, 0.1, or 0.15, while maintain other parameters at the baseline setting. With increasing $PVE$, all methods gain power. (C) We vary Minor Allele Frequency ($MAF)$ to be either 0.1, 0.3, or 0.5, while maintaining other parameters at the baseline setting. With increasing $MAF$, all methods gain power. (D) We vary average read counts per site ($TR)$ to be either 10, 20, or 30, while maintaining other parameters at the baseline setting. With increasing $TR$, all methods gain power. (E) We vary over-dispersion variance ($\sigma^{2}$) to be either 0.3,0.5, or 0.7, while maintaining other parameters at the baseline setting. With increasing $\sigma^{2}$, all methods gain power. (F) We vary mean methylation ratio ($\pi_{0}$) to be either 0.1, 0.5, or 0.9, while maintaining other parameters at the baseline setting. The power of all methods is highest for a moderate methylation level $\pi_{0}$, but reduces for hypermethylated and hypomethylated sites. FDR: false discovery rate.

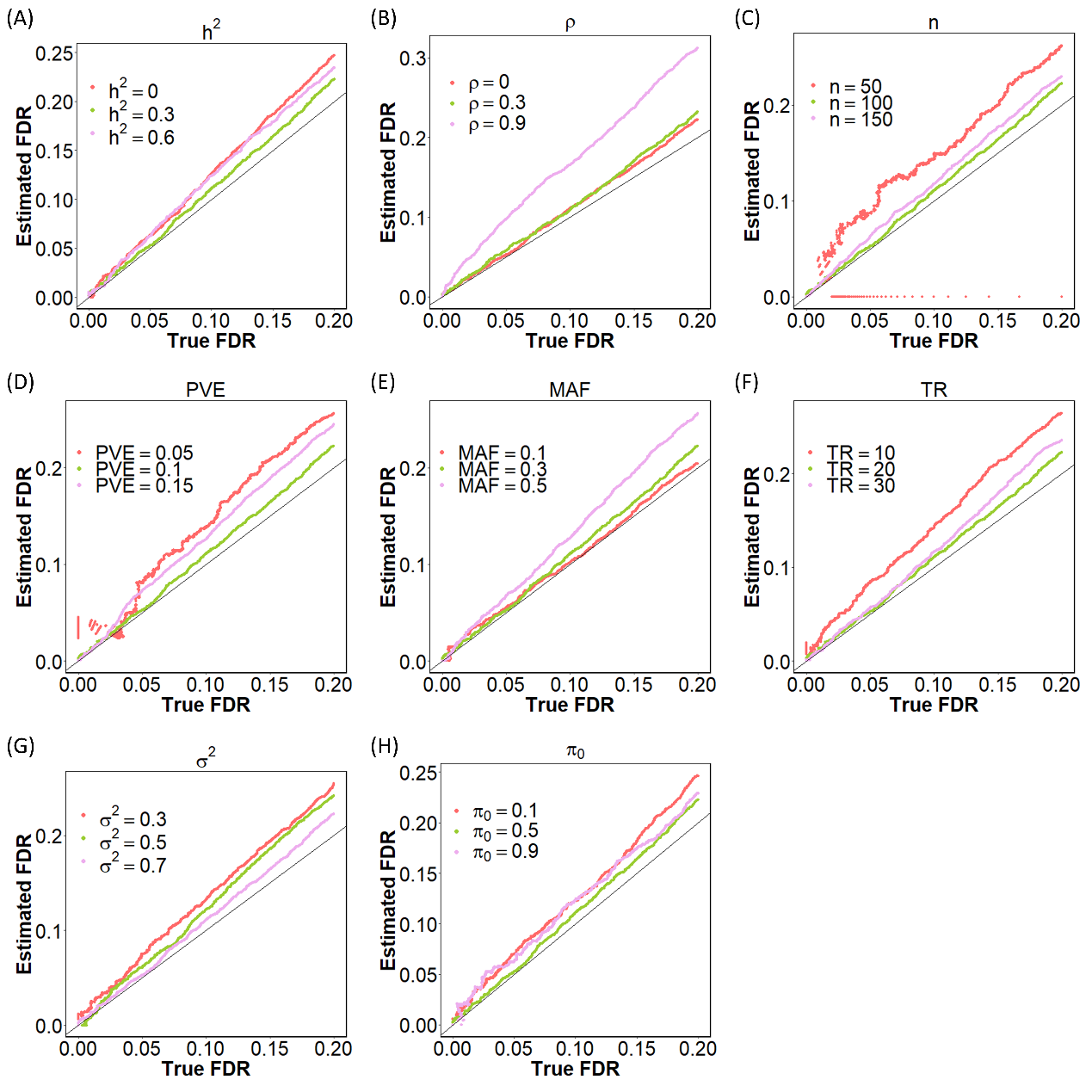

**Supplementary Figure 2**: IMAGE yields reasonably well-calibrated false discovery rate (FDR) estimates across a range of parameter values in simulations. Each simulation setting includes 100,000 simulated SNP-CpG pairs, with 10,000 associated pairs and 90,000 unassociated pairs. In each simulation setting, we performed 10 permutations to estimate the empirical FDR (y-axis), which is plotted against the true FDR (x-axis). In (A), we vary $h^{2}$ to be 0 (red), 0.3 (green), or 0.6 (pink). In (B), we vary $\rho$ to be either 0 (red), 0.3 (green), or 0.9 (pink). In (C), we vary $n$ to be either 50 (red), 100 (green), or 150 (pink). In (D), we vary $\mathrm{PVE}$ to be either 0.05 (red), 0.1 (green), or 0.15 (pink). In (E), we vary $\mathrm{MAF}$ to be either 0.05 (red), 0.3 (green), or 0.5 (pink). In (F), we vary $\mathrm{TR}$ to be either 10 (red), 20 (green), or 30 (pink). In (G), we vary $\sigma^{2}$ to be either 0.3 (red), 0.5 (green), or 0.7 (pink). In (H), we vary $\pi_{0}$ to be either 0.1 (red), 0.3 (green), or 0.5 (pink). In each panel, we maintain the other parameters as in the baseline setting. Note that the upward deviation from the x=y line reflects a conservative estimate of the FDR: thus, IMAGE is not likely to generate more false discoveries than expected at a given FDR threshold, although it will miss a small number of likely true positives.

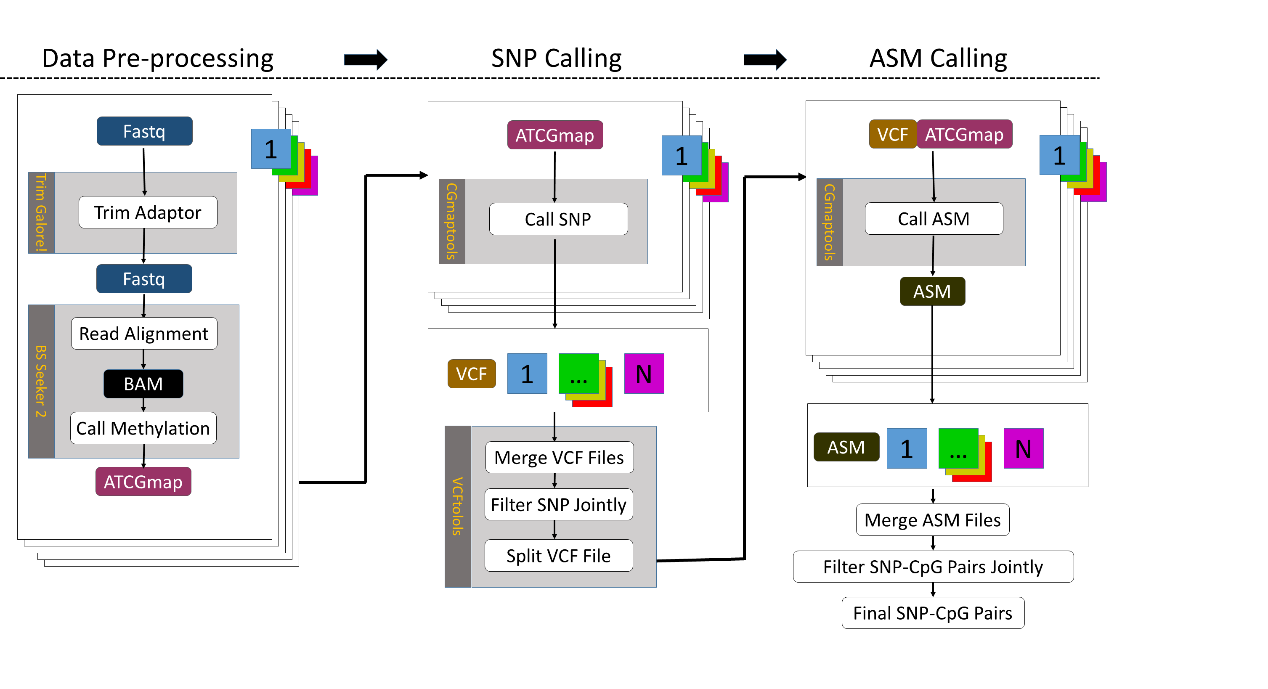

**Supplementary Figure 3**: Schematic of the data processing steps for the baboon and wolf data sets. We processed input files in the fastq format to obtain both SNP information and methylation information, which were paired together and subjected through various filtering steps to yield a final set of SNP-CpG pairs for analysis. Briefly, steps include: (1) trimmed the adaptors in the fastq files using TrimGalore!; (2) Mapped to the baboon reference genome *Panu2.0* or the dog reference genome *canFam3.1* using BSseeker2; (3) Called SNPs using CGmaptools with the BayesWC strategy; (4) Merged and filtered VCF files using VCFtools; (5) Called allele-specific methylation using CGmaptools; (6) Merged and filtered ASM data to yield a final set of SNP-CpG pairs. In the above figure, Fastq, BAM, ATCGmap, VCF, and ASM refer to different file formats.

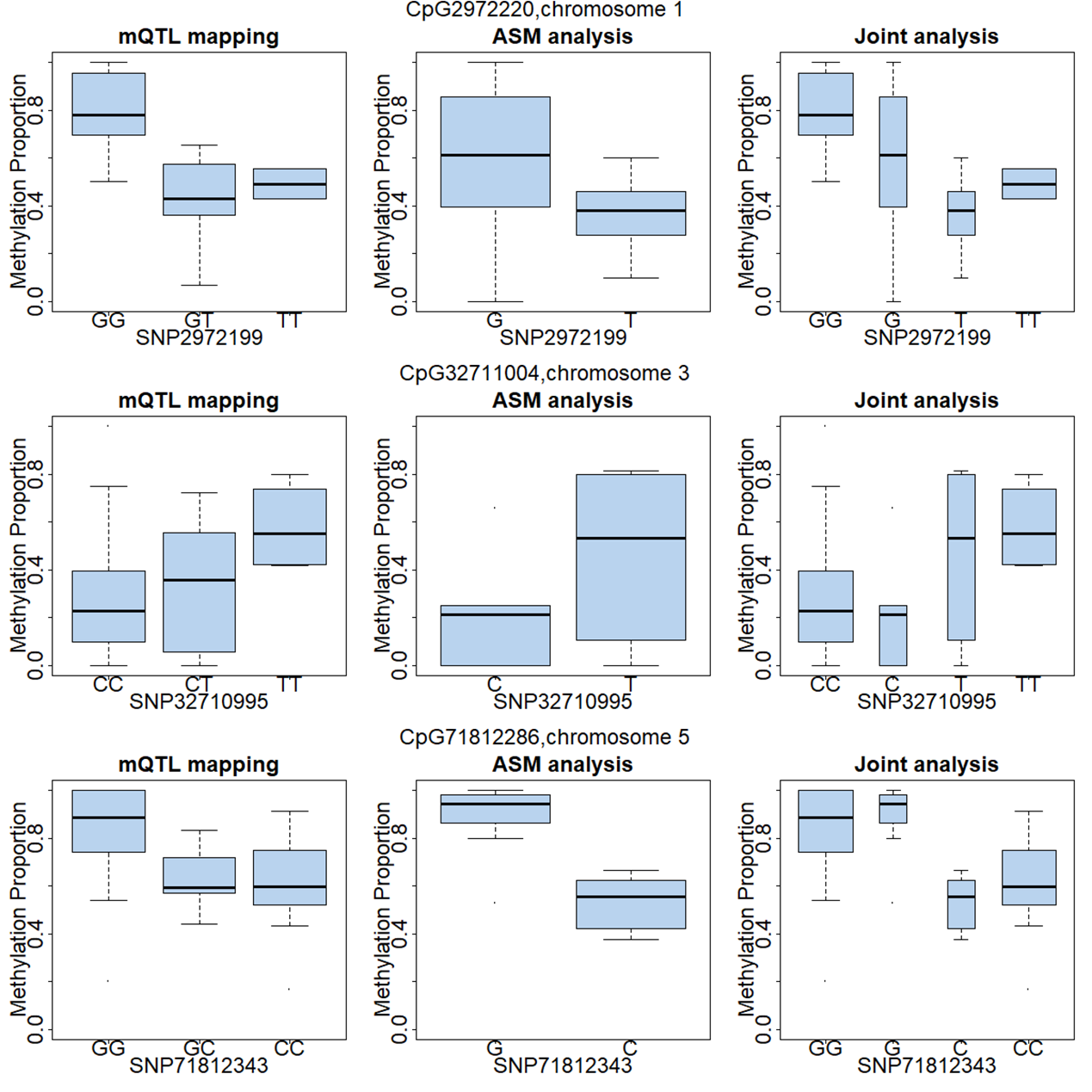

**Supplementary Figure 4**: Examples of associated SNP-CpG pairs detected by IMAGE in the baboon data. Each row contains one SNP-CpG pair. Boxplots on the three left panels show individual-level methylation information used by mQTL mapping methods. Boxplots on the three middle panels show the allele-specific methylation information used by ASM analysis methods. Boxplots on the three right panels show both individual-level methylation information and allele-specific methylation information used by IMAGE.

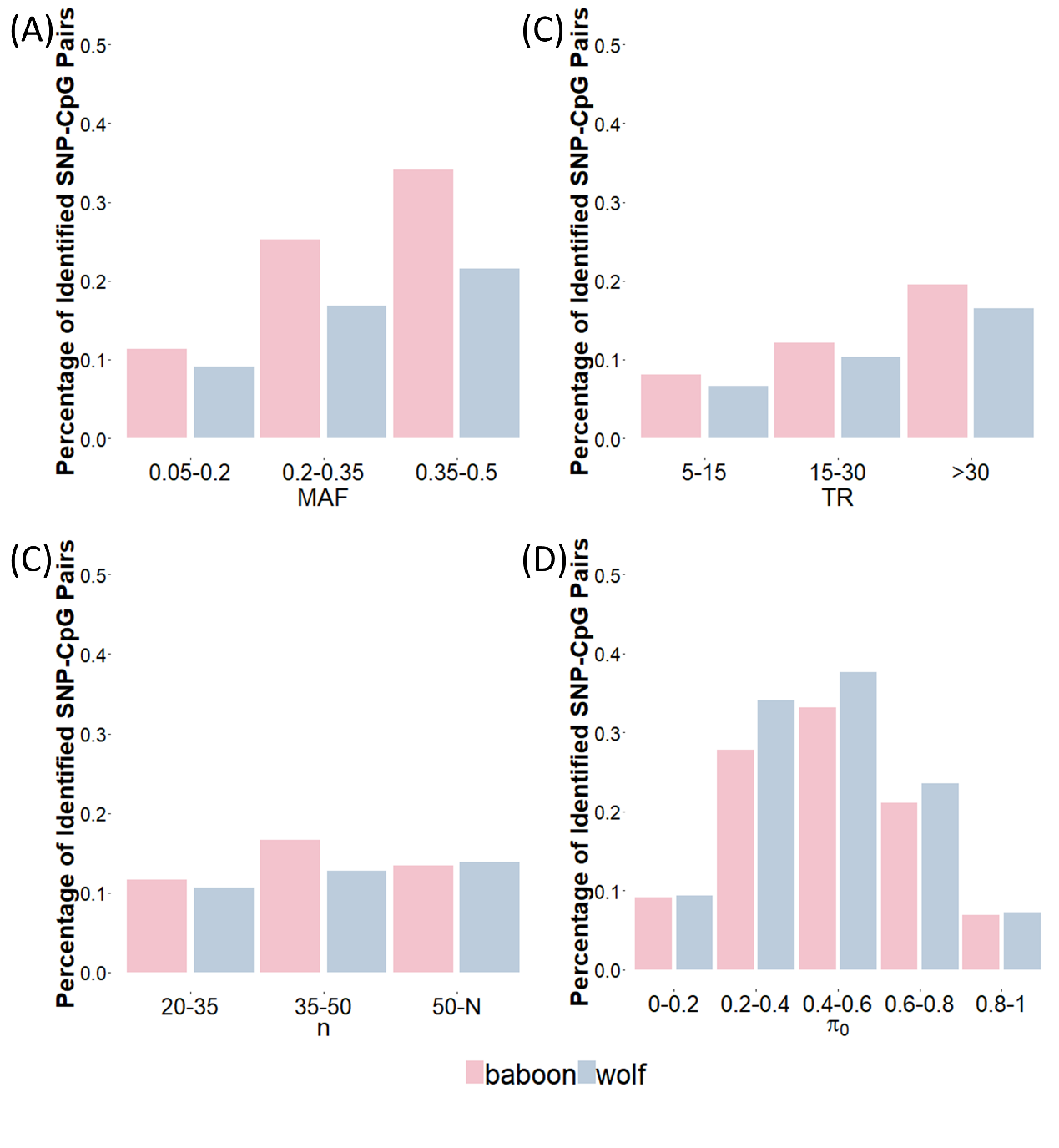

**Supplementary Figure 5**: Number of associated SNP-CpG pairs identified by IMAGE for sites stratified by MAF, TF, n, and methylation level in the baboon (pink) and wolf data (blue). IMAGE achieves higher power with increasing MAF (A), increasing read depth (B), increasing sample size (C), and intermediate methylation levels (D). In (C), N represents the sample size (67 for the baboon data and 63 for the wolf data).

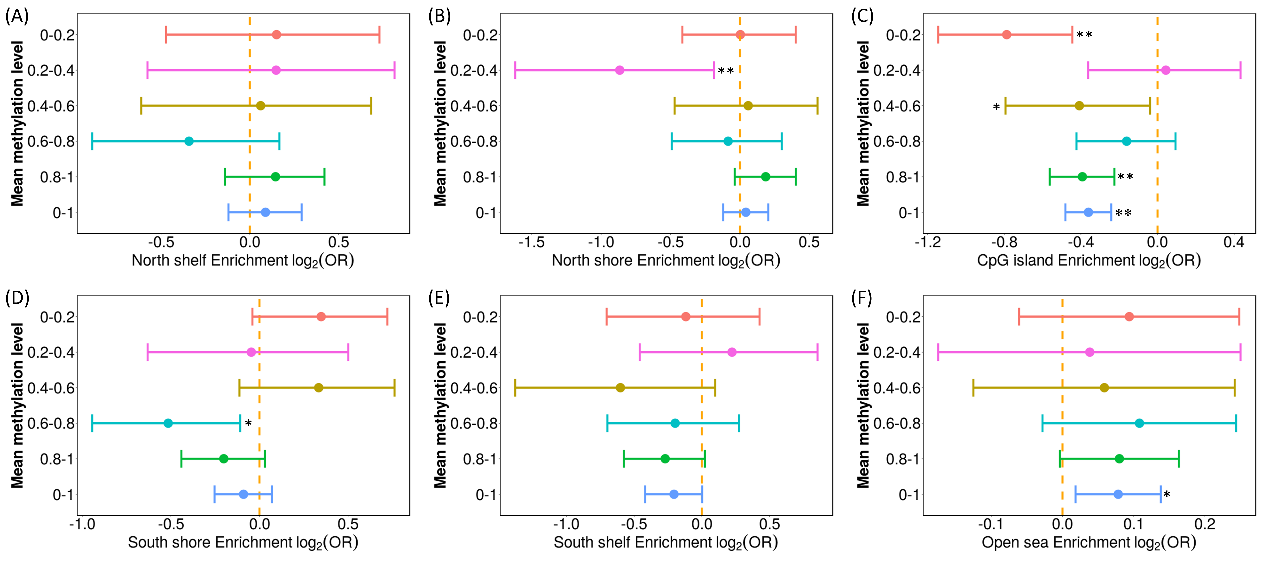

**Supplementary Figure 6**: CpG sites with IMAGE-identified mQTL, regardless of methylation levels, are enriched in open sea regions and depleted in CpG islands in the baboon data. log2 odds ratio (x-axis) of detecting associated SNP-CpG pairs, together with the 95% CI, is computed for CpG sites residing in different annotated genomic regions, further stratified with regard to methylation levels (y-axis). CpG sites with IMAGE-identified mQTL are enriched in open sea regions and depleted in CpG islands. *P<0.05. **P<0.01.

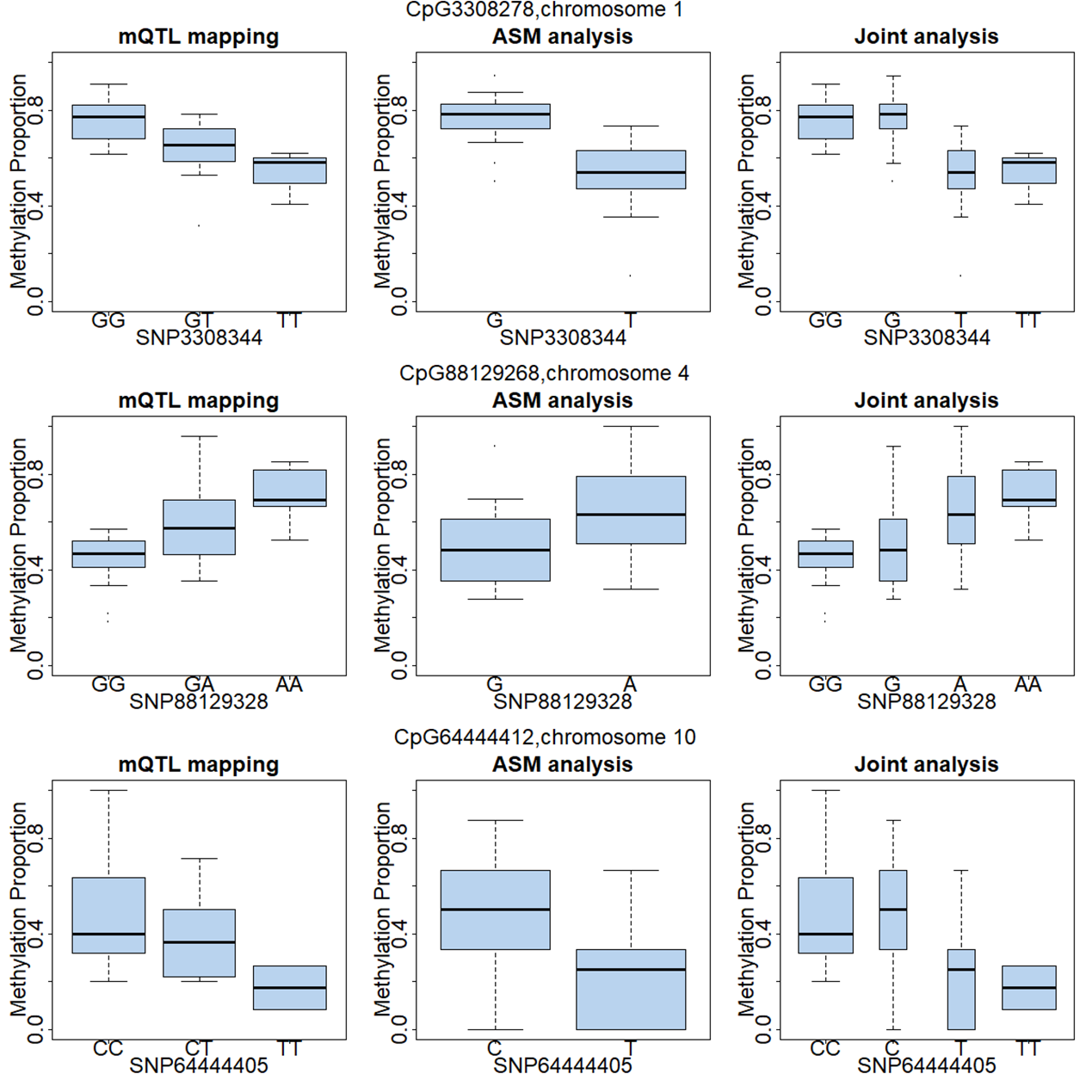

**Supplementary Figure 7**: Examples of associated SNP-CpG pairs detected by IMAGE in the wolf data. Each row contains one SNP-CpG pair. Boxplots on the three left panels show individual-level methylation information used by mQTL mapping methods. Boxplots on the three middle panels show the allele-specific methylation information used by ASM analysis methods. Boxplots on the three right panels show both individual-level methylation information and allele-specific methylation information used by IMAGE.

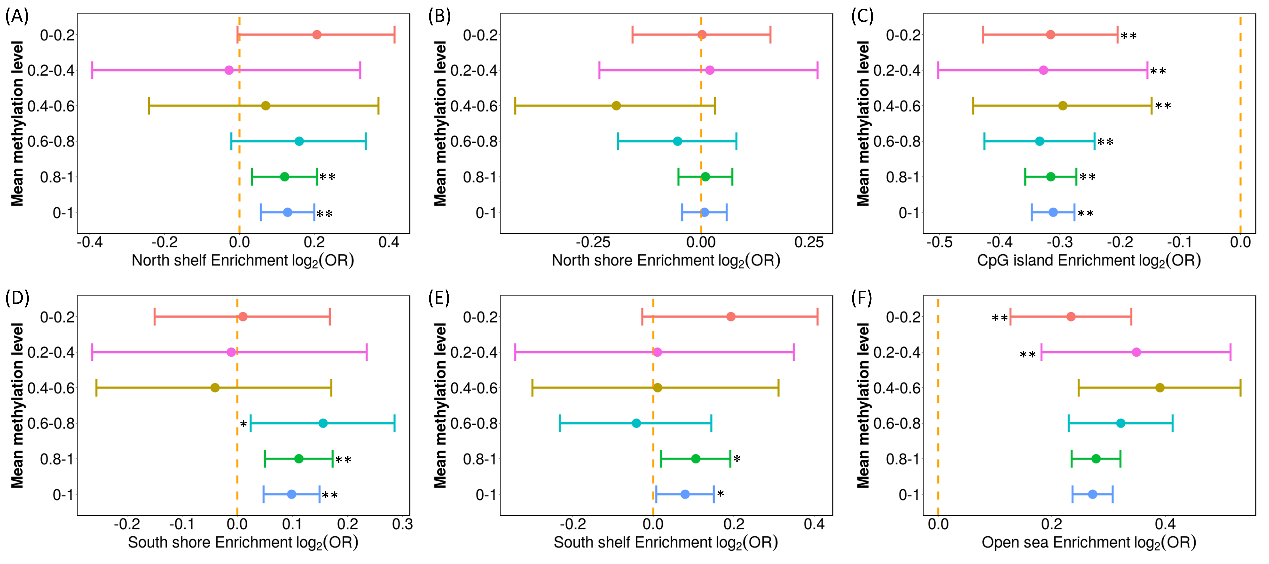
**Supplementary Figure 8**: CpG sites with IMAGE-identified mQTL, regardless of methylation levels, are enriched in open sea regions and depleted in CpG islands in the wolf data. log2 odds ratio (x-axis) of detecting associated SNP-CpG pairs, together with the 95% CI, is computed for CpG sites residing in different annotated genomic regions, further stratified with regard to methylation levels (y-axis). CpG sites with IMAGE-identified mQTL are enriched in open sea regions and depleted in CpG islands. *P<0.05. **P<0.01.

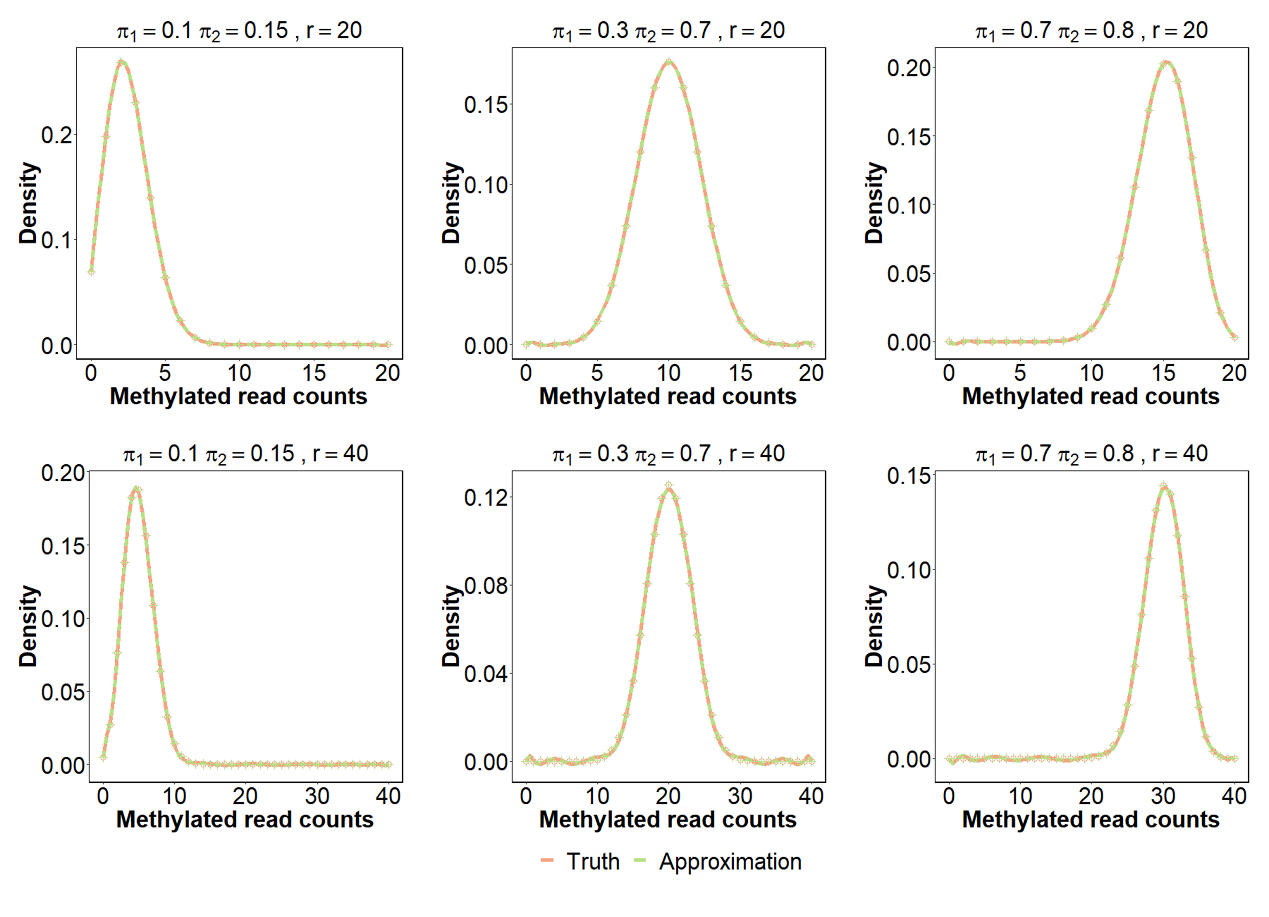

**Supplementary Figure 9**: The binomial approximation used in IMAGE through matching moments approximates the true underlying density in a variety of parameter settings. The parameters examined include $\pi_{1}, \pi_{2},r$ (on top of each panel) and the methylated read counts (on x-axis). The approximation works well across a range of settings.

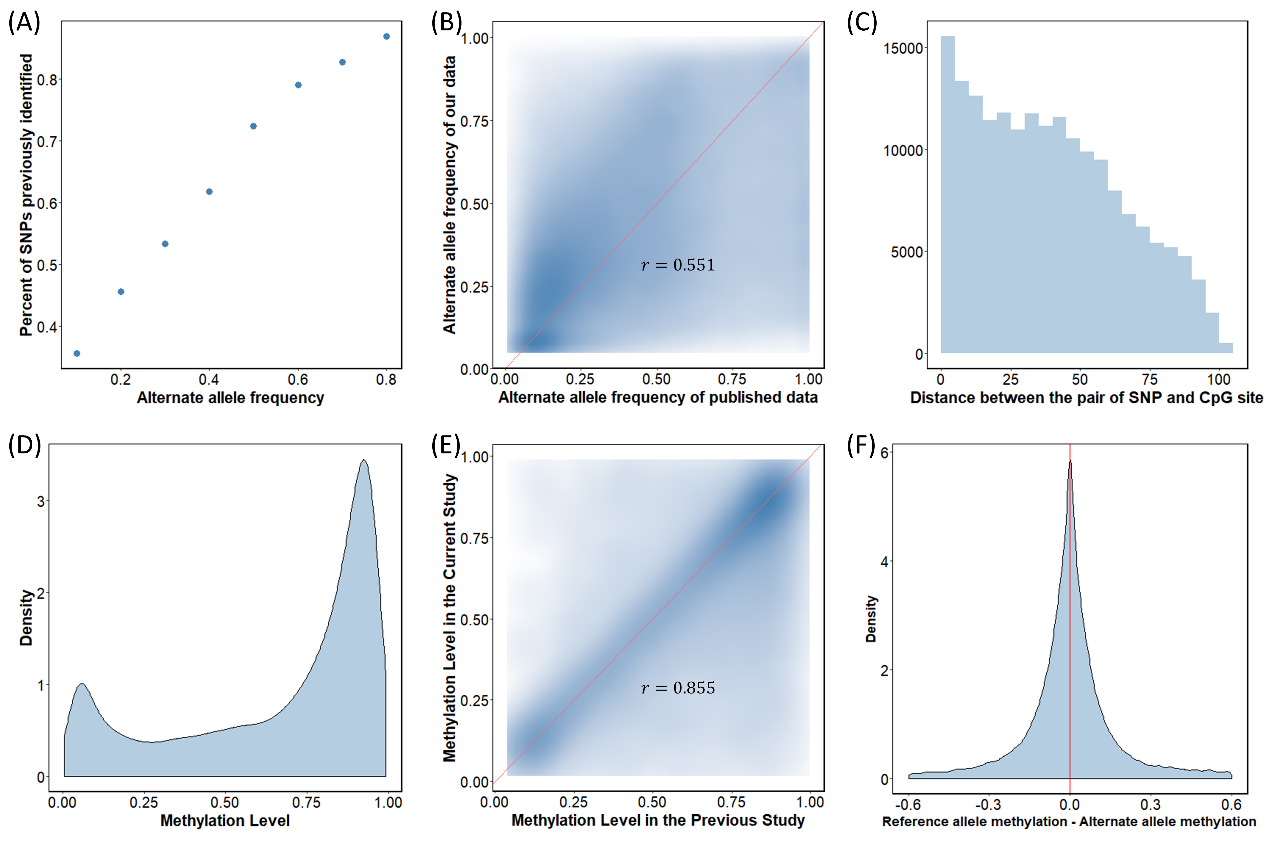

**Supplementary Figure 10**: Quality control in the baboon data. (A) Variants called in our dataset overlap with variants identified through DNA sequencing in a separate sample in East African baboons from a previous study [5]. The overlap proportion (y-axis) increases with increasing alternate allele frequency (x-axis). (B) Among the variants identified in both our data and the previous DNA sequencing study, the alternate allele frequency in our data (y-axis) is highly correlated with the alternate allele frequency in the DNA sequencing study (x-axis). (C) Distribution of the distance between SNP and CpG site across all analyzed SNP-CpG pairs in the baboon data. Because the data were generated using 100 bp reads, 100 bp is the maximum possible distance in our data set. (D) Density plot for the distribution of methylation levels across analyzed CpG sites in the baboon data. The methylation level is measured as the proportion of methylated reads. A bimodal pattern, with both hypomethylated and hypermethylated CpG sites, is clearly visible. (E) The methylation level in our processing pipeline is highly correlated with that measured through in a different processing pipeline in a previous study using the same baboon data [6]. (F) Density plot of the difference in methylation level between the reference allele and the alternate allele across CpG sites in the baboon data. The difference is symmetric, suggesting no apparent reference mapping bias in the data.

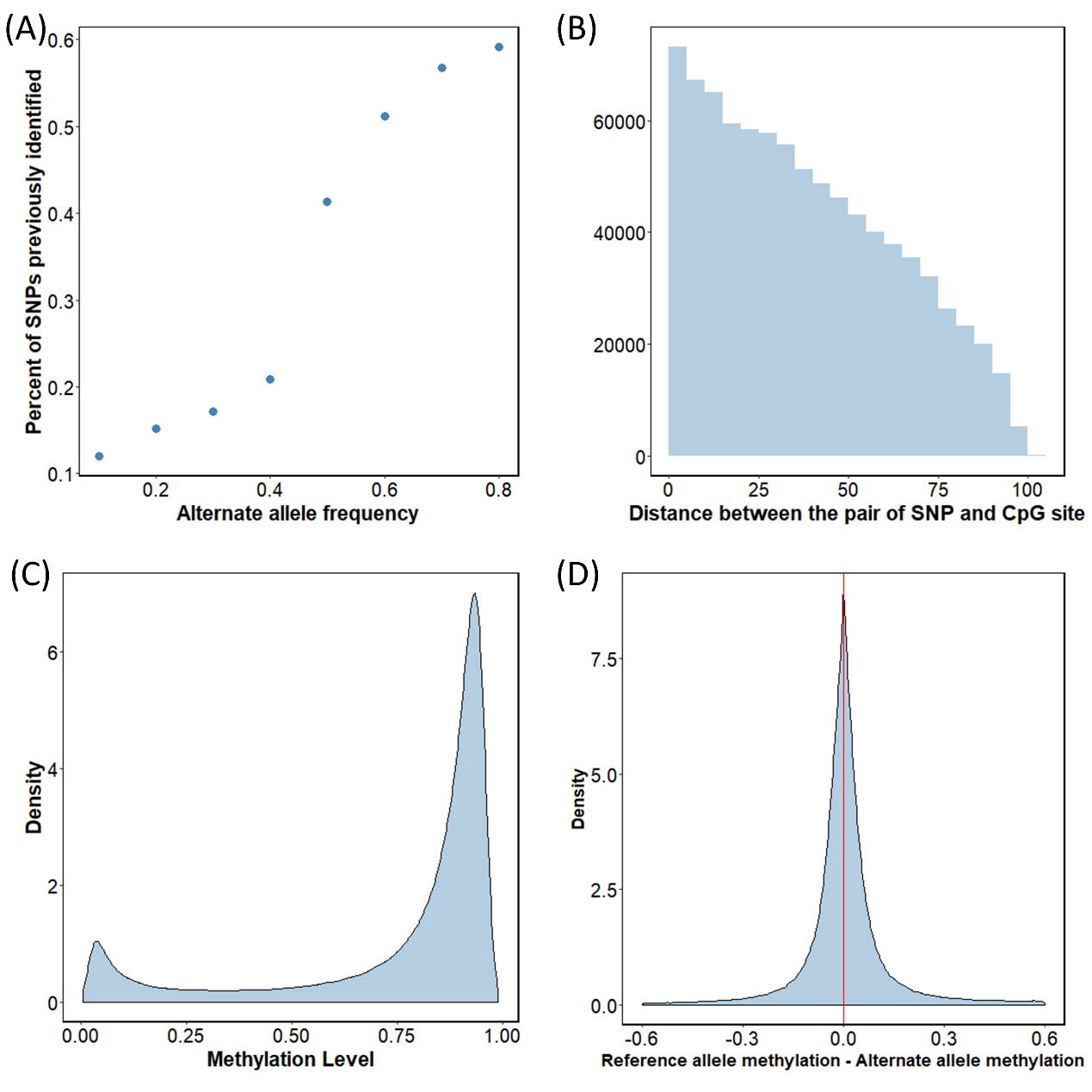

**Supplementary Figure 11**: Quality control in the wolf data. (A) Common SNP variants in our wolf dataset are likely to overlap with the variants from the current Ensembl release for dogs. Increasing the alternate allele frequency increases the proportion of SNPs that were previously identified. (B) The histogram of distance between SNPs and CpG sites of the wolf data. Because the data were generated using 100 bp reads, 100 bp is the maximum possible distance in our data set. (C) Density plot of the distribution of the average methylation levels in wolf data. As expected for RRBS data, the data contain a large number of hypomethylated and hypermethylated CpG sites. (D) Density plot of the difference in methylation level between reference and alternate alleles in wolf data. For most CpG sites, there is little or no difference in methylation level between the reference and the alternate alleles.

**Supplementary Tables**

**Supplementary Table 1**: Enrichment of CpG sites in different genomic regions for CpG sites with mQTL versus CpG sites without mQTL in the baboon data. Percentage of CpG sites in different genomic regions (open sea; north/south shelf; north/south shore; CpG island) for CpG sites with mQTL (3^rd^ column) versus CpG sites without mQTL (4^th^ column). A fold enrichment is computed as the odds ratio (5^th^ column) and tested through a Fisher’s exact test (6^th^ column). mQTL are detected by different methods (IMAGE, IMAGE-I, IMAGE-A, MACAU, GEMMA, BB) based on a permutation-based empirical false discovery rate of 0.05. Bold font highlights significant enrichment (p-value < 0.05).

|  |  | CpG sites with mQTL | CpG sites without mQTL | Fold enrichment | p-value |
| --- | --- | --- | --- | --- | --- |
| IMAGE | North shelf | 3.70% | 3.49% | 1.063 | 0.4013 |
|  | North shore | 6.00% | 5.80% | 1.034 | 0.5474 |
|  | **CpG island** | **11.16%** | **14.33%** | **0.7788** | $\boldsymbol{1.056\times}\mathbf{10}^{\mathbf{-9}}$ |
|  | South shore | 5.97% | 6.35% | 0.9396 | 0.2803 |
|  | South shelf | 3.43% | 3.95% | 0.8672 | 0.0547 |
|  | **Open sea** | **69.74%** | **66.08%** | **1.0554** | $\mathbf{0.0106}$ |
| IMAGE-I | North shelf | 3.08% | 3.57% | 0.8623 | 0.1608 |
|  | North shore | 6.30% | 5.81% | 1.0858 | 0.2564 |
|  | **CpG island** | **11.43%** | **14.17%** | **0.8067** | $\boldsymbol{7.415\times}\mathbf{10}^{\mathbf{-5}}$ |
|  | **South shore** | **5.28%** | **6.36%** | **0.8296** | **0.0184** |
|  | South shelf | 4.43% | 3.87% | 1.144 | 0.1303 |
|  | Open sea | 69.48% | 66.22% | 1.049 | 0.08541 |
| IMAGE-A | North shelf | 3.58% | 3.44% | 1.041 | 0.6375 |
|  | North shore | 6.14% | 5.79% | 1.060 | 0.3865 |
|  | **CpG island** | **12.94%** | **14.40%** | **0.8986** | **0.0202** |
|  | South shore | 6.16% | 6.67% | 0.9235 | 0.2187 |
|  | South shelf | 3.85% | 4.09% | 0.9413 | 0.4749 |
|  | Open sea | 67.33% | 65.60% | 1.026 | 0.2981 |
| MACAU | North shelf | 3.25% | 3.53% | 0.9207 | 0.4965 |
|  | North shore | 6.16% | 5.85% | 1.0530 | 0.5140 |
|  | **CpG island** | **11.63%** | **14.20%** | **0.8190** | $\boldsymbol{6.542\times}\mathbf{10}^{\mathbf{-4}}$ |
|  | South shore | 5.88% | 6.35% | 0.9260 | 0.3635 |
|  | South shelf | 4.46% | 3.86% | 1.1554 | 0.1246 |
|  | Open sea | 68.62% | 66.21% | 1.0364 | 0.2379 |
| GEMMA | North shelf | 2.89% | 3.55% | 0.8141 | 0.0942 |
|  | North shore | 6.57% | 5.83% | 1.1269 | 0.1524 |
|  | **CpG island** | **11.95%** | **14.15%** | **0.8445** | $\mathbf{0.0062}$ |
|  | South shore | 5.38% | 6.38% | 0.8433 | 0.0578 |
|  | South shelf | 4.63% | 3.86% | 1.1995 | 0.0652 |
|  | Open sea | 68.59% | 66.23% | 1.0356 | 0.2804 |
| BB | North shelf | 3.18% | 3.56% | 0.8933 | 0.3137 |
|  | North shore | 6.35% | 5.82% | 1.0910 | 0.2533 |
|  | **CpG island** | **10.77%** | **14.13%** | **0.7622** | $\boldsymbol{2.434\times}\mathbf{10}^{\mathbf{-6}}$ |
|  | **South shore** | **5.23%** | **6.38%** | **0.8197** | **0.0158** |
|  | South shelf | 4.33% | 3.84% | 1.1276 | 0.1954 |
|  | Open sea | 70.14% | 66.27% | 1.0584 | 0.0514 |

**Supplementary Table 2**: Proportion of CpG sites directly disrupted by the SNP. The proportion of CpG sites that are directly disrupted by the SNP is higher for mQTL pairs versus non-mQTL pairs in the baboon data. Data are sorted in rows based on different association methods (IMAGE, IMAGE-I, IMAGE-A, MACAU, GEMMA, BB). Note that MethlQTL-A was applied to analyze 38,250 SNP-CpG pairs while the other methods were applied to analyze all 49,196 SNP-CpG pairs. P-values are calculated using Fisher’s exact tests.

|  | CpG sites with mQTL | CpG sites without mQTL | P-values |
| --- | --- | --- | --- |
| IMAGE | 3.72% | 0.43% | $\text{<2.2×}\text{10}^{\text{-16}}$ |
| IMAGE-I | 4.25% | 0.63% | $\text{<2.2×}\text{10}^{\text{-16}}$ |
| IMAGE-A | 1.49% | 0.70% | $\text{3.107×}\text{10}^{\text{-7}}$ |
| MACAU | 5.66% | 0.62% | $\text{<2.2×}\text{10}^{\text{-16}}$ |
| GEMMA | 4.35% | 0.70% | $\text{<2.2×}\text{10}^{\text{-16}}$ |
| BB | 4.55% | 0.63% | $\text{<2.2×}\text{10}^{\text{-16}}$ |

**Supplementary Table 3**: Enrichment of CpG sites in different genomic regions for CpG sites with mQTL versus CpG sites without mQTL in the wolf data. Percentage of CpG sites in different genomic regions (open sea; north/south shelf; north/south shore; CpG island) for CpG sites with mQTL (3^rd^ column) versus CpG sites without mQTL (4^th^ column). A fold enrichment is computed as the odds ratio (5^th^ column) and tested through a Fisher’s exact test (6^th^ column). mQTL are detected by different methods (IMAGE, IMAGE-I, IMAGE-A, MACAU, GEMMA, BB), based on a permutation-based empirical false discovery rate of 0.05. Bold font highlights significant enrichment (p-value < 0.05).

|  |  | CpG sites with mQTL | CpG sites with no mQTL | Fold enrichment | p-values |
| --- | --- | --- | --- | --- | --- |
| IMAGE | **North shelf** | **6.28%** | **5.74%** | **1.094** | $\boldsymbol{4.249\times}\mathbf{10}^{\mathbf{-4}}$ |
|  | North shore | 12.46% | 12.39% | 1.006 | 0.7704 |
|  | **CpG island** | **30.17%** | **37.43%** | **0.8060** | $\boldsymbol{<2.2\times}\mathbf{10}^{\mathbf{-16}}$ |
|  | **South shore** | **13.11%** | **12.24%** | **1.071** | $\boldsymbol{1.45\times}\mathbf{10}^{\mathbf{-4}}$ |
|  | **South shelf** | **6.22%** | **5.88%** | **1.0578** | **0.0296** |
|  | **Open sea** | **31.77%** | **26.31%** | **1.2075** | $\boldsymbol{<2.2\times}\mathbf{10}^{\mathbf{-16}}$ |
| IMAGE-I | **North shelf** | **6.92%** | **5.76%** | **1.2014** | $\boldsymbol{5.74\times}\mathbf{10}^{\mathbf{-5}}$ |
|  | North shore | 12.07% | 12.38% | 0.9750 | 0.4831 |
|  | **CpG island** | **32.42%** | **36.80%** | **0.8810** | $\boldsymbol{2.412\times}\mathbf{10}^{\mathbf{-8}}$ |
|  | South shore | 13.03% | 12.30% | 1.0594 | 0.0848 |
|  | South shelf | 6.40% | 5.90% | 1.0847 | 0.0731 |
|  | **Open sea** | **29.05%** | **26.85%** | **1.0819** | $\boldsymbol{1.07\times}\mathbf{10}^{\mathbf{-3}}$ |
| IMAGE-A | North shelf | 5.82% | 5.89% | 0.9881 | 0.6716 |
|  | **North shore** | **11.64%** | **12.59%** | **0.9245** | $\boldsymbol{2.022\times}\mathbf{10}^{\mathbf{-4}}$ |
|  | CpG island | 35.34% | 35.88% | 0.9849 | 0.2589 |
|  | Sourth shore | 12.44% | 14.04% | 0.8860 | 0.5704 |
|  | Sourth shelf | 5.80% | 5.93% | 0.9949 | 0.4656 |
|  | **Opea sea** | **28.96%** | **24.21%** | **1.1962** | $\boldsymbol{<2.2\times}\mathbf{10}^{\mathbf{-16}}$ |
| MACAU | **North shelf** | **6.91%** | **5.76%** | **1.1997** | $\boldsymbol{3.878\times}\mathbf{10}^{\mathbf{-5}}$ |
|  | North shore | 12.40% | 12.37% | 1.0024 | 0.934 |
|  | **CpG island** | **32.34%** | **36.66%** | **0.8822** | $\boldsymbol{1.295\times}\mathbf{10}^{\mathbf{-8}}$ |
|  | **South shore** | **13.22%** | **12.31%** | **1.0739** | **0.02684** |
|  | **South shelf** | **6.58%** | **5.90%** | **1.1153** | **0.0131** |
|  | **Open sea** | **28.54%** | **27.03%** | **1.0559** | **0.0197** |
| GEMMA | **North shelf** | **6.98%** | **5.77%** | **1.2097** | $\boldsymbol{5.418\times}\mathbf{10}^{\mathbf{-4}}$ |
|  | North shore | 12.51% | 12.36% | 1.0121 | 0.7731 |
|  | **CpG island** | **31.82%** | **36.61%** | **0.8962** | $\boldsymbol{3.437\times}\mathbf{10}^{\mathbf{-7}}$ |
|  | **South shore** | **12.84%** | **12.33%** | **1.0414** | **0.0268** |
|  | South shelf | 6.50% | 5.90% | 1.1017 | 0.0834 |
|  | **Open sea** | **29.35%** | **27.03%** | **1.0858** | $\boldsymbol{4.393\times}\mathbf{10}^{\mathbf{-3}}$ |
| BB | **North shelf** | **6.63%** | **5.76%** | **1.1510** | $\boldsymbol{1.443\times}\mathbf{10}^{\mathbf{-3}}$ |
|  | North shore | 12.33% | 12.37% | 0.9968 | 0.9353 |
|  | **CpG island** | **32.93%** | **36.65%** | **0.8985** | $\boldsymbol{6.143\times}\mathbf{10}^{\mathbf{-7}}$ |
|  | South shore | 13.07% | 12.28% | 1.0643 | 0.0538 |
|  | South shelf | 6.35% | 5.89% | 1.0781 | 0.0880 |
|  | **Open sea** | **28.70%** | **27.05%** | **1.0610** | $\mathbf{0.0101}$ |

**Supplementary Table 4**: Proportion of CpG sites directly disrupted by the SNP. The proportion of CpG sites that are directly disrupted by the SNP is higher in the associated SNP-CpG pairs versus the non-associated SNP-CpG pairs in the wolf data. Data are sorted in rows based on different association methods (IMAGE, IMAGE-I, IMAGE-A, MACAU, GEMMA, BB). Note that IMAGE-A was applied to analyze 236,092 SNP-CpG pairs while the other methods were applied to analyze all 279,223 SNP-CpG pairs. P-values are calculated through Fisher exact test.

|  | CpG sites with mQTL | CpG sites with no mQTL | P-values |
| --- | --- | --- | --- |
| IMAGE | 3.66% | 0.18% | $\text{<2.2×}\text{10}^{\text{-16}}$ |
| IMAGE-I | 8.06% | 0.34% | $\text{<2.2×}\text{10}^{\text{-16}}$ |
| IMAGE-A | 0.83% | 0.57% | $\text{2.14×}\text{10}^{\text{-6}}$ |
| MACAU | 8.91% | 0.33% | $\text{<2.2×}\text{10}^{\text{-16}}$ |
| GEMMA | 7.62% | 0.38% | $\text{<2.2×}\text{10}^{\text{-16}}$ |
| BB | 8.25% | 0.33% | $\text{<2.2×}\text{10}^{\text{-16}}$ |

**Supplementary Table 5**: Type I error control of different methods in the null simulations across three different heritability values. The type I error of different methods (IMAGE, IMAGE-I, IMAGE-A, MACAU, GEMMA, and BB) are shown at significance thresholds ranging from 0.05 to $1\times{10}^{-5}$.

|  | Method | $5\times{10}^{-2}$ | $1\times{10}^{-2}$ | $5\times{10}^{-3}$ | $1\times{10}^{-3}$ | $5\times{10}^{-4}$ | $1\times{10}^{-4}$ | $5\times{10}^{-5}$ | $1\times{10}^{-5}$ |
| --- | --- | --- | --- | --- | --- | --- | --- | --- | --- |
| $h^{2}=0$ | IMAGE | $5.16\times{10}^{-2}$ | 1.14$\times{10}^{-2}$ | 5.92$\times{10}^{-3}$ | 1.35$\times{10}^{-3}$ | 6.2$\times{10}^{-4}$ | 1.8$\times{10}^{-4}$ | 1.2$\times{10}^{-4}$ | 6$\times{10}^{-5}$ |
|  | IMAGE-I | 5.29$\times{10}^{-2}$ | 1.11$\times{10}^{-2}$ | 5.53$\times{10}^{-3}$ | 1.11$\times{10}^{-3}$ | 5.5$\times{10}^{-4}$ | 1.1$\times{10}^{-4}$ | 8$\times{10}^{-5}$ | 0 |
|  | IMAGE-A | 6.28$\times{10}^{-2}$ | 1.61$\times{10}^{-2}$ | 9.15$\times{10}^{-3}$ | 2.56$\times{10}^{-3}$ | 1.59$\times{10}^{-3}$ | 4.7$\times{10}^{-4}$ | 3$\times{10}^{-4}$ | 1.5$\times{10}^{-4}$ |
|  | MACAU | 4.74$\times{10}^{-2}$ | 9.28$\times{10}^{-3}$ | 4.49$\times{10}^{-3}$ | 8.7$\times{10}^{-4}$ | 4.3$\times{10}^{-4}$ | 1.1$\times{10}^{-4}$ | 5$\times{10}^{-5}$ | 0 |
|  | GEMMA | 4.88$\times{10}^{-2}$ | 9.41$\times{10}^{-3}$ | 4.55$\times{10}^{-3}$ | 7.4$\times{10}^{-4}$ | 3.4$\times{10}^{-4}$ | 7$\times{10}^{-5}$ | 5$\times{10}^{-5}$ | 0 |
|  | BB | 5.38$\times{10}^{-2}$ | 1.07$\times{10}^{-2}$ | 5.37$\times{10}^{-3}$ | 9.9$\times{10}^{-4}$ | 4.4$\times{10}^{-4}$ | 1$\times{10}^{-4}$ | 5$\times{10}^{-5}$ | 0 |
| $h^{2}=0.3$ | IMAGE | 5.38$\times{10}^{-2}$ | 1.16$\times{10}^{-2}$ | 6.44$\times{10}^{-3}$ | 1.62$\times{10}^{-3}$ | 9$\times{10}^{-4}$ | 1.9$\times{10}^{-4}$ | 8$\times{10}^{-5}$ | 0 |
|  | IMAGE-I | 5.45$\times{10}^{-2}$ | 1.21$\times{10}^{-2}$ | 6.86$\times{10}^{-3}$ | 1.67$\times{10}^{-3}$ | 1.05$\times{10}^{-3}$ | 3.5$\times{10}^{-4}$ | 1.6$\times{10}^{-4}$ | 4$\times{10}^{-5}$ |
|  | IMAGE-A | 6.23$\times{10}^{-2}$ | 1.54$\times{10}^{-2}$ | 8.64$\times{10}^{-3}$ | 2.55$\times{10}^{-3}$ | 1.5$\times{10}^{-3}$ | 6.6$\times{10}^{-4}$ | 4.1$\times{10}^{-4}$ | 1.2$\times{10}^{-4}$ |
|  | MACAU | 4.87$\times{10}^{-2}$ | 1.02$\times{10}^{-2}$ | 5.57$\times{10}^{-3}$ | 1.37$\times{10}^{-3}$ | 8.1$\times{10}^{-4}$ | 2.8$\times{10}^{-4}$ | 1.4$\times{10}^{-4}$ | 4$\times{10}^{-5}$ |
|  | GEMMA | 5.12$\times{10}^{-2}$ | 1.02$\times{10}^{-2}$ | 5.15$\times{10}^{-3}$ | 1.1$\times{10}^{-3}$ | 5.2$\times{10}^{-4}$ | 6$\times{10}^{-5}$ | 2$\times{10}^{-5}$ | 1$\times{10}^{-5}$ |
|  | BB | 5.72$\times{10}^{-2}$ | 1.27$\times{10}^{-2}$ | 6.77$\times{10}^{-3}$ | 1.52$\times{10}^{-3}$ | 9.1$\times{10}^{-4}$ | 2.5$\times{10}^{-4}$ | 1.6$\times{10}^{-4}$ | 3$\times{10}^{-5}$ |
| $h^{2}=0.6$ | IMAGE | 5.25$\times{10}^{-2}$ | 1.15$\times{10}^{-2}$ | 6.02$\times{10}^{-3}$ | 1.3$\times{10}^{-3}$ | 8.1$\times{10}^{-4}$ | 1.8$\times{10}^{-4}$ | 1.1$\times{10}^{-4}$ | 5$\times{10}^{-5}$ |
|  | IMAGE-I | 5.44$\times{10}^{-2}$ | 1.21$\times{10}^{-2}$ | 6.36$\times{10}^{-3}$ | 1.34$\times{10}^{-3}$ | 7.5$\times{10}^{-4}$ | 1.9$\times{10}^{-4}$ | 1$\times{10}^{-4}$ | 5$\times{10}^{-5}$ |
|  | IMAGE-A | 5.94$\times{10}^{-2}$ | 1.54$\times{10}^{-2}$ | 8.65$\times{10}^{-3}$ | 2.36$\times{10}^{-3}$ | 1.41$\times{10}^{-3}$ | 4.2$\times{10}^{-4}$ | 2.7$\times{10}^{-4}$ | 7$\times{10}^{-5}$ |
|  | MACAU | 4.83$\times{10}^{-2}$ | 1.02$\times{10}^{-2}$ | 5.04$\times{10}^{-3}$ | 1.03$\times{10}^{-3}$ | 5.9$\times{10}^{-4}$ | 1.1$\times{10}^{-4}$ | 8$\times{10}^{-5}$ | 4$\times{10}^{-5}$ |
|  | GEMMA | 5.17$\times{10}^{-2}$ | 9.9$\times{10}^{-3}$ | 4.97$\times{10}^{-3}$ | 9.7$\times{10}^{-4}$ | 5.1$\times{10}^{-4}$ | 9$\times{10}^{-5}$ | 4$\times{10}^{-5}$ | 1$\times{10}^{-5}$ |
|  | BB | 5.99$\times{10}^{-2}$ | 1.31$\times{10}^{-2}$ | 6.95$\times{10}^{-3}$ | 1.57$\times{10}^{-3}$ | 8.5$\times{10}^{-4}$ | 1.7$\times{10}^{-4}$ | 9$\times{10}^{-5}$ | 4$\times{10}^{-5}$ |

**Supplementary Table 6**: Type I error control of different methods in the null simulations across three different sample sizes. The type I error of different methods (IMAGE, IMAGE-I, IMAGE-A, MACAU, GEMMA, and BB) are shown at significance thresholds ranging from 0.05 to $1\times{10}^{-5}$.

|  | Method | $5\times{10}^{-2}$ | $1\times{10}^{-2}$ | $5\times{10}^{-3}$ | $1\times{10}^{-3}$ | $5\times{10}^{-4}$ | $1\times{10}^{-4}$ | $5\times{10}^{-5}$ | $1\times{10}^{-5}$ |
| --- | --- | --- | --- | --- | --- | --- | --- | --- | --- |
| $n=50$ | IMAGE | 5.58$\times{10}^{-2}$ | 1.28$\times{10}^{-2}$ | 7.10$\times{10}^{-3}$ | 1.$78\times{10}^{-3}$ | 9$.7\times{10}^{-4}$ | 3.$0\times{10}^{-4}$ | $1.3\times{10}^{-4}$ | 6$\times{10}^{-5}$ |
|  | IMAGE-I | 5.89$\times{10}^{-2}$ | 1.46$\times{10}^{-2}$ | 8.50$\times{10}^{-3}$ | $2.83\times{10}^{-3}$ | 1.99$\times{10}^{-3}$ | 1.15$\times{10}^{-3}$ | 9.$7\times{10}^{-4}$ | $8.2\times{10}^{-4}$ |
|  | IMAGE-A | 7.05$\times{10}^{-2}$ | 2.07$\times{10}^{-2}$ | 1.27$\times{10}^{-2}$ | 4.32$\times{10}^{-3}$ | 2.62$\times{10}^{-3}$ | 9.4$\times{10}^{-4}$ | 6.0$\times{10}^{-4}$ | 1.8$\times{10}^{-4}$ |
|  | MACAU | 4.65$\times{10}^{-2}$ | 1.01$\times{10}^{-2}$ | 5.38$\times{10}^{-3}$ | 1.37$\times{10}^{-3}$ | 7.2$\times{10}^{-4}$ | 2.$0\times{10}^{-4}$ | 9$\times{10}^{-5}$ | 2$\times{10}^{-5}$ |
|  | GEMMA | 5.05$\times{10}^{-2}$ | 9.60$\times{10}^{-3}$ | 4.70$\times{10}^{-3}$ | 9.3$\times{10}^{-4}$ | 4.6$\times{10}^{-4}$ | 8$\times{10}^{-5}$ | $1\times{10}^{-5}$ | 0 |
|  | BB | 5.95$\times{10}^{-2}$ | 1.35$\times{10}^{-2}$ | 7.05$\times{10}^{-3}$ | 1.58$\times{10}^{-3}$ | 8.8$\times{10}^{-4}$ | 2.1$\times{10}^{-4}$ | 9$\times{10}^{-5}$ | 1$\times{10}^{-5}$ |
| $n=100$ | IMAGE | 5.38$\times{10}^{-2}$ | 1.16$\times{10}^{-2}$ | 6.44$\times{10}^{-3}$ | 1.62$\times{10}^{-3}$ | 9$\times{10}^{-4}$ | 1.9$\times{10}^{-4}$ | 8$\times{10}^{-5}$ | 0 |
|  | IMAGE-I | 5.45$\times{10}^{-2}$ | 1.21$\times{10}^{-2}$ | 6.86$\times{10}^{-3}$ | 1.67$\times{10}^{-3}$ | 1.05$\times{10}^{-3}$ | 3.5$\times{10}^{-4}$ | 1.6$\times{10}^{-4}$ | 4$\times{10}^{-5}$ |
|  | IMAGE-A | 6.23$\times{10}^{-2}$ | 1.54$\times{10}^{-2}$ | 8.64$\times{10}^{-3}$ | 2.55$\times{10}^{-3}$ | 1.5$\times{10}^{-3}$ | 6.6$\times{10}^{-4}$ | 4.1$\times{10}^{-4}$ | 1.2$\times{10}^{-4}$ |
|  | MACAU | 4.87$\times{10}^{-2}$ | 1.02$\times{10}^{-2}$ | 5.57$\times{10}^{-3}$ | 1.37$\times{10}^{-3}$ | 8.1$\times{10}^{-4}$ | 2.8$\times{10}^{-4}$ | 1.4$\times{10}^{-4}$ | 4$\times{10}^{-5}$ |
|  | GEMMA | 5.12$\times{10}^{-2}$ | 1.02$\times{10}^{-2}$ | 5.15$\times{10}^{-3}$ | 1.1$\times{10}^{-3}$ | 5.2$\times{10}^{-4}$ | 6$\times{10}^{-5}$ | 2$\times{10}^{-5}$ | 1$\times{10}^{-5}$ |
|  | BB | 5.72$\times{10}^{-2}$ | 1.27$\times{10}^{-2}$ | 6.77$\times{10}^{-3}$ | 1.52$\times{10}^{-3}$ | 9.1$\times{10}^{-4}$ | 2.5$\times{10}^{-4}$ | 1.6$\times{10}^{-4}$ | 3$\times{10}^{-5}$ |
| $n=150$ | IMAGE | $5.06\times{10}^{-2}$ | 1.07$\times{10}^{-2}$ | 5.06$\times{10}^{-3}$ | 1.14$\times{10}^{-3}$ | 6.1$\times{10}^{-4}$ | 1.9$\times{10}^{-4}$ | 8$\times{10}^{-5}$ | 4$\times{10}^{-5}$ |
|  | IMAGE-I | 5.20$\times{10}^{-2}$ | 1.32$\times{10}^{-2}$ | 6.03$\times{10}^{-3}$ | 1.39$\times{10}^{-3}$ | 7.5$\times{10}^{-4}$ | 2.1$\times{10}^{-4}$ | 1.4$\times{10}^{-4}$ | 4$\times{10}^{-5}$ |
|  | IMAGE-A | 5.66$\times{10}^{-2}$ | 1.61$\times{10}^{-2}$ | 7.25$\times{10}^{-3}$ | 1.75$\times{10}^{-3}$ | 9.4$\times{10}^{-4}$ | 3.0$\times{10}^{-4}$ | 2$\times{10}^{-4}$ | 5$\times{10}^{-5}$ |
|  | MACAU | 4.77$\times{10}^{-2}$ | 9.47$\times{10}^{-3}$ | 5.29$\times{10}^{-3}$ | 1.17$\times{10}^{-3}$ | 5.8$\times{10}^{-4}$ | 1.8$\times{10}^{-4}$ | 1$\times{10}^{-4}$ | 2$\times{10}^{-5}$ |
|  | GEMMA | 5.03$\times{10}^{-2}$ | 9.66$\times{10}^{-3}$ | 4.58$\times{10}^{-3}$ | 9.2$\times{10}^{-4}$ | 4.6$\times{10}^{-4}$ | 9$\times{10}^{-5}$ | 3$\times{10}^{-5}$ | 0 |
|  | BB | 5.54$\times{10}^{-2}$ | 1.21$\times{10}^{-2}$ | 6.59$\times{10}^{-3}$ | 1.55$\times{10}^{-3}$ | 7.6$\times{10}^{-4}$ | 1$.9\times{10}^{-4}$ | 1.2$\times{10}^{-4}$ | 3$\times{10}^{-5}$ |

**Supplementary Table 7**: Type I error control of different methods in the null simulations across three different minor allele frequency. The type I error of different methods (IMAGE, IMAGE-I, IMAGE-A, MACAU, GEMMA, and BB) are shown at significance thresholds ranging from 0.05 to $1\times{10}^{-5}$.

|  | Method | $5\times{10}^{-2}$ | $1\times{10}^{-2}$ | $5\times{10}^{-3}$ | $1\times{10}^{-3}$ | $5\times{10}^{-4}$ | $1\times{10}^{-4}$ | $5\times{10}^{-5}$ | $1\times{10}^{-5}$ |
| --- | --- | --- | --- | --- | --- | --- | --- | --- | --- |
| $MAF=0.1$ | IMAGE | 5.44$\times{10}^{-2}$ | 1.28$\times{10}^{-2}$ | 7.16$\times{10}^{-3}$ | 1.95$\times{10}^{-3}$ | 1.16$\times{10}^{-3}$ | 3.3$\times{10}^{-4}$ | 2.4$\times{10}^{-4}$ | 5$\times{10}^{-5}$ |
|  | IMAGE-I | 5.31$\times{10}^{-2}$ | 1.15$\times{10}^{-2}$ | 6.04$\times{10}^{-3}$ | 1.44$\times{10}^{-3}$ | 7.9$\times{10}^{-4}$ | 2.$6\times{10}^{-4}$ | 1.6$\times{10}^{-4}$ | 4$\times{10}^{-5}$ |
|  | IMAGE-A | 7.47$\times{10}^{-2}$ | 2.24$\times{10}^{-2}$ | 1.43$\times{10}^{-2}$ | 5.19$\times{10}^{-3}$ | 3.29$\times{10}^{-3}$ | 1.2$\times{10}^{-3}$ | 8.5$\times{10}^{-4}$ | 4$\times{10}^{-4}$ |
|  | MACAU | 4.84$\times{10}^{-2}$ | 1$.03\times{10}^{-2}$ | 5.36$\times{10}^{-3}$ | 1.23$\times{10}^{-3}$ | 6.6$\times{10}^{-4}$ | 2$\times{10}^{-4}$ | 1.3$\times{10}^{-4}$ | 3$\times{10}^{-5}$ |
|  | GEMMA | 5.44$\times{10}^{-2}$ | 1.34$\times{10}^{-2}$ | 7.85$\times{10}^{-3}$ | 2.4$\times{10}^{-3}$ | 1.46$\times{10}^{-3}$ | 4$\times{10}^{-4}$ | 3$\times{10}^{-4}$ | 8$\times{10}^{-5}$ |
|  | BB | 5.45$\times{10}^{-2}$ | 1.14$\times{10}^{-2}$ | 5.86$\times{10}^{-3}$ | 1.29$\times{10}^{-3}$ | 6.8$\times{10}^{-4}$ | 2.2$\times{10}^{-4}$ | 9$\times{10}^{-5}$ | 3$\times{10}^{-5}$ |
| $MAF=0.3$ | IMAGE | 5.38$\times{10}^{-2}$ | 1.16$\times{10}^{-2}$ | 6.44$\times{10}^{-3}$ | 1.62$\times{10}^{-3}$ | 9$\times{10}^{-4}$ | 1.9$\times{10}^{-4}$ | 8$\times{10}^{-5}$ | 0 |
|  | IMAGE-I | 5.45$\times{10}^{-2}$ | 1.21$\times{10}^{-2}$ | 6.86$\times{10}^{-3}$ | 1.67$\times{10}^{-3}$ | 1.05$\times{10}^{-3}$ | 3.5$\times{10}^{-4}$ | 1.6$\times{10}^{-4}$ | 4$\times{10}^{-5}$ |
|  | IMAGE-A | 6.23$\times{10}^{-2}$ | 1.54$\times{10}^{-2}$ | 8.64$\times{10}^{-3}$ | 2.55$\times{10}^{-3}$ | 1.5$\times{10}^{-3}$ | 6.6$\times{10}^{-4}$ | 4.1$\times{10}^{-4}$ | 1.2$\times{10}^{-4}$ |
|  | MACAU | 4.87$\times{10}^{-2}$ | 1.02$\times{10}^{-2}$ | 5.57$\times{10}^{-3}$ | 1.37$\times{10}^{-3}$ | 8.1$\times{10}^{-4}$ | 2.8$\times{10}^{-4}$ | 1.4$\times{10}^{-4}$ | 4$\times{10}^{-5}$ |
|  | GEMMA | 5.12$\times{10}^{-2}$ | 1.02$\times{10}^{-2}$ | 5.15$\times{10}^{-3}$ | 1.1$\times{10}^{-3}$ | 5.2$\times{10}^{-4}$ | 6$\times{10}^{-5}$ | 2$\times{10}^{-5}$ | 1$\times{10}^{-5}$ |
|  | BB | 5.72$\times{10}^{-2}$ | 1.27$\times{10}^{-2}$ | 6.77$\times{10}^{-3}$ | 1.52$\times{10}^{-3}$ | 9.1$\times{10}^{-4}$ | 2.5$\times{10}^{-4}$ | 1.6$\times{10}^{-4}$ | 3$\times{10}^{-5}$ |
| $MAF=0.5$ | IMAGE | 5.05$\times{10}^{-2}$ | 1.08$\times{10}^{-2}$ | 5.7$\times{10}^{-3}$ | 1.1$\times{10}^{-3}$ | 6.4$\times{10}^{-4}$ | 1$.8\times{10}^{-4}$ | 1$\times{10}^{-4}$ | 3$\times{10}^{-5}$ |
|  | IMAGE-I | 5.65$\times{10}^{-2}$ | 1.25$\times{10}^{-2}$ | 6.45$\times{10}^{-3}$ | 1.46$\times{10}^{-3}$ | 8.5$\times{10}^{-4}$ | 1.9$\times{10}^{-4}$ | 1.2$\times{10}^{-4}$ | 5$\times{10}^{-5}$ |
|  | IMAGE-A | 5.82$\times{10}^{-2}$ | 1.34$\times{10}^{-2}$ | 7.6$\times{10}^{-3}$ | 1.91$\times{10}^{-3}$ | 9.4$\times{10}^{-4}$ | 2.8$\times{10}^{-4}$ | 1.9$\times{10}^{-4}$ | 5$\times{10}^{-5}$ |
|  | MACAU | 4.8$\times{10}^{-2}$ | 9.87$\times{10}^{-3}$ | 4.7$\times{10}^{-3}$ | 1.08$\times{10}^{-3}$ | 5.9$\times{10}^{-4}$ | 1.1$\times{10}^{-4}$ | 5$\times{10}^{-5}$ | 1$\times{10}^{-5}$ |
|  | GEMMA | 4.95$\times{10}^{-2}$ | 8.82$\times{10}^{-3}$ | 4.08$\times{10}^{-3}$ | 6.6$\times{10}^{-4}$ | 3.9$\times{10}^{-4}$ | 8$\times{10}^{-5}$ | 4$\times{10}^{-5}$ | 0 |
|  | BB | 6.23$\times{10}^{-2}$ | 1.42$\times{10}^{-2}$ | 7.33$\times{10}^{-3}$ | 1.74$\times{10}^{-3}$ | 9.3$\times{10}^{-4}$ | 1.8$\times{10}^{-4}$ | 1$\times{10}^{-4}$ | 4$\times{10}^{-5}$ |

**REFERENCE**

1.	Breslow, N.E. and D.G. Clayton, *Approximate Inference in Generalized Linear Mixed Models.* Journal of the American Statistical Association, 1993. **88**(421): p. 9-25.

2.	Chen, H., et al., *Control for Population Structure and Relatedness for Binary Traits in Genetic Association Studies via Logistic Mixed Models.* American Journal of Human Genetics, 2016. **98**(4): p. 653-666.

3.	Gilmour, A.R., R. Thompson, and B.R. Cullis, *Average information REML: An efficient algorithm for variance parameter estimation in linear mixed models.* Biometrics, 1995. **51**(4): p. 1440-1450.

4.	Yang, J.A., et al., *GCTA: A Tool for Genome-wide Complex Trait Analysis.* American Journal of Human Genetics, 2011. **88**(1): p. 76-82.

5.	Wall, J.D., et al., *Genomewide ancestry and divergence patterns from low-coverage sequencing data reveal a complex history of admixture in wild baboons.* Molecular Ecology, 2016. **25**(14): p. 3469-3483.

6.	Lea, A.J., et al., *Resource base influences genome-wide DNA methylation levels in wild baboons (Papio cynocephalus).* Molecular Ecology, 2016. **25**(8): p. 1681-1696.
